# A synthetic C2 auxotroph of *Pseudomonas putida* for evolutionary engineering of alternative sugar catabolic routes

**DOI:** 10.1101/2022.07.21.500976

**Authors:** Nicolas T. Wirth, Nicolás Gurdo, Nicolas Krink, Àngela Vidal Verdú, Lorena Férnandez-Cabezón, Tune Wulff, Pablo I. Nikel

## Abstract

Acetyl-coenzyme A (AcCoA) is a metabolic hub in virtually all living cells, serving as both a key precursor of essential biomass components and a metabolic sink for catabolic pathways of a large variety of substrates. Owing to this dual role, tight growth-production coupling schemes can be implemented around the AcCoA node. Inspired by this concept, a synthetic C2 auxotrophy was implemented in the platform bacterium *Pseudomonas putida* through an *in silico*-guided engineering approach. A growth-coupling strategy, driven by AcCoA demand, allowed for direct selection of an alternative sugar assimilation route—the phosphoketolase (PKT) shunt from bifidobacteria. Adaptive laboratory evolution forced the synthetic auxotroph to integrate the PKT shunt to restore C2 prototrophy. Large-scale structural chromosome rearrangements were identified as possible mechanisms for adjusting the network-wide proteome profile, resulting in improved PKT-dependent growth phenotypes. ^13^C-based metabolic flux analysis revealed an even split between the native Entner-Doudoroff and the synthetic PKT pathway for glucose processing, leading to enhanced carbon conservation. These results demonstrate that the *P. putida* metabolism can be radically rewired to incorporate a synthetic C2 metabolism, creating novel network connectivities and highlighting the importance of unconventional engineering strategies to support efficient microbial production.

## Introduction

Acetyl-coenzyme A (AcCoA) is a key component in the metabolism of virtually all living organisms [1,2]. The thioester is a metabolite hub and the entry point to the tricarboxylic acid (TCA) cycle, in which most of the energy-generating reducing equivalents are produced. AcCoA also plays a central role in biochemical pathways leading to essential biomass constituents, e.g., amino acids, cell wall components (acetylated amino sugars), fatty acids [3], and multiple compounds of industrial interest (i.e., polyketides, isoprenoids, alcohols, and polyhydroxyalkanoates) can be produced from AcCoA [4–6]. Conversely, various catabolic pathways for a broad spectrum of compounds form a ‘funnel’ converging at the AcCoA node [7]. Given its central role as a biomass precursor, the pivotal AcCoA node can be harnessed for constructing synthetic pathways where both product formation and consumption are coupled to growth— provided that they represent the only AcCoA source in a suitable selection strain.

Having adapted to frequent periods of carbon starvation, the metabolism of soil generalist microbes (e.g., *Pseudomonas putida*) is optimized for the utilization of a wide variety of substrates without producing overflow metabolites [8–11]. The intracellular AcCoA concentration—and its relative levels compared to nonesterified CoA—is maintained at low values in *P. putida* regardless of the metabolic status [12], in contrast to facultatively anaerobic bacteria [13,14]. In the two most prevalent glycolytic routes in bacteria, the Embden-Meyerhof-Parnas (EMP) and the Entner-Doudoroff (ED) pathway, AcCoA is almost exclusively formed through pyruvate oxidation [15–17]. This reaction, catalyzed by the pyruvate dehydrogenase complex (PDHc), involves substrate decarboxylation and consequently leads to net carbon loss [18]. In contrast, the phosphoketolase (PKT) pathway [19], originally described in heterolactic fermentative bacteria, is an alternative glycolytic route that provides AcCoA without CO_2_ release. PKTs form a unique class of enzymes, closely related to transketolase (EC 2.2.1.1), an essential component of the pentose phosphate (PP) pathway [20]. They require thiamine diphosphate and divalent metal ions as cofactors, catalyzing the cleavage of xylulose 5-phosphate (Xu5P) or fructose 6-phosphate (F6P) to acetyl-phosphate (Ac-P) and glyceraldehyde 3-phosphate (G3P) or erythrose 4-phosphate (E4P), respectively. The enzymatic reaction involves ketol cleavage, dehydration, and phosphorolysis in a single step [21]. Ac-P can be converted into AcCoA *via* phosphate acetyltransferase (Pta), thereby allowing direct conversion of sugar phosphates into AcCoA without carbon loss [22]. Streamlining sugar utilization through the use of this carbon-conserving pathway has the potential of increasing maximum theoretical yields for AcCoA-derived products in biotechnological applications [23–25]. Such a strategy has been harnessed both in bacteria and yeast, most often as an additional route to the linear EMP glycolysis [2,26,27]. The question is whether a synthetic PKT metabolism can be implanted in *P. putida* to boost AcCoA levels—but such a goal entails a radical rewiring of the metabolic connections that lead to AcCoA formation in this obligate aerobe [28] that runs a cyclic, ED-based sugar catabolism [29,30].

The inherent complexity of metabolic systems poses an obstacle when deliberately modifying core traits, yet adaptive laboratory evolution (ALE) aids in improving the performance of whole-cell biocatalysts for specific biochemical tasks [31,32]. A requisite for evolutionary engineering approaches is that the desired property can be subject to a selection pressure that couples an improved catalytic performance with a fitness gain—e.g., by creating a metabolic deficiency in selection strains [33,34]. Building on these concepts, this study adopted *in silico* modeling to identify key reactions involved in AcCoA turnover in *P. putida*, and a synthetic two-carbon (C2) auxotrophy was established by systematically eliminating all of the AcCoA-forming reactions. As proof-of-principle of C2-based selection schemes, an artificial PKT pathway was implemented in the rewired strain, and evolutionary engineering was applied to optimize the synthetic metabolism. Multi-omic characterization, including functional genomics, targeted proteomics, and ^13^C-based fluxomics, showed that network-wide adaptations enabled carbon conservation in the evolved clones—as evidenced by a ca. 80% reduction in CO_2_ formation in bioreactor batch cultures. Taken together, our results demonstrate the suitability of establishing alternative routes for sugar catabolism in *P. putida*, supported by *in silico*-predicted metabolic rewiring and evolutionary engineering, thus paving the way towards carbon-efficient bioproduction.

## Materials and Methods

### Bacterial strains and culture conditions

The bacterial strains employed in this study are listed in **Table 1**. *E. coli* and *P. putida* were incubated at 37°C and 30°C, respectively. For propagation and storage, routine cloning procedures, and during genome engineering manipulations, cells were grown in lysogeny broth (LB) medium (10 g L^−1^ tryptone, 5 g L^−1^ yeast extract, and 10 g L^−1^ NaCl). Liquid cultures were performed using either 50-mL centrifuge tubes with a medium volume of 5-10 mL, or in 500-mL Erlenmeyer flasks covered with cellulose plugs (Carl Roth, Karlsruhe, Germany) with a medium volume of 50 mL. All liquid cultures were agitated at 250 rpm (MaxQ™8000 incubator; ThermoFisher Scientific, Waltham, MA, USA). Solid culture media contained 15 g L^−1^ agar. Selection of plasmid-harboring cells was achieved by adding kanamycin (Km), gentamicin (Gm), or streptomycin (Sm) to the media at 50 µg mL^−1^, 10 µg mL^−1^, and 50 µg mL^−1^, respectively.

**Table 1.**
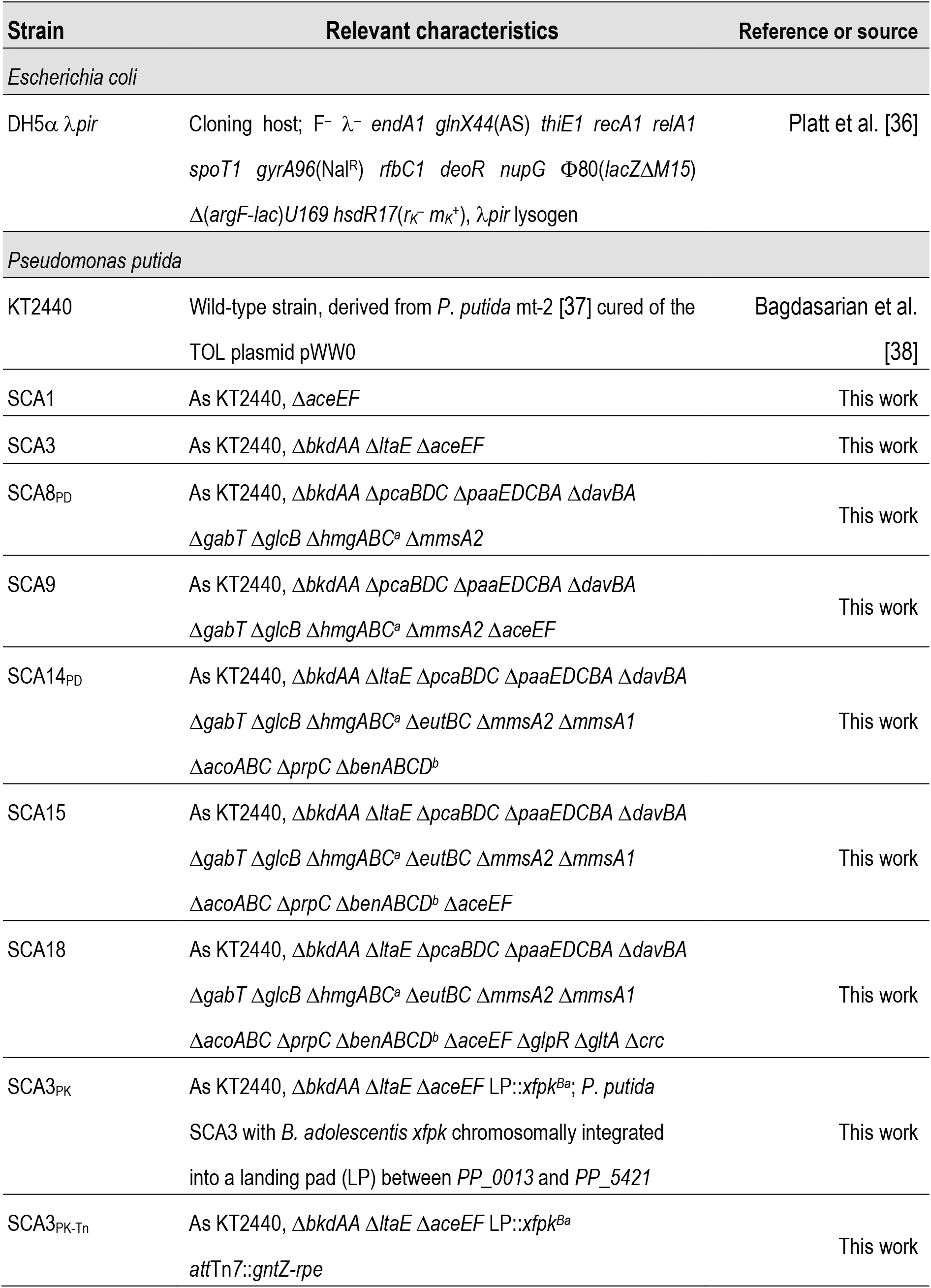
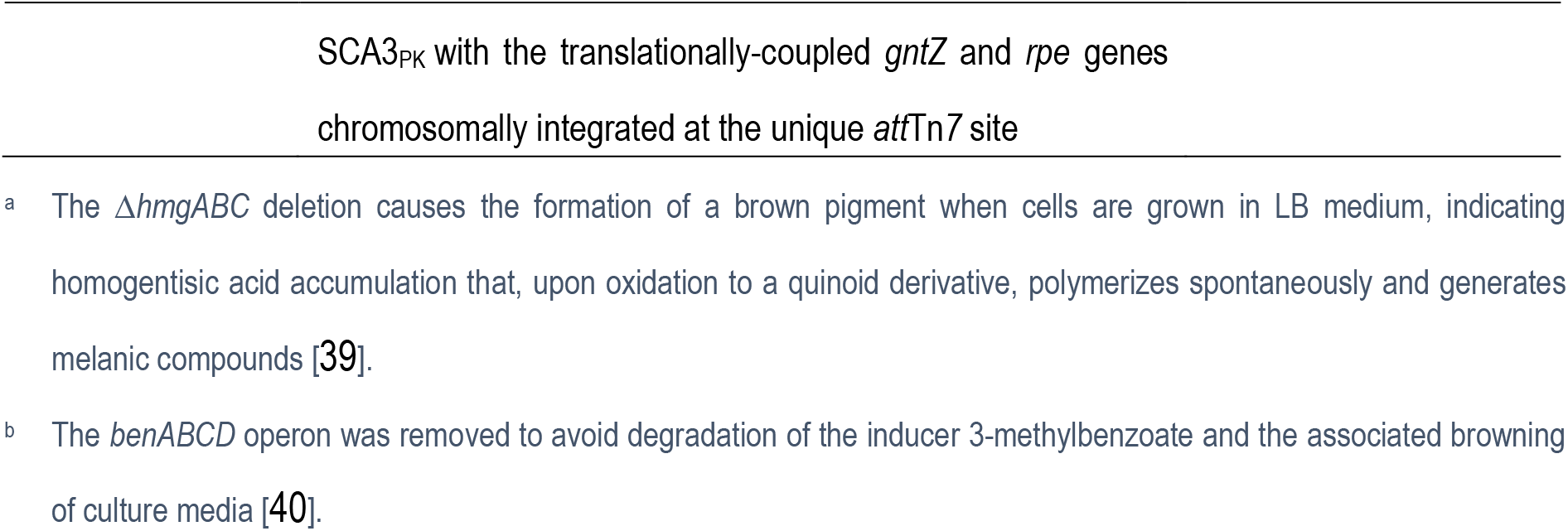
Bacterial strains used in this study.

| Strain | Relevant characteristics | Reference or source |
| --- | --- | --- |
| *Escherichia coli* |  |  |
| DH5α λpir | Cloning host; F<sup>-</sup> λ- *endA1 glnX44(AS) thiE1 recA1 relA1 spoT1 gyrA96(Nal<sup>R</sup>) rfbC1 deoR nupG* Φ80(*lacZ*ΔM15) $\Delta$(*argF-lac*)U169 *hsdR17(r<sub>K</sub><sup>-</sup> m<sub>K</sub><sup>+</sup>)*, λpir lysogen | Platt et al. [36] |
| *Pseudomonas putida* |  |  |
| KT2440 | Wild-type strain, derived from *P. putida* mt-2 [37] cured of the TOL plasmid pWW0 | Bagdasarian et al. [38] |
| SCA1 | As KT2440, ΔaceEF | This work |
| SCA3 | As KT2440, ΔbkdAA ΔltaE ΔaceEF | This work |
| SCA8<sub>PD</sub> | As KT2440, ΔbkdAA ΔpcaBDC ΔpaaEDCBA ΔdavBA ΔgabT ΔglcB ΔhmgABC<sup>a</sup> ΔmmsA2 | This work |
| SCA9 | As KT2440, ΔbkdAA ΔpcaBDC ΔpaaEDCBA ΔdavBA ΔgabT ΔglcB ΔhmgABC<sup>a</sup> ΔmmsA2 ΔaceEF | This work |
| SCA14<sub>PD</sub> | As KT2440, ΔbkdAA ΔltaE ΔpcaBDC ΔpaaEDCBA ΔdavBA ΔgabT ΔglcB ΔhmgABC<sup>a</sup> ΔeutBC ΔmmsA2 ΔmmsA1 ΔacoABC ΔprpC ΔbenABCD<sup>b</sup> | This work |
| SCA15 | As KT2440, ΔbkdAA ΔltaE ΔpcaBDC ΔpaaEDCBA ΔdavBA ΔgabT ΔglcB ΔhmgABC<sup>a</sup> ΔeutBC ΔmmsA2 ΔmmsA1 ΔacoABC ΔprpC ΔbenABCD<sup>b</sup> ΔaceEF | This work |
| SCA18 | As KT2440, ΔbkdAA ΔltaE ΔpcaBDC ΔpaaEDCBA ΔdavBA ΔgabT ΔglcB ΔhmgABC<sup>a</sup> ΔeutBC ΔmmsA2 ΔmmsA1 ΔacoABC ΔprpC ΔbenABCD<sup>b</sup> ΔaceEF ΔglpR ΔgltA Δcrc | This work |
| SCA3<sub>PK</sub> | As KT2440, ΔbkdAA ΔltaE ΔaceEF LP::*xfpk*<sup>Ba</sup>; *P. putida* SCA3 with *B. adolescentis* *xfpk* chromosomally integrated into a landing pad (LP) between PP_0013 and PP_5421 | This work |
| SCA3<sub>PK-Tn</sub> | As KT2440, ΔbkdAA ΔltaE ΔaceEF LP::*xfpk*<sup>Ba</sup> *attTn7::gntZ-rpe* | This work |

- <sup>a</sup> The $\Delta hmgABC$ deletion causes the formation of a brown pigment when cells are grown in LB medium, indicating homogentisic acid accumulation that, upon oxidation to a quinoid derivative, polymerizes spontaneously and generates melanic compounds [39]. - <sup>b</sup> The *benABCD* operon was removed to avoid degradation of the inducer 3-methylbenzoate and the associated browning of culture media [40].

For phenotypic characterizations in microtiter plates as well as in shaken-flask cultivations, experiments were performed in de Bont minimal (DBM) medium [35] additionally buffered with 5 g L^−1^ 3-(*N*-morpholino)propane sulfonic acid (MOPS) at pH = 7.0 and supplemented with different carbon compounds as explained in the text. Before launching plate reader (for 96-well plates: ELx808, BioTek Instruments; Winooski, VT, USA; for 24-well plates: Synergy H1, BioTek Instruments) experiments with SCA strains and varying carbon sources, cells were pre-grown in LB medium. For all other experiments, the pre-culture media were identical to those used for the experiment unless otherwise indicated. The precultures were harvested by centrifugation at 8,000×*g* for 2 min, washed with DBM medium without any carbon substrate, and resuspended in the final media at the desired starting optical density measured at 600 nm (OD_600_). Cell growth was monitored by measuring the OD_600_. ALE of strains was performed in 24-well deep-well plates covered with a Sandwich Cover (PreSens Precision Sensing, Regensburg, Germany) and filled with 2 mL of DBM medium (containing an appropriate substrate) per well; other ALE conditions are described in the text.

### Cloning procedures and plasmid construction

All plasmids used in this work are listed in **Table 2**. Oligonucleotides used for the PCR-amplification of fragments and genotyping via colony PCR are listed in **Table S1** in Additional File 1. Uracil-excision (*USER*) cloning [41–43] was used for constructing all plasmids. The AMUSER tool was employed for oligonucleotide design [44]. All genetic manipulations followed protocols described previously [40,45]. DNA fragments employed in assembly reactions were amplified using the Phusion™ *U* high-fidelity DNA polymerase (Thermo Fisher Scientific, Waltham, Massachusetts, USA) according to the manufacturer’s specifications. The identity and correctness of all plasmids and DNA constructs were confirmed by Sanger sequencing (Eurofins Genomics, Ebersberg, Germany). For genotyping experiments after cloning procedures and genome manipulations, colony PCRs were performed using the commercial *OneTaq*™ master mix (New England BioLabs; Ipswich, MA, USA) according to the manufacturer’s instructions. *E. coli* DH5α λ*pir* was employed for cloning purposes. Chemically-competent *E. coli* cells were prepared and transformed with plasmids according to well-established protocols [46–51]. *P. putida* was rendered electro-competent by treatment Choi et al. [52].

**Table 2.** Plasmids used in this study.

| Plasmid name | Relevant characteristics <sup>a</sup> | Source or reference |
| --- | --- | --- |
| pGNW2 | Suicide vector used for genetic manipulations in Gram-negative bacteria; *oriT*, *traJ*, *lacZα*, *ori*(R6K), *P*<sub>14g</sub>(*BCD2*)→*msfGFP*; Km<sup>R</sup> | Wirth *et al.* [45] |
| pSNW2 | Derivative of vector pGNW2 with the translation initiation sequence of *msfGFP* replaced by the very strong translational coupling sequence *BCD2* | Volke *et al.* [40] |
| pQURE6-H | Conditionally-replicating vector; derivative of vector pJBSD1 carrying *XylS/Pm*→*I-SceI* and *P*<sub>14g</sub>(*BCD2*)→*mRFP*; Gm<sup>R</sup> | Volke *et al.* [40] |
| pSNW2·Δ*aceEF* | Derivative of pGNW2 carrying HAs to delete *aceEF* in *P. putida* KT2440 | This study |
| pSNW2·Δ*gltA* | Derivative of pGNW2 carrying HAs to delete *gltA* in *P. putida* KT2440 | This study |
| pSNW2·Δ*acoABC* | Derivative of pGNW2 carrying HAs to delete *acoABC* in *P. putida* KT2440 | This study |
| pSNW2·Δ*amaC-dpkA* | Derivative of pGNW2 carrying HAs to delete *amaC-dpkA* in *P. putida* KT2440 | This study |
| pSNW2·Δ*bkdAA* | Derivative of pGNW2 carrying HAs to delete *bkdAA* in *P. putida* KT2440 | This study |
| pSNW2·Δ*crc* | Derivative of pGNW2 carrying HAs to delete *crc* in *P. putida* KT2440 | This study |
| pSNW2·Δ*davBA* | Derivative of pGNW2 carrying HAs to delete *davBA* in *P. putida* KT2440 | This study |
| pSNW2·Δ*eutBC* | Derivative of pGNW2 carrying HAs to delete *eutBC* in *P. putida* KT2440 | This study |
| pSNW2·Δ*gabT* | Derivative of pGNW2 carrying HAs to delete *gabT* in *P. putida* KT2440 | This study |
| pSNW2·$\Delta glcB$ | Derivative of pGNW2 carrying HAs to delete *glcB* in *P. putida* KT2440 | This study |
| pSNW2·$\Delta hmgABC$ | Derivative of pGNW2 carrying HAs to delete *hmgABC* in *P. putida* KT2440 | This study |
| pSNW2·$\Delta ltaE$ | Derivative of pGNW2 carrying HAs to delete *ltaE* in *P. putida* KT2440 | This study |
| pSNW2·$\Delta mmsA1$ | Derivative of pGNW2 carrying HAs to delete *mmsA1* in *P. putida* KT2440 | This study |
| pSNW2·$\Delta mmsA2$ | Derivative of pGNW2 carrying HAs to delete *mmsA2* in *P. putida* KT2440 | This study |
| pSNW2·$\Delta paaEDCBA$ | Derivative of pGNW2 carrying HAs to delete *paaEDCBA* in *P. putida* KT2440 | This study |
| pSNW2·$\Delta pcaBDC$ | Derivative of pGNW2 carrying HAs to delete *pcaBDC* in *P. putida* KT2440 | This study |
| pSNW2·$\Delta prpC$ | Derivative of pGNW2 carrying HAs to delete *prpC* in *P. putida* KT2440 | This study |
| pTn7-M | Tn7 integration vector; *oriV*(R6K); Km<sup>R</sup>, Gm<sup>R</sup> | Choi et al. [53] |
| pTn7·P<sub>14g</sub>(BCD10)→*gntZ-rpe* | Derivative of pTn7-M used for chromosomal integration of a synthetic *gntZ-rpe* operon under the control of a P<sub>14g</sub>(BCD10) element. The two genes within the synthetic operon were translationally coupled by overlapping the *STOP* codon (in bold) of *gntZ* and the *START* codon of *rpe* (underlined), i.e., <u>ATGA</u> | This study |
| pTNS2 | Helper plasmid constitutively expressing the *tnsABCD* genes encoding the Tn7 transposase; *oriV*(R6K); Amp<sup>R</sup> | Choi and Schweizer [54] |
| pS438·*pta*<sup>Ec</sup> | Expression vector; *oriV*(pBBR1), Sm<sup>R</sup>; XylS/Pm→*pta*<sup>Ec</sup> | This work |

1 <sup>a</sup> Antibiotic markers: *Amp*, ampicillin; *Km*, kanamycin; *Gm*, gentamicin; and *Sm*, streptomycin. *HA*, homology arms.

### Genome-scale metabolic model (GSMM) simulations

The most recent genome-scale metabolic network reconstruction for *P. putida* KT2440, iJN1463 [55], was subjected to flux balance analysis [FBA [56]] within the COBRApy toolbox [57] to predict optimal flux distributions [58–60] and to identify reactions contributing to AcCoA turnover. Glucose uptake rate and ATP maintenance requirement were fixed at 6.3 mmol g cell dry weight (CDW)^−1^ h^−1^ and at 0.92 mmol gCDW^−1^ h^−1^ [experimentally determined by del Castillo et al. [61] and Ebert *et al*. [9], respectively]. After an initial deletion of the PDHc reaction, the following steps were iterated to identify potential alternative AcCoA sources (reaction loops were not permitted): (i) optimizing flux distribution for biomass formation; (ii) identifying AcCoA-forming reaction with the highest flux under the current distribution; (iii) tracing fluxes leading to AcCoA to pinpoint the metabolic pathway(s) involved therein; and (iv) removing the entry reaction of the pathway, simultaneously avoiding the creation of auxotrophies. The identified pathways were then analyzed regarding the flux towards the first biomass constituent in the AcCoA-generating pathway and ranked according to their value within the flux distribution of the wild-type model. This ranking was used to inform a hierarchy of gene deletions to be established in *P. putida* to create a family of synthetic C2 auxotrophic strains.

### Targeted proteomics assisted by mass spectrometry

*P. putida* strains were pre-cultured overnight in DBM medium supplemented with 30 mM glucose or 30 mM glucose and 30 mM acetate; cultures with identical media were inoculated at an initial OD_600_ of 0.05. Samples were taken in the mid-exponential phase, and cells were harvested by centrifugation at 17,000×*g* for 2 min at 4°C. After removal of the supernatant, cell pellets were frozen and kept at −80°C until proteomics analysis was performed as described previously [62]. To assign the detected peptides to their functions, a protein database consisting of the *P. putida* reference proteome [UP000000556 [10]] was used, supplemented with heterologously expressed proteins.

### Statistical analysis of proteomics data

Proteomics data, acquired as described above, were analyzed using a customized R script in RStudio (version 2021.09.2). First, entries with missing values were filtered to retain only those proteins that had been detected in every replicate of at least one condition. The abundance values were log_2_-transformed and normalized *via* variance stabilization normalization using the *vsn* package [63]. Next, missing values were imputed via random draws from a left-shifted distribution [“MinProb” method in the *MSnbase* package with a q value of 0.01 [64]]. A differential enrichment test was performed for each contrast in the dataset based on protein-wise linear models and empirical Bayes statistics using *limma* [65]. *P*-values were adjusted using *fdrtool* with the Benjamini-Hochberg method [66]. The adjusted *P*-value threshold for significant observations was set at a value of 0.05. Volcano plots were generated with the *ggplot2* package. The *pheatmap* package was used to create heatmaps of protein abundances within all samples. Enrichment analysis of KEGG pathways [67] was performed using the *clusterProfiler* package [68], with all genes listed for *P. putida* KT2440 as background ‘universe’ and a *P*-value threshold of 0.1.

### Adaptive laboratory evolution

Evolution was performed in 24-well deep-well plates filled with 2 mL of DBM medium supplemented with 30 mM glucose and 0.5 mM 3-*m*Bz. A volume of 50 µL (after glucose cultures) or 20 µL (after culture steps with glucose and acetate) of each culture was passed into fresh medium every other day. For every second passage, the medium contained an additional 20 mM of potassium acetate to recover biomass at higher concentrations. After 1 month of continued cultivation, ALE was continued using DBM supplemented with only glucose.

### Bioreactor fermentations

The batch fermentations were performed in BIOSTAT™ Qplus 1-L bioreactors (Sartorius, Göttingen, Germany) with a working culture volume of 800 mL. Precultures were grown overnight in 500-mL Erlenmeyer flasks filled with 10% (v/v) DBM medium supplemented with 5 g L^−1^ MOPS and 30 mM glucose. The same medium was used in the bioreactors. The precultures were harvested by centrifugation at 4,000×*g* for 10 min and resuspended in culture medium to inoculate the bioreactors. Aeration of the vessels was achieved via horseshoe*-*shaped spargers with ambient air at a flow rate of 0.824 L min^−1^. The dissolved oxygen concentration (dO_2_) was measured online using an oxygen electrode and kept above 40% of saturation by increasing the stirring speed from 500 to 1,000 rpm. The lower stirring speed threshold was reduced to 250 rpm after 5.5 h to reduce the shear stress on the cells and to prevent biofilm formation. Sigma-Aldrich Antifoam 204 reagent (MilliporeSigma, St. Louis, Missouri, USA) was added to reduce foaming by adding single droplets when required (only for strain KT2440). The cultures were maintained at 30°C and pH = 7.0 by online measurements with standard thermometers and pH electrodes and the addition of 7 M NaOH. The composition of the off-gas was analyzed online with a Prima BT Bench Top Process mass spectrometer (Thermo Fisher Scientific).

Physiological parameters for *P. putida* KT2440 and SCA3_PK-Tn_/pS438·*pta*^*Ec*^ were obtained as follows: cell growth was followed via OD_600_ measurements and, for strain KT2440, CDW determinations were carried out with a Moisture analyzer WBA-110M (Witeg Labortechnik, Wertheim, Germany). CDW values were derived from OD_600_ measurements by applying correlation factors obtained from standard curves with the absorbance and CDW determinations for each strain. Specific growth rates (µ) were determined as described below. Biomass yields on glucose *Y*_X/S_ (or the sum of all six-carbon moieties) were obtained as the slope of linear correlations of biomass (g_CDW_ L^−1^)-over-substrate (g_S_ L^−1^) plots. The CO_2_ production was calculated from the different CO_2_ contents of the in-gas and the off-gas. Dissolved inorganic carbon species (carbonate and dissolved CO_2_) were neglected. The C-mol amount in biomass was derived from the CDW and the elemental biomass composition of *P. putida* KT2440 [69].

### ^13^C-labeling experiments

For parallel isotopic tracer experiments, *P. putida* KT2440 and strain SCA3_PK-Tn_/pS438·*pta*^*Ec*^ were grown as indicated in section *2*.*1* by supplementing the medium with the corresponding labeled glucose. Three different isotopic tracers were used: (i) 99% [3–^13^C_1_]-glucose, (ii) 99% [4–^13^C_1_]-glucose, and (iii) a 50%:50% mixture of naturally labeled ^12^C and 99% [^13^C_6_]-glucose. All labeled substrates were purchased from Cambridge Isotope Laboratories Inc. (Teddington, Middlesex, United Kingdom). Before tracer experiments, cells were streaked from a cryo stock onto an LB agar plate and grown overnight for 18 h.

Pre-cultures were performed in 12-mL plastic tubes containing 3 mL of DBM medium (with the respective labeled carbon sources) by picking one colony per biological replicate from the plate. The pre-cultures were incubated at 250 rpm in an orbital shaker at 30°C. The cultures were inoculated at an initial OD_600_ of 0.05 in a 100-mL Erlenmeyer flask containing 20 mL of DBM medium with the specific tracer. Aliquots corresponding to the respective biomass were taken from the pre-cultures, centrifuged at 10,000×*g* and 4°C for 5 min, and washed twice using DBM medium without a carbon source. All experiments were performed in two biological replicates and two technical replicates. GC-MS labeling analysis of amino acids and sugars was performed according to Kohlstedt and Wittmann [70].

### Reaction network and computational design for flux estimation

The metabolic networks of *P. putida* KT2440 and strain SCA3_PK-Tn_/pS438·*pta*^*Ec*^ were built based on the most updated genome-scale metabolic model (Nogales et al., 2020). In total, 84 reactions were included as part of the central carbon metabolism in *P. putida* KT2440. In the case of strain SCA3_PK-Tn_/pS438·*pta*^*Ec*^, three additional reactions were added: (i) F6P to Ac-P and E4P, (ii) Xu5P to Ac-P and G3P [(i) and (ii) are the reactions catalyzed PKT], and (iii) Ac-P to AcCoA. All included reactions as well as the carbon atom transitions are listed in Additional File 2. The INCA software package was utilized to analyze the metabolic network for ^13^C-metabolic flux analysis (^13^C-MFA) [71]. Specific growth rates, glucose, gluconate, 2-ketogluconate, and pyruvate uptake or secretion rates were used to constrain the MFA model. The biomass equation was derived from the normalized precursor drainage to calculate experimental growth rates [70]. The relative intracellular fluxes (expressed as a percentage of the *q*_S_) were calculated by minimizing the sum-of-squared residuals between computationally simulated and experimentally determined measurements (MDVs or mass distribution vectors for each fragment analyzed). The flux distributions of *P. putida* KT2440 and strain SCA3_PK-Tn_/pS438·*pta*^*Ec*^ were visualized by mapping the computed flux values onto custom metabolic maps using the R package *fluctuator*. Thereby, the Glcnt and 2KG uptake rates were combined into a single reaction, since their relative contribution could not be determined based on the carbon transitions.

### Metabolite analyses via HPLC

Sugars and organic acids were analyzed using a Dionex Ultimate 3000 HPLC with an Aminex™ HPX-87X Ion Exclusion (300×7.8 mm) column (BioRad, Hercules, CA) as well as RI-150 refractive index and UV (260, 277, 304 and 210 nm) detectors. For analysis, the column was maintained at 65°C and a 5 mM H_2_SO_4_ solution was used as the mobile phase at a flow rate of 0.5 mL min^−1^. HPLC data were processed using the Chromeleon 7.1.3 software (Thermo Fisher Scientific), and compound concentrations were calculated from peak areas using calibration curves with five different standard concentrations.

### Determination of in vitro phosphoketolase activity

PKT activity was determined in cell lysates according to the hydroxamate method adapted from Lipmann and Tuttle [72]. This assay relies on the reaction of enzymatically-produced Ac-P with hydroxylamine to form hydroxamate and inorganic phosphate. Hydroxamate is converted into a ferric hydroxamate complex via the addition of FeCl_3_ (**Figure 3**). The complex exhibits an orange-brown to purplish-brown color, depending on the Ac-P concentration, and can be quantified spectrophotometrically between 480 and 540 nm. FeCl_3_ shows no absorption at these wavelengths. For these determinations, *P. putida* strains were cultured overnight in 50 mL of LB medium supplemented with 0.5 mM 3-*m*Bz to induce *xfpk* expression. All following steps were performed on ice and with pre-chilled solutions. The cells were collected by centrifugation at 5,000×*g* for 10 min at 4°C, washed twice with phosphate buffer (0.05 M phosphate buffer, 0.5 g L^−1^ L-cysteine, and 1 mM MgCl_2_; pH = 6.5), and suspended in 3 mL of the same buffer. The suspensions were split into 1-mL aliquots in 2-mL cryotubes, and 0.3 g of acid-washed glass beads were added. The cells were lysed using a bead-beater homogenizer (BioSpec Products, Bartlesville, USA) at 6,000 rpm (two cycles, 20 s). Clear lysates were obtained by centrifugation at 17,000×*g* for 2 min at 4°C. The protein concentrations were determined via the Bradford assay [73] and all lysates were adjusted to the lowest concentration measured by adding phosphate buffer. The enzyme assay was performed in 1.5-mL micro reaction tubes. Therefore, 600 µL of lysate was added to 150 µL of halogen solution (6 mg mL^−1^ NaF and 10 mg mL^−1^ iodoacetic acid) and 150 µL of 80 mg mL^−1^ D-fructose-6-phosphate. The reaction mixtures were incubated at 30°C, and 75-µL samples were taken in 10-min intervals. The samples were added into wells of a 96-well plate pre-filled with 75 µL of 2 M hydroxylamine (freshly adjusted to pH = 7 with 1 M HCl) and incubated for 10 min at room temperature. Subsequently, 50 µL of 150 mg mL^−1^ trichloroacetic acid, 50 µL of 4 M HCl, and 50 µL of 50 g L^−1^ FeCl_3_ in 0.1 M HCl were added to the mixture. After the last tube was processed, the absorption at 505 nm was measured for all samples in a Synergy H1 plate reader. A calibration curve with Ac-P (lithium) salt was created by subjecting solutions of different concentrations to the same assay procedure.

### Whole-genome sequencing

The sequencing of chromosomal (and plasmid) DNA to analyze the effects of evolution was performed by Novogene (Cambridge, UK). The DNA content of strains was purified using the PureLink™ Genomic DNA Mini Kit (Invitrogen, CA, USA). The raw data was processed in Geneious Prime 2021.1.1 (Biomatters Ltd.) by read pairing, trimming (*BBDuk* plugin with settings [Trim adapters], [Trim low quality], [Minimum Quality: 20], [Trim Adapters based on paired read overhangs with a minimum overlap of 20], [Discard Short Reads, with minimum length 20]), and mapping to the reference genome sequence (with [Mapper: Geneious], [Medium Sensitivity], [Find structural variants, short insertions, and deletions of any size], and [Minimum support for structural variant discovery: 10 reads]). Sequence polymorphisms were identified by using the build-in *Find Variations/SNPs* function with default settings [31,74]. Single-nucleotide polymorphisms (SNPs) found in both pre-evolved and evolved clones were discarded.

### Data and statistical analyses

All experiments were performed with at least three biological replicates unless otherwise stated, and the mean values ± standard deviation are presented. Statistical significances were determined by calculating p-values with the T.Test function in Microsoft Excel 2016. Maximum exponential growth rates (µ_*max*_) were determined by Gaussian process regression using the Python-based tool deODorizer [75]. Average specific growth rates over the exponential phase (µ) were determined by fitting Ln-transformed OD_600_ data versus time to simple linear regressions. Biomass-specific substrate consumption rates (*q*_S_) and metabolite secretion rates (*q*_P_) were calculated by applying polynomial fits to concentration-vs.-time plots, calculating the derivatives of the functions at each sampling time point, and dividing the derivatives by the CDW concentration at the respective time point. The results are given as the mean *q* values of all sampling time points within the log growth phase ± standard deviation. All linear regressions, as well as data visualizations, were performed in OriginPro 2021 (OriginLab Corporation). Figures and Illustrations were created in OriginPro 2021 and Adobe Illustrator 2020. Geneious Prime 2021.1.1 (Biomatters Ltd.) served as a database for any kind of DNA sequences, design plasmids and constructs, and analyze Sanger and NGS sequencing results.

## Results

### Design of a synthetic C2-auxotroph strain informed by genome-scale metabolic modeling and in silico analysis of a phosphoketolase-based synthetic metabolism

To generate a robust C2 auxotroph platform strain on glucose for evolutionary engineering applications, we employed an *in silico*-guided approach to identify native pathways involved in AcCoA supply (39 reactions) and consumption (19 reactions; **Figure 1A**). Next, we performed FBA using the genome-scale model *i*JN1463 of *P. putida* KT2440 [55] with glucose as the sole carbon source (**Figure 1B**, in green). The flux towards all of the intermediates that contribute to biomass formation (representing the first step of the corresponding catabolic pathways) was computed for each reaction identified *via* FBA (**Table S2**), and the reactions were ranked according to their absolute flux value. With these reactions ‘knocked-out’ in iJN1463 (i.e., an *in silico* C2 auxotroph), the two possible reactions of a bifunctional PKT enzyme acting as both fructose 6-phosphate phosphoketolase (FPK) and xylulose 5-phosphate phosphoketolase (XPK) were introduced into the model (**Figure 1B**, in red).

**Figure 1.**
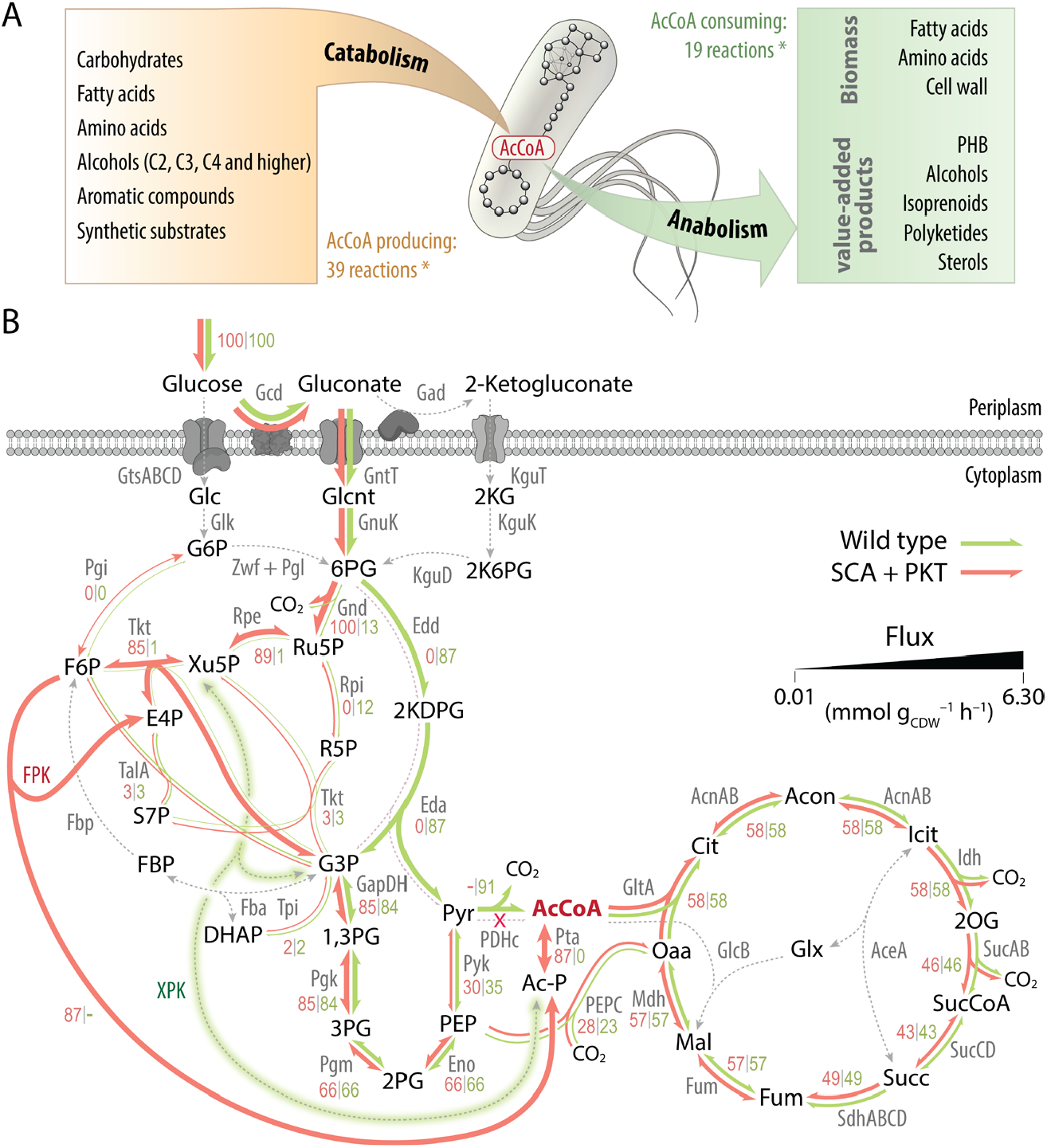
Modeling the role of AcCoA in the central carbon metabolism of *P. putida* towards establishing a synthetic C2 metabolism. **(A)** Numerous carbon substrates are funneled through catabolic pathways with AcCoA as the end- or side-product [* number of reactions listed in the BioByc database [76]]. Conversely, AcCoA serves as the starting point (or co-substrate) for the biosynthesis of many essential biomass constituents or secondary metabolites of industrial interest. *PHB*, poly(3-hydroxybutyrate). **(B)** *In silico* flux distributions estimated via the genome-scale reconstruction iJN1463 of *P. putida* KT2440. The flux distribution of the wild-type model with glucose as the sole carbon source is shown with green arrows; red arrows indicate the fluxes after deleting AcCoA-producing pathways and introducing the PKT reactions. The thicknesses of reaction arrows are drawn in scale reflecting their flux value (from 0.01 to 6.30 mmol gCDW^−1^ h^−1^). Reactions with no flux are drawn as dashed lines. Abbreviations: 1,3PG, 1,3-bisphosphoglycerate; 2K6PG, 2-keto-gluconate-6-P; 2PG, glycerate-2-P; 3PG, glycerate-3-P; 6PG, 6-phosphogluconate; 2KG, 2-ketogluconate; Ac-P, acetyl-P; AcCoA, acetyl-coenzyme A; DHAP, dihydroxyacetone-P; E4P, erythrose-4-P; F6P, fructose-6-P; FBP, fructose-1; G3P, glyceraldehyde-3-P; G6P, glucose-6-P; Glc, glucose; Glcnt, gluconate; KDPG, 2-keto-3-deoxy-6-phosphogluconate; PEP, phosphoenolpyruvate; R5P, ribose-5-P; Ru5P, ribulose-5-P; S7P, sedoheptulose-7-P; 2OG, 2-oxogluconate; Acon, aconitate; Cit, citrate; Fum, fumarate; Glx, glyoxylate; Icit, isocitrate; Mal, malate; Oaa, oxaloacetate; Pyr, pyruvate; Succ, succinate; SucCoA, succinyl-CoA; Xu5P, xylulose-5-P; Abbreviations for enzymes: AceA, isocitrate lyase; AcnAB, aconitase; Eda, 2-dehydro-3-deoxy-phosphogluconate aldolase; Edd, phosphogluconate dehydratase; Eno, enolase; Fba, fructose-1; FPK, fructose-6-P phosphoketolase; Fbp, Fructose-1; Fum, fumarate hydratase; GADH, gluconate dehydrogenase; GapDH, glyceraldehyde-3-P dehydrogenase; Gcd, glucose dehydrogenase; GlcB, malate synthase.; Glk, glucose kinase; GltA, citrate synthase; Gnd, phosphogluconate dehydrogenase; GntT, gluconate/H^+^ symporter; GnuK, gluconate kinase; GtsABCD, D-glucose ABC-transporter; MaeB, malic enzyme; Idh, isocitrate dehydrogenase; KguD, 2K6PG reductase; KguK, 2-ketogluconate kinase; KguT, putative 2-ketogluconate transporter; Mdh, malate dehydrogenase; PEPC, phosphoenolpyruvate carboxylase; PC, pyruvate carboxylase; PDHc, pyruvate dehydrogenase complex; Pgi, phosphohexose-isomerase; Pgk, phosphoglycerate kinase; Pgl, phosphogluconolactonase; Pgm, phosphoglycerate mutase; Pta, phosphotransacetylase; Pyk, pyruvate kinase; Rpe, ribulose-5-P 3-epimerase; Rpi, ribose-5-P isomerase; Sdh, succinate dehydrogenase; SucAB, 2-oxoglutarate dehydrogenase; SucCD, succinyl-CoA synthetase; Tal, transaldolase; Tkt, transketolase; Tpi, triosephosphate isomerase; Zwf, glucose-6-P dehydrogenase. Only compounds participating in carbon transitions are shown, and reactions catalyzed by multiple isoenzymes (e.g., Pgi, Zwf, or GapDH), were lumped together for the sake of simplicity. The numbers shown within the map show the relative fluxes normalized to the glucose uptake rate (6.30 mmol gCDW^−1^ h^−1^, set as 100).

Based on the stoichiometries within the metabolic network, the maximum growth rate achievable with an alternative, PKT-based synthetic metabolism for glucose (0.61 h^−1^) was close to that of the wild-type model (0.62 h^−1^). The ED pathway (flux through Edd and Eda) was completely inactive. Therefore, the PKT pathway constitutes a feasible alternative to the ED pathway in this obligate aerobic bacterium. Under these conditions, all metabolic flux is channeled into the pentose phosphate pathway (PPP), with a 7-fold increased phosphogluconate dehydrogenase (Gnd) activity compared to that in the wild-type model. The flux through the ribulose 5-phosphate 3-epimerase (Rpe) reaction is also greatly enhanced to provide F6P or Xu5P needed as precursors for the PKT reactions.

The flux distribution shown in **Figure 1B** represents only one possible FBA solution in which PKT acts exclusively on F6P. However, there is an infinite number of possible flux distributions in which the total flux of 5.46 is distributed between FPK and XPK, providing equal amounts of Ac-P and G3P. PPP metabolite levels are balanced in each solution by the transaldolase (TalA) and transketolase (Tkt) reactions. In both simulations, glucose was exclusively fed into central metabolism through the 6-phosphogluconate (6PG) route, contrasting with experimentally determined metabolic flux distributions, in which only 78% of the consumed glucose enters the cells in the form of gluconate, 10% as glucose, and 12% as 2-ketogluconate [2KG [29]]. The equilibrating effect of the PPP enzymes potentially allows for a balanced flux through the PK pathway and downstream glycolysis. Hence, no significant increase in anaplerotic reactions is observed. To experimentally complement these *in silico* predictions, a synthetic C2 auxotroph (SCA) of *P. putida* was constructed as explained in the next section.

### A synthetic auxotroph of P. putida requires acetate supplementation to support growth on sugars

Informed by the *in silico* prediction with *i*JN1463, a whole family of SCA strains was created with an increasing number of deletions of genes involved in the potential formation of AcCoA (**Table 1, Table S2**). The auxotrophy was confirmed by testing their growth in DBM medium with various carbon sources (**Figure 2A, Figure S1A-E**). All strains deficient in PDHc (i.e., strains SCA1, SCA3, SCA9, SCA15, and SCA18) could not grow within 120 h after inoculation with glucose as the carbon source (**Figure 2A**, only the first 24 h are shown). These strains could not grow on fructose and gluconate, either (data not shown). Thus, the deletion of *aceE* and *aceF* suffices to establish auxotrophy for C2 compounds on glycolytic substrates in wild-type *P. putida*. Cells that had only PDHc inactivated within central carbon metabolism (strain SCA3) were not impaired in their ability to utilize acetate as the only carbon source (**Figure S1A, Table S3**). Furthermore, when PDHc was left intact, the 14 implemented gene deletions in strain SCA14_PD_ did not affect phenotypes when grown on glycolytic substrates (**Figure 2A**) or citrate, a TCA cycle metabolite and a preferred carbon substrate of *P. putida* (**Figure S1B**). However, removing several genes involved in the assimilation of amino acids, among other targets, resulted in decreased biomass yields and more pronounced diauxic shifts when grown in a rich medium (**Figure 2B**).

**Figure 2.**
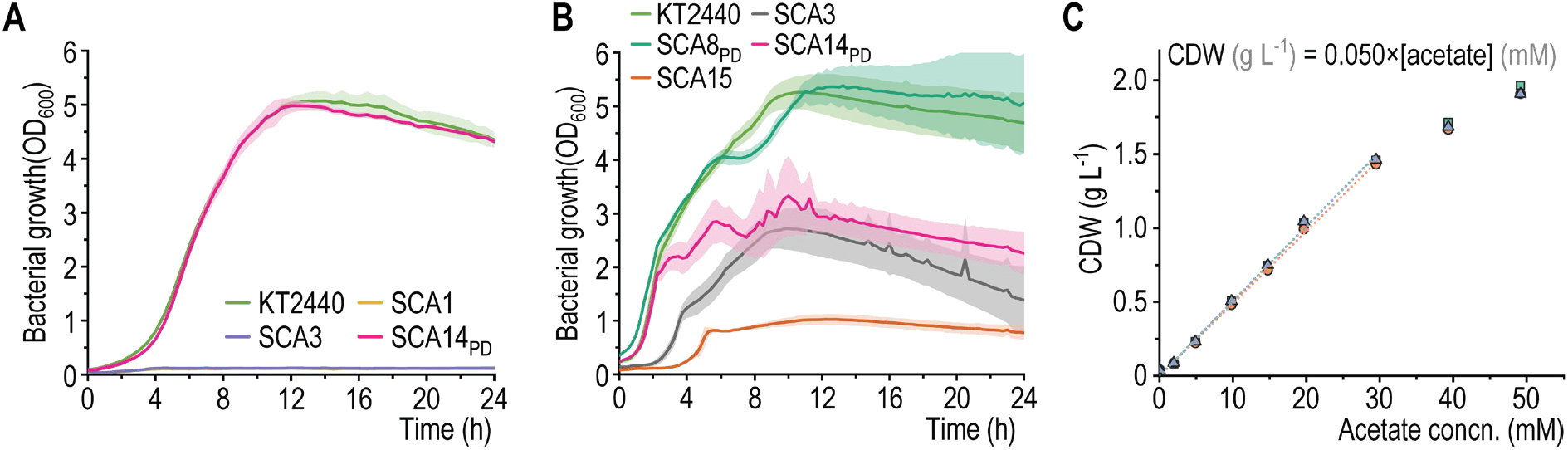
Phenotypic characterization of synthetic C2 auxotrophs. SCA strains and wild-type *P. putida* KT2440 were cultured in microtiter plates (96-well). **(A)** Growth in DBM medium supplemented with 30 mM glucose as the sole source of carbon. **(B)** Growth in LB medium. In all cases, the error bars represent standard deviations from three biological replicates, indicated by shaded ribbons around the growth curves. **(C)** Growth-dependency of strain SCA3 on acetate for biomass formation (indicated as the concentration of cell dry weight, *CDW*). The strain was grown in DBM medium supplemented with 30 mM glucose and varying concentrations of acetate. Shown are the individual measurements and linear correlations of three independent biological replicates. *Concn*., concentration.

Glucose (C6) and acetate (C2) are co-consumed in *P. putida* KT2440 (**Figure S2**). Thereby, glucose is used as an energy source as long as acetate is assimilated. After the depletion of acetate, the carbon skeleton of glucose is utilized for biomass formation. With their co-utilization, it seemed possible to restore growth in SCA strains via the co-feeding of acetate in addition to glucose. To gain insights into the requirements for acetate as co-substrate, strain SCA3 was cultured in DBM medium with 30 mM glucose as well as varying concentrations of potassium acetate (KAc; **Figure S1C**). The maximum biomass concentration reached by the strains showed a linear dependency on acetate when supplied up to 1.8 g L^−1^ (30 mM, **Figure 2C**). At higher cell concentrations, oxygen supply becomes a limiting factor in microtiter plates due to low transfer rates, leading to a disproportional rise in cell densities at increasing substrate concentrations. Importantly, the maximum specific growth rate (µ_*max*_) of strain SCA3 was equal to that of wild-type *P. putida* on glucose and acetate (**Table S3**).

The cumulative effects of twelve additional deletions in SCA15 led to a reduction in µ_*max*_ of ca. one-half while lowering the acetate-specific biomass yield (*Y*_X/S_) by about one-third (**Figure S1D** and **E, Table S3**). The reduced µ_*max*_ and *Y*_X/S_ were likely the results of glyoxylate shunt inactivation *via* deletion of malate synthase (*glcB*), causing all carbon moieties introduced into the TCA cycle in the form of acetate-derived AcCoA to be converted into CO_2_. Thus, essential TCA cycle intermediates need to be replenished from the glycolytic end-metabolites pyruvate and PEP. The inactivation of the glyoxylate shunt also prevented the utilization of acetate as the only carbon source for *P. putida* SCA15 (**Figure S1A**). All SCA strains, including SCA15, were still able to grow on citrate as the sole source of carbon, albeit severely impaired (**Figure S1B**). To rule out any side-activity of citrate synthase in the reverse direction, *gltA* was deleted alongside genes involved in carbon catabolite repression. However, the resulting strain, *P. putida* SCA18, was still able to grow on citrate as the sole carbon source, with no apparent growth deficiencies compared to SCA15. These results form the basis for establishing a synthetic assimilation route for sugars as indicated below.

### Quantitative proteome analysis exposes the dependency of synthetic C2 auxotrophs on acetate and highlights bottlenecks for a functional phosphoketolase pathway

To uncover the effects of removing intracellular AcCoA sources, the quantitative proteome of exponentially-growing strains SCA3 and KT2440 was assessed via mass spectrometry. Cells were grown in DBM medium supplemented with 30 mM glucose and 30 mM acetate, whereby a total of 110 proteins differed significantly [false discovery rate (FDR) of ≤ 0.05] with an absolute log_2_(fold change) of ≥ 1 (**Figure S3**). Among these, 45 were identified to have metabolic functions (**Table S4**). Within central carbon metabolism, a down-regulation of gluconokinase (GnuK), glucose 6-phosphate 1-dehydrogenase [ZwfA [77]], and 6-phosphogluconolactonase (Pgl) was observed together with an up-regulation of 2-keto-gluconate-6-P (2K6PG) reductase (KguD; **Figure S4**). This pattern suggests a shift from the preferred glucose utilization route [i.e., glucose oxidation and gluconate uptake [29]] towards further oxidation to 2KG and utilization thereof. We had observed before that engineered *P. putida* strains convert supplied glucose into 2KG at significant rates if growth is hampered by genetic modifications [78].

Strain SCA3 showed a diverging pattern in the expression of enzymes involved in the anaplerotic reactions connecting the downstream EMP route with the TCA cycle. Compared to wild-type, the oxaloacetate-producing pyruvate carboxykinase (PycA and PycB) was down-regulated by 70%. On the other hand, oxaloacetate decarboxylase (OadC, *PP_1389*) was up-regulated by a factor of 6.4. The two enzymes constituting the glyoxylate shunt, isocitrate lyase (AceA) and malate synthase (GlcB), were 3.5- and 2.0-fold more abundant in SCA3. Furthermore, two enzymes associated with fatty acid degradation (and conversely, AcCoA formation), AcCoA acetyltransferase (YqeF, 3.5-fold) and β-ketoadipyl-CoA thiolase subunit β (FadA, 1.6-fold)] were elevated. (3*R*)-3-Hydroxydecanoyl-ACP dehydratase (FabA), associated with fatty acid biosynthesis, was down-regulated by a factor of 0.6. The results of pathway enrichment analysis with pathways listed in the KEGG database are shown in **Figures S5** and **S6**. These observations suggest a substantial utilization of acetate as a carbon substrate for central metabolism as opposed to energy provision. The abundance of fatty acid metabolism enzymes reflects the recruitment of lipid pools to replenish AcCoA, whereas the activity of the glyoxylate shunt is essential for acetate utilization since it bypasses the two decarboxylation steps of the TCA cycle. The thereby conserved carbon could be further fed into gluconeogenesis, as indicated by OadC overexpression.

In an attempt to identify potential bottlenecks towards a synthetic metabolism based on the PKT reaction, the relative protein abundances for strain KT2440 grown on glucose were compared to that during growth on glucose and acetate (**Figure S7**). Most enzymes within the upstream glycolysis were produced at statistically significantly lower amounts in the presence of acetate as co-substrate. Only Pta was expressed 8-fold higher in the presence of glucose and acetate compared with sugars as the sole carbon source. The functionality of Pta in the conversion of Ac-P to AcCoA was experimentally demonstrated [28], and the low abundance in glucose-grown *P. putida* suggests a potential bottleneck for a functional PKT pathway. Additionally, the low flux through the PPP in *P. putida* [29] was also reflected in the protein levels of all PPP enzymes in all experimental conditions, except for Tkt and Tal. Flux control over these reactions is likely exerted on a metabolite level [79], likely allowing for a quicker response to changes in the flux distribution within the EDEMP cycle. These observations indicate that the fluxes through the non-oxidative branch of the PPP and the conversion of Ac-P to AcCoA should be enhanced in order to establish a synthetic PKT metabolism in *P. putida*.

### Implementing the phosphoketolase pathway in a synthetic C2 auxotroph restores prototrophy on glucose

The PKT pathway was implemented by introducing the gene encoding Xfpk^Ba^ from *Bifidobacterium adolescentis* E194, a bifunctional PKT enzyme with a reported *in vitro* activity ratio on Xu5P:F6P of 3:2 [80], in a landing pad in the intergenic region between *PP_0013* (*gyrB*) and *PP_5421* [45] in the chromosome of *P. putida* SCA3, yielding strain SCA3_PK_ (**Table 1**). Additionally, the strain was transformed with plasmid pS438·*pta*^*Ec*^ (encoding the highly-active Pta enzyme from *E. coli*) to boost the Ac-P conversion into AcCoA. In this plasmid, *pta*^*Ec*^ is placed under transcriptional control of the XylS/*Pm* expression system, and the construct also provides XylS *in trans* to initiate transcription from the *Pm* promoter that leads to the expression of *xfpk*^*Ba*^ in the chromosome.

The production and functionality of Xfpk^*Ba*^ were tested *in vitro* with cell lysates via indirect chemical detection through the hydroxamate method (**Figure 3A**). To this end, the soluble fraction of cell-free extracts obtained from strains SCA3/pS438·*pta*^*Ec*^ and SCA3_PK_/pS438·*pta*^*Ec*^ were adjusted to a final total protein concentration of 6 mg mL^−1^, and the F6P-dependent formation of Ac-P was monitored over time. After 1 h of incubation, PKT activity in the sample containing Xfpk^Ba^ was evident by the development of a red pigment, whereas no such coloration was observed in the absence of Xfpk^Ba^ (**Figure 3B**). **Figure 3C** shows the time-resolved PKT activity of the two cell lysates. No formation of Ac-P was observed for the strain without *xfpk*^*Ba*^ (*P. putida* SCA3/pS438·*pta*^*Ec*^), while lysates of strain SCA3_PK_/pS438·*pta*^*Ec*^ converted F6P into Ac-P at a rate of 0.049 mmol Ac-P min^-1^, corresponding to a specific activity of 0.0082 mmol mg_protein_^−1^ min^−1^. The results of the hydroxamate assay indicated that Xfpk^Ba^ was produced and functionally active in *P. putida*. Two other Xfpk variants, chromosomally introduced in SCA3 as codon-optimized sequences of the *xfpk* genes from *Pseudomonas fluorescens* F113 and *Bifidobacterium mongoliense* YIT 10443, were tested under the same conditions but showed no significant formation of Ac-P in cell-free extracts of the corresponding engineered strains (data not shown).

**Figure 3.**
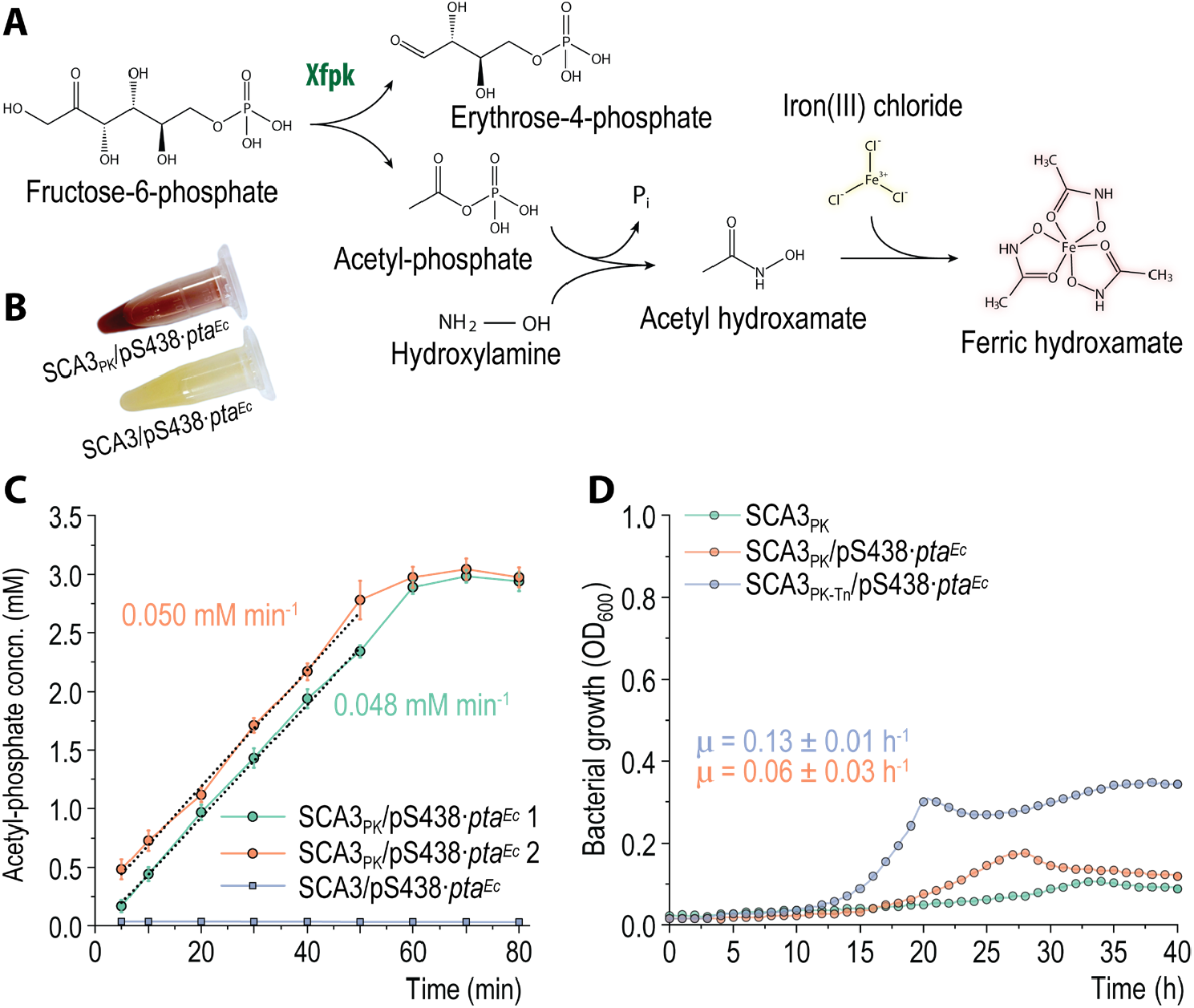
Expression of *xfpk*^*Ba*^ in a synthetic C2 auxotroph strain. Cell lysates were prepared from cultures grown in LB medium supplemented with 0.5 mM 3-*m*Bz. Total protein concentrations of 6 mg mL^−1^ were used for the conversion of F6P to Ac-P, the concentration of which was determined using the hydroxamate method. **(A)** Reaction scheme of the PKT assay via ferric hydroxamate. **(B)** Reaction samples illustrating the enzymatic conversion Ac-P to ferric hydroxamate after 1 h incubation. The red color in the assay for strain SCA3PK/pS438·*pta*^*Ec*^ indicates Ac-P formation. **(C)** *In vitro* determination of FPK activity in *P. putida* SCA3/pS438·*pta*^*Ec*^ and SCA3PK/pS438·*pta*^*Ec*^. Each strain was tested with two biological replicates with three technical replicates; only one of two (comparable) biological replicates is shown for strain SCA3/pS438·*pta*^*Ec*^. Error bars represent standard deviations. **(D)** *In vivo* demonstration of PKT activity via growth of strain SCA3 harboring a chromosomally integrated *xfpk*^*Ba*^ gene with and without plasmid pS438·*pta*^*Ec*^ or overexpressed *gntZ* and *rpe*. Cells were cultured in a microtiter plate (96-well) in DBM medium supplemented with 10 mM glucose and 0.5 mM 3-*m*Bz. The OD600 increase for strain SCA3PK could be attributed to the formation of colored compounds produced from 3-*m*Bz *via* oxidation. *Concn*., concentration.

To test whether Xfpk^Ba^ could restore the growth of synthetic C2 auxotroph strains, SCA3_PK_/pS438·*pta*^*Ec*^ and SCA3/pS438·*pta*^*Ec*^ were cultured in DBM medium supplemented with 10 mM glucose as well as 0.5 mM 3-*m*Bz as the inducer of *xfpk*^*Ba*^ expression (**Figure 3D**). Indeed, the expression of both *xfba*^*Ba*^ and *pta*^*Ec*^ enabled glucose-dependent growth of *P. putida* SCA3_PK_/pS438·*pta*^*Ec*^, with a *Y*_X/S_ of 0.02 g_CDW_ gglc^−1^ and a µ_*max*_ of 0.06 h^−1^. Interestingly, expressing *pta*^*Ec*^ alone did not promote bacterial growth (data not shown), indicating that Xfpk^Ba^ is involved in sugar catabolism.

As indicated in the previous sections, the reactions performed by 6-phosphogluconate dehydrogenase (Gnd) and ribulose 5-phosphate 3-epimerase (Rpe) were elevated in the optimal flux prediction obtained with *i*JN1463 for a C2-auxotrophic design compared to the wild-type model. Thus, the expression of *gntZ* and *rpe* was artificially enhanced *via* the chromosomal integration of a Tn7-encoded, synthetic *gntZ-rpe* operon. The expression of *gntZ* was controlled by the strong, constitutive P_*14g*_ promoter [81] and the bi-cistronic translational coupling sequence *BCD10* [82]. In this design, the *rpe* gene was translationally coupled to *gntZ*. Chromosomal integration was achieved by cloning the synthetic operon into a mini-Tn*7* transposon borne by plasmid pTn*7*·P_*14g*_*(BCD10)*→*gntZ-rpe* (**Table 2**). Overexpression of *gntZ* and *rpe* led to a 2- and 1.7-fold increase in µ_*max*_ and *Y*_X/S_, respectively, for the resulting strain SCA3_PK-Tn_/pS438·*pta*^*Ec*^ compared to SCA3_PK_/pS438·*pta*^*Ec*^ (**Figure 3D, Table 3**). Hence, and as predicted by FBA simulations, the implementation of a PKT shunt combined with increasing Gnd, Rpe, and Pta production proved to be a successful strategy to restore growth in a C2-auxotroph strain. The growth rate and biomass yield observed for strain SCA3_PK_/pS438·*pta*^*Ec*^ were deemed sufficient to support evolutionary engineering towards enhancing glucose-dependent growth of engineered *P. putida*.

**Table 3.** Quantitative physiology parameters of glucose-grown PKT-shunt strains.

| *P. putida* strain | Modifications | | | Carbon source | $\mu_{max}$ (h<sup>-1</sup>) | Biomass yield, $Y_{X/S}$ (g<sub>CDW</sub> g<sub>S</sub><sup>-1</sup>) |
| --- | --- | --- | --- | --- | --- | --- |
|  | *xfpkBa* | *gntZ-rpe* | Evolved |  |  |  |
| SCA3 | – | Native | – | Glucose | N.G. | N.G. |
| SCA3<sub>PK</sub>/pS438·*pta*<sup>Ec</sup> (pre-evolved/evolved) | + | Native | – / + | Glucose | 0.06 ± 0.03 | 0.02 ± 0.01 |
| SCA3<sub>PK-Tn</sub>/pS438·*pta*<sup>Ec</sup> (pre-evolved) | + | Overexpressed | – | Glucose | 0.13 ± 0.01 | 0.04 ± 0.01 |
| SCA3<sub>PK-Tn</sub>/pS438·*pta*<sup>Ec</sup> (evolved) | + | Overexpressed | + | Glucose | 0.28 ± 0.03 | 0.11 ± 0.01 |
| SCA3<sub>PK-Tn</sub>/pS438·*pta*<sup>Ec</sup> (evolved) | + | Overexpressed | + | Gluconate | 0.23 ± 0.03 | 0.04 ± 0.01 |

Maximum exponential growth rates ( $\mu_{max}$ ) were determined via Gaussian process regression for growth experiments in microtiter plates. Biomass yields ( $Y_{X/S}$ ), in g of cell dry weight (g<sub>CDW</sub>) per g of substrate (g<sub>S</sub>), were calculated for each indicated substrate. Given are the average values ± standard deviation from at least three biological replicates. N.G., no growth.

### Adaptive laboratory evolution accomplished the integration of the phosphoketolase pathway into the metabolic network of P. putida

To optimize the flux through the synthetic PKT metabolism, strains SCA3_PK-Tn_/pS438·*pta*^*Ec*^, SCA3, SCA3/pS438·*pta*^*Ec*^, and SCA3_Tn_/pS438·*pta*^*Ec*^ were subjected to ALE. After an estimated number of 140 generations (ca. 2 months) in DBM medium containing glucose, no significant difference in growth was observed for *P. putida* strains SCA3, SCA3/pS438·*pta*^*Ec*^, and SCA3_Tn_/pS438·*pta*^*Ec*^ (results not shown). However, strain SCA3_PK-Tn_/pS438·*pta*^*Ec*^ had a significant improvement in growth parameters, and µ_*max*_ and *Y*_X/S_ increased by 2- and 3-fold, respectively (**Figure 4A, Table 3**, and **Figure S8A, B**). The enhanced growth observed for the evolved strain on glucose was stable even after five sub-culturing steps in a rich medium (LB). Hence, evolution—rather than a mere adaptation—was responsible for the improved phenotype.

**Figure 4.**
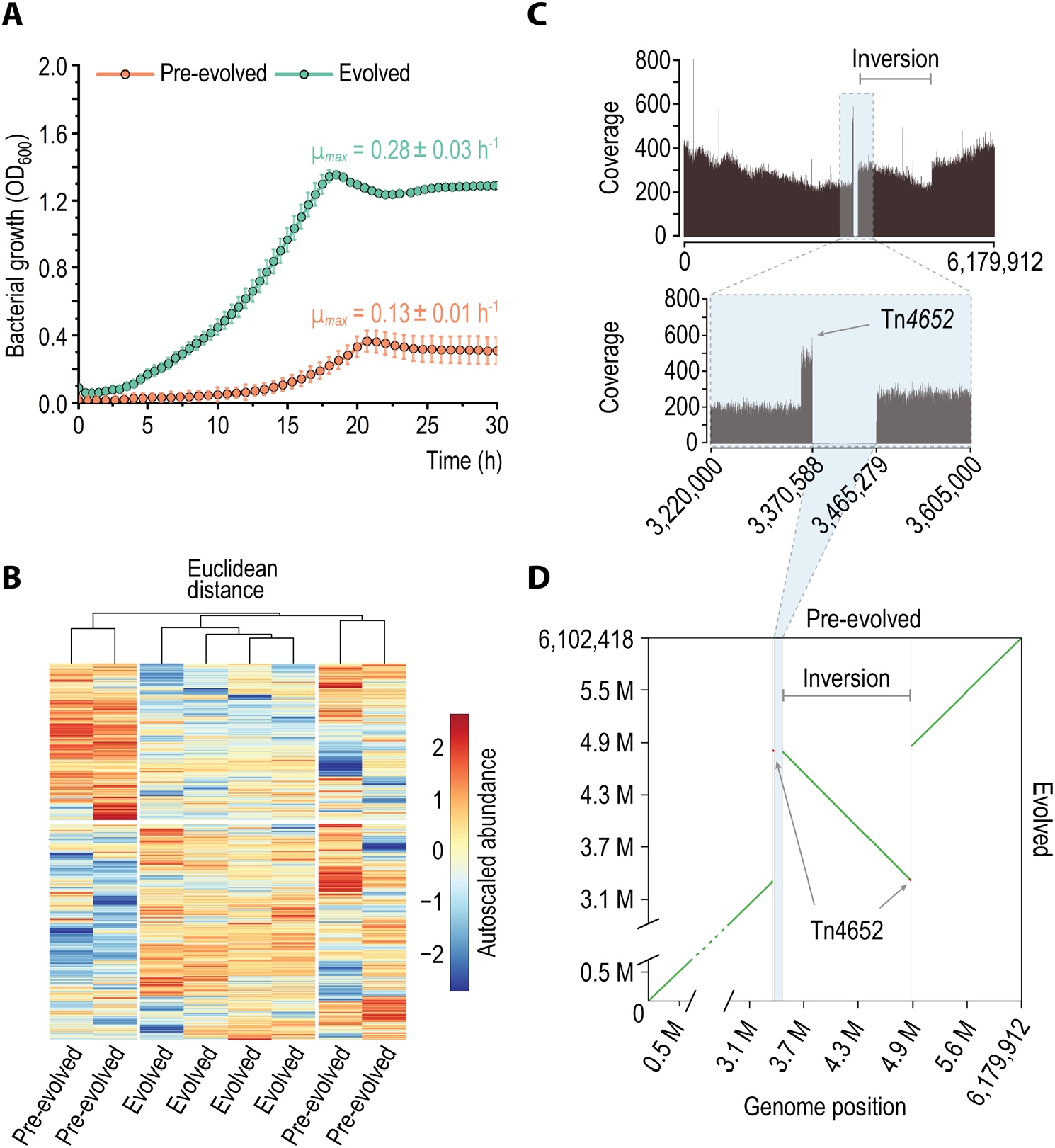
Evolution of synthetic PKT metabolism in engineered *P. putida*. **(A)** Batch (24-well plate) cultivation of pre-evolved and evolved SCA3PK-Tn/pS438·*pta*^*Ec*^ in DBM medium supplemented with 30 mM glucose and 0.5 mM 3-*m*Bz. *OD600*, optical density at 600 nm. **(B)** Network-wide proteome of the evolved and pre-evolved *P. putida* strain. The protein abundance values were normalized, auto-scaled, and plotted according to their Euclidean distance for four biological replicates. **(C)** Coverage plot of whole-genome sequencing data (Illumina SBS technology) mapped to the reference chromosomal sequence of SCA3PK-Tn/pS438·*pta*^*Ec*^. The progression of sequencing coverage downstream of the deletion indicates the inversion of a large chromosomal segment. **(D)** Dot plot of the reconstructed chromosomal sequence of evolved SCA3PK-Tn/pS438·*pta*^*Ec*^ compared to the reference sequence in the pre-evolved strain. A duplicated DNA segment comprising the Tn*4652* transposon is illustrated in red, the deleted stretch is shaded in blue.

When grown on gluconate (i.e., a more oxidized sugar than glucose), the growth rate of the evolved clones was not significantly altered. However, evolved SCA3_PK-Tn_/pS438·*pta*^*Ec*^ reached only 36% of the cell density obtained on glucose (**Figure S8B**). This *Y*_X/S_ reduction suggests a stoichiometric bottleneck in the production of either energy or reducing equivalents on gluconate. During growth on glucose, 26% of membrane-bound ubiquinol in the electron transport chain is predicted to be produced by direct glucose oxidation (as suggested by FBA with *i*JN1463), acting as an ATP source. Alternatively, *P. putida* can produce NAD(P)H via the oxidation of glucose 6-phosphate, contributing to the intracellular pool of reducing equivalents [77]. The basis for these rearrangements was investigated by quantitative proteomics as disclosed in the next section.

### Comparative proteomic profiling of pre-evolved and evolved cells carrying a synthetic PKT metabolism

Comparative proteome analysis of glucose-grown SCA3_PK-Tn_/pS438·*pta*^*Ec*^ cultures before and after evolution revealed only 28 proteins with significantly altered abundance (**Figure S9**). However, the pre-evolved clones exhibited noticeably higher variability in their protein content, reflected in large Euclidean distances between each clone’s dataset (**Figure 4B**), and we performed pathway enrichment analysis on the proteins that were found to differ significantly (**Figure S10**). Most notably, ThiL, encoding thiamine monophosphate kinase, could not be detected in the proteome of pre-evolved *P. putida* SCA3_PK-Tn_/pS438·*pta*^*Ec*^, whereas it was present in all replicates of the evolved strain. Overexpression of *thiL* could be the result of increased thiamine pyrophosphate demands due to the production of Xfpk. Secondly, the ethanolamine ammonia-lyase EutB was highly overrepresented in all evolved clones, and this gene had been added to the list AcCoA-forming reactions (**Table S2**) due to its potential to form acetaldehyde (and consequently, AcCoA) from ethanolamine, the most abundant phospholipid head group in bacteria. Lastly, the abundance of GltD (glutamate synthase subunit β) was increased 3-fold. These three metabolic functions could not adequately account for the changes observed in glucose-dependent growth of the engineered *P. putida* strains, and we adopted functional genomic approaches to study potential modifications brought about by ALE.

### Large-scale structural reorganizations occurred via transposition of mobile genetic elements

Four evolved SCA3_PK-Tn_/pS438·*pta*^*Ec*^ clones with enhanced growth on glucose, and four pre-evolved isolates were subjected to whole-genome sequencing. Polymorphisms that were identified in both pre-evolved and evolved clones were discarded, and no mutations were found in plasmid pS438·*pta*^*Ec*^. In the sequenced chromosome, a segment of 94,691 bp was absent in the read coverage of evolved SCA3_PK-Tn_/pS438·*pta*^*Ec*^ clones (**Figure 4C**). This chromosomal region stretches from *PP_2985* to *PP_3089*, and harbors the previously identified 35.6-kb long prophage 2 [*PP_3026*-*PP_3066* [83,84]*]. The loss of this prophage has not been reported so far in P. putida* KT2440. Furthermore, the excised segment is located 1,111 bp downstream of the transposon Tn*4652*, which was shown to be activated under carbon starvation that can create novel fusion promoters on both ends upon insertion into a target locus with low target sequence specificity [85–88]. In the course of evolution, Tn*4652* was duplicated and inserted into the gene encoding the flagellar biosynthesis protein FlhA, thereby disrupting the open reading frame. Removal of the flagellar machinery has been shown to beneficially affect the energy status and reducing power availability of *P. putida* KT2440, resulting in multifaceted physiological improvements [89]. Downstream of the insertion site, the flagellar operon continues with *PP_4345*, encoding a GntR family transcriptional regulator, and *ddlA*, encoding D-alanine−D-alanine ligase A. Accordingly, DdlA was found to be significantly overrepresented in the proteome of evolved SCA3_PK-Tn_/pS438·*pta*^*Ec*^ clones (**Figure S9**), likely the result of enhanced transcriptional activity downstream of Tn*4652*. The 1,458,179-bp chromosomal segment flanked by both Tn*4652* copies was inverted (**Figure 4D**), which could have wide-ranging effects on the three-dimensional structure of the chromosome and therefore on gene regulation.

A full list of genes located within the deleted 94.7-kb region is given in **Table S5**. The absence of the 94.7-kb chromosomal sequence was confirmed in the respective clones *via* PCR-amplification of the region (**Figure S11**). Apart from many proteins with unknown functions, an array of functions contributing to the production of pyocins are encoded within this region, as well as proteins with predicted regulatory and enzymatic roles. Among the potential regulatory proteins are FlaR (DNA topology modulation kinase), PP_3006 (ECF family RNA polymerase σ^70^ factor), PP_3002 (AraC family transcriptional regulator), ClpP (ATP-dependent protease), PP_5558 (Xre family transcriptional regulator), PP_3086 (RNA polymerase σ^70^ factor), PP_3075 (putative transcriptional regulator), and PP_3087 (excinuclease ABC subunit A). While the exact functions of these proteins have not been characterized, the loss of *flaR, PP_3006, clpP, PP_3086*, and *PP_3086* could have caused systemic effects on DNA topology, DNA transcription, or protein degradation. Within the set of enzymes encoded in the missing chromosomal region, *PP_3071*, encoding an acetoacetyl-CoA synthetase, could have a direct effect on the intracellular AcCoA concentrations. The next step was to evaluate the performance of evolved clones of *P. putida*, bearing a synthetic PKT metabolism, under conditions compatible with sugar-dependent bioproduction.

### The synthetic PKT metabolism enables conservation of carbon during growth on glucose

To characterize the physiology of evolved PKT-utilizing *P. putida* SCA3_PK-Tn_/pS438·*pta*^*Ec*^, aerobic batch fermentations were performed in bioreactors under controlled conditions and compared to wild-type strain KT2440 (**Figure 5A-D**). Glucose was used as the only carbon substrate at 30 mM. **Figure 5A** shows the time-resolved measurements for the dissolved oxygen concentration (DO), biomass, differential CO_2_ formation rates, and extracellular metabolite concentrations (i.e., glucose, Glcnt, 2KG, and pyruvate).

**Figure 5.**
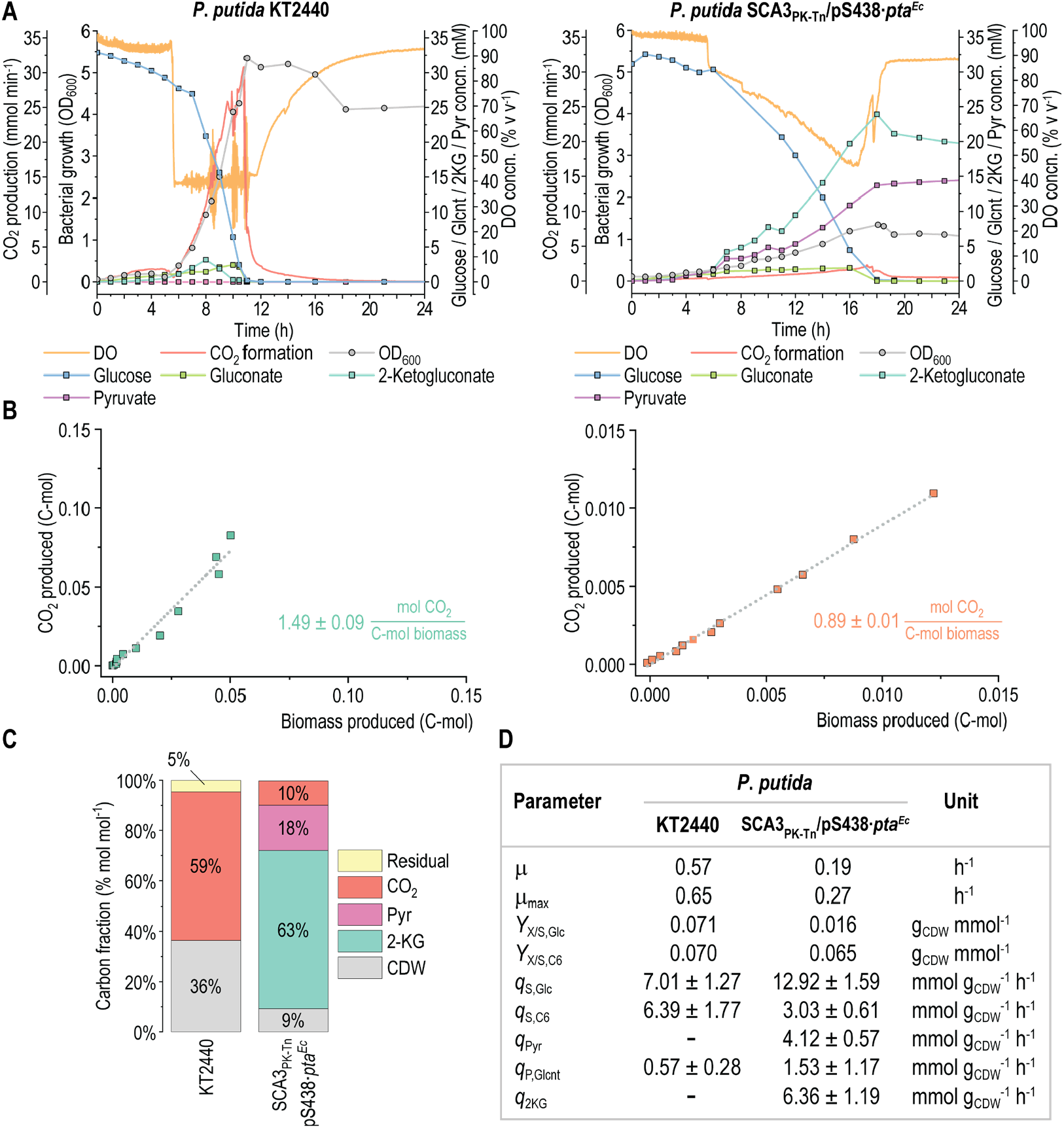
Bioreactor cultivation of *P. putida* KT2440 and evolved SCA3PK-Tn/pS438·*pta*^*Ec*^. Cells were grown in lab-scale bioreactors in batch mode in DBM medium supplemented with 30 mM glucose and, for SCA3PK-Tn/pS438·*pta*^*Ec*^, 0.5 mM 3-*m*Bz. **(A)** Selected process parameters and concentrations of substrates, metabolites, and biomass over time. No pyruvate could be detected in the medium of wild-type strain KT2440. *OD600*, optical density at 600 nm. **(B)** Correlation of the cumulative CO2 produced (C-mol) with biomass (C-mol) at different sampling time points. **(C)** Carbon balance for batch fermentations. Shown are all carbon-containing entities measured at the end of the growth phases for each strain; carbon fractions represent the relative value of each entity compared to the overall amount of C-mol at the beginning of the fermentations. The amount of CO2 was determined via the integration of measured CO2 production rates over the whole fermentation time. **(D)** Summary of physiological parameters determined for the bioreactor cultivations. *Concn*., concentration.

Wild-type strain KT2440 reached its maximum cell density at an OD_600_ of 5.4 within 11 h (µ = 0.57 h^−1^), while strain SCA3_PK-Tn_/pS438·*pta*^*Ec*^ grew with a µ = 0.19 h^−1^ to a maximum OD_600_ of 1.5 within 18 h. Accompanied by transient secretion of Glcnt (≤ 2.4 mM) and 2KG (≤ 3.2 mM), the wild-type *P. putida* strain consumed the substrate at *q*_S_ = 6.39 ± 1.77 mmol g_CDW_^−1^ h^−1^ (all three forms of the sugar) over the exponential growth phase (**Figure 5D**), and the *Y*_X/S_ on glucose was 0.40 g_CDW_ g_Glc_^−1^. No metabolites other than the three six-carbon moieties were detected in the medium.

The growth profile of strain SCA3_PK-Tn_/pS438·*pta*^*Ec*^ closely resembled that previously observed during physiological characterizations in 24-well plates (**Figure 4A, Figure S8**). Glucose and its two oxidized products were consumed with a combined *q*_S_ = 3.03 ± 0.61 mmol g_CDW_^−1^ h^−1^. Biomass was formed with a *Y*_X/S_ of 0.47 g_CDW_ g_C6_^−1^ and 0.09 g_CDW_ g_Glc_^−1^. Besides generating biomass, SCA3_PK-Tn_/pS438·*pta*^*Ec*^ additionally secreted significant amounts of pyruvate (14.6 mM) into the medium with a specific rate of 0.35 ± 0.05 g_Pyr_ g_CDW_^−1^ h^−1^. Pyruvate production occurred in parallel to glucose consumption, and no re-consumption occurred after the cells reached the stationary phase. The accumulation of pyruvate indicates a strong activity of the ED pathway, involving the splitting of 2KDPG into pyruvate and G3P.

The synthetic PKT metabolism reduced the amount of CO_2_ produced per C-mol of biomass by 46% compared to the original metabolic architecture in the wild-type strain (**Figure 5B**). However, two-thirds of the supplied glucose was oxidized to 2KG, which was secreted into the medium at a specific rate of 1.37 ± 0.26 g_2KG_ g_CDW_^−1^ h^−1^. With a significant fraction of glucose being exclusively used as an energy source, the biomass yield from the consumed carbon was higher in SCA3_PK-Tn_/pS438·*pta*^*Ec*^ compared to wild-type strain KT2440. **Figure 5C** shows the carbon balance at the end of the growth phase for strain KT2440 (11 h) and SCA3_PK-Tn_/pS438·*pta*^*Ec*^ (18 h), including all measured carbon-containing entities. A small residual amount of carbon could not be attributed to any of the measured components for strain KT2440, likely due to inaccuracies in the measurement of biomass concentrations or the volume left in the bioreactor. Both strains had fully consumed all supplied glucose by the end of the growth phase. The only carbon species identified for KT2440 were biomass (ca. 33% of the total carbon balance) and CO_2_ (representing the remaining 64%). On the other hand, strain SCA3_PK-Tn_/pS438·*pta*^*Ec*^ converted 81% of the supplied carbon into 2KG and pyruvate. The cause of this drastic overflow metabolism was explored in detail by unraveling the distribution of intracellular metabolic fluxes as explained below.

### ^13^C-based metabolic flux analysis reveals a hybrid Entner-Doudoroff−phosphoketolase metabolism

The co-utilization of the ED pathway and the synthetic PK shunt resulted in the pronounced overflow of 2KG and pyruvate, suggesting an imbalance in the dissimilation of glucose towards biomass formation. To thoroughly characterize this metabolic architecture, we performed labeling experiments with [3-^13^C_1_]-glucose, [4-^13^C_1_]-glucose, or a mixture of each 50% unlabelled and uniformly labeled glucose ([U-^13^C_6_]-glucose, **Figure 6**). The high conservation of glucose ^13^C-3 and ^13^C-4 in the metabolites of interest allowed for the precise resolution of every flux node within central metabolism, including reactions to and contributions of the two Xfpk activities, acting on either F6P or Xu5P. The use of 50% [U-^13^C_6_]-glucose, in contrast, helped to further increase the accuracy in the determination of flux ratios in the PPP as well as the cycling within the EDEMP architecture. Strain SCA3_PK-Tn_/pS438·*pta*^*Ec*^ produced CO_2_ at a rate that amounts to 20% of wild-type *P. putida* KT2440 (**Figure S12**), as determined for the bioreactor fermentations, which underscores the significant re-routing of carbon towards metabolite and biomass formation rather than oxidative decarboxylation. The PDHc reaction was entirely inactive in the network, as was expected after deleting *aceEF*.

**Figure 6.**
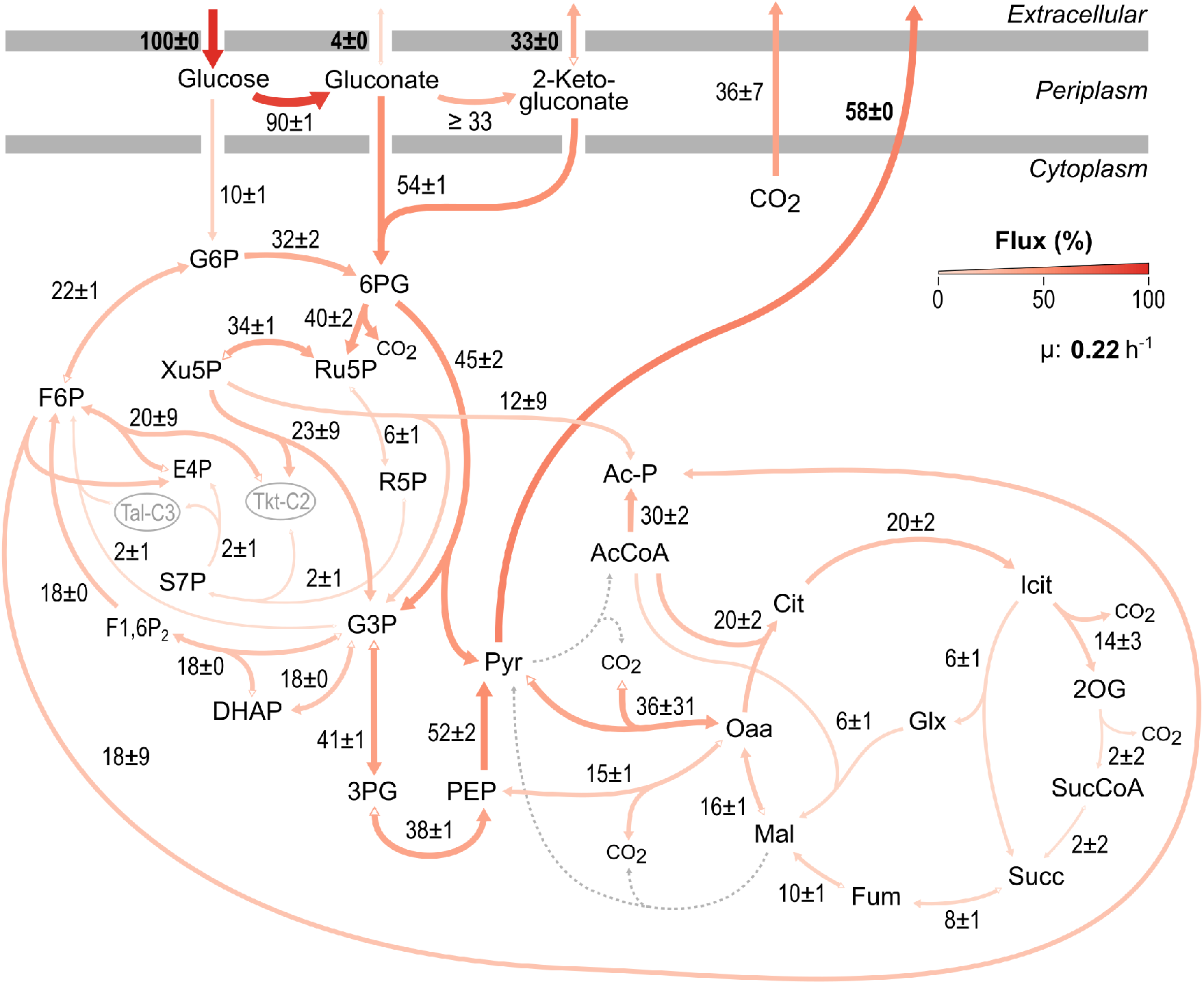
*In vivo* distribution of carbon fluxes in the evolved *P. putida* SCA3PK-Tn/pS438·*pta*^*Ec*^ strain grown on glucose. Experimentally-determined rates were set as fixed parameters: glucose consumption rate, 14.08 ± 1.15 mmol gCDW^−1^ h^−1^; gluconate secretion rate, 0.52 ± 0.07 mmol gCDW^−1^ h^−1^; 2-KG secretion rate, 4.65 ± 0.22 mmol gCDW^−1^ h^−1^; pyruvate secretion rate, 8.19 ± 0.18 mmol gCDW^−1^ h^−1^; and exponential growth rate, 0.22 ± 0.01 h^−1^. All fluxes were normalized to the specific glucose consumption rate. Flux values represent the mean ± standard deviations from three biological replicates. Abbreviations are as indicated in the legend of Figure 1.

Consistent with initial predictions *via* FBA, high activities within the PPP support the utilization of the new Xfpk-based network architecture. Overexpression of *gntZ* and *rpe* resulted in an equal partitioning between the ED pathway and the oxidative PPP, and almost all of the Ru5P generated was converted to Xu5P. One-third thereof was cleaved by Xfpk into G3P and Ac-P. The remaining two-thirds were used by Tkt to recycle the E4P formed *via* the FPK activity of Xfpk back to F6P. This resulted in a significant flux towards G3P through the PPP, of which a major fraction (65%) was recycled through the EDEMP cycle to produce reducing equivalents *via* G6P oxidation (and 6PG decarboxylation). The G3P that was formed through the ED pathway was further processed in the downstream glycolysis to PEP and Pyr. Around 70% of the ED-derived Pyr was secreted into the medium, while the remainder was deployed in biomass-forming pathways or funneled into the TCA cycle through CO_2_-incorporating anaplerotic reactions. Half of the AcCoA formed through the FPK and XPK reactions was used for the production of biomass constituents; the rest fueled the TCA cycle. In there, AcCoA was partially utilized to produce the essential biomass precursor 2-OG (concomitant with the formation of NADH and CO_2_). The remaining carbon within AcCoA was preserved by the glyoxylate shunt.

Stark contrasts were revealed when examining the energy metabolism of wild-type *P. putida* KT2440 and strain SCA3_PK-Tn_/pS438·*pta*^*Ec*^. In the wild-type strain, approximately 60% of the reducing equivalents used for ATP generation *via* oxidative phosphorylation were formed in the TCA cycle in the form of NADH and FADH_2_ (**Figure S12, Table S6**). The remaining energy-generating reducing equivalents were produced *via* G3P oxidation (14%) as well as periplasmic glucose oxidation (22%). In *P. putida* SCA3_PK-Tn_/pS438·*pta*^*Ec*^, this balance was quite different. Less than 1% of the electrons for the respiratory chain were generated in reactions of the TCA cycle, whereas the two periplasmic glucose oxidation steps provided the largest contribution to the NADH, FADH_2_, and UQH_2_ pools. The remainder was provided by the oxidation of G3P.

## Discussion and Conclusions

### A synthetic C2 auxotroph P. putida provides a tight selection regime for AcCoA-producing pathways and reveals unknown metabolic connectivities on gluconeogenic substrates

In this study, we employed a genome-scale model-guided approach to design synthetic C2-auxotroph selection platforms based on *P. putida* KT2440. The auxotrophy of the created strains was experimentally confirmed and shown to be reversible by external acetate addition. Thus, the SCA strains can be easily propagated by supplying acetate or by cultivation in complex media. The strong dependency of biomass formation on acetate provides a tight selection scheme with a large evolutionary landscape in which gradual improvements in the AcCoA supply stimulate fitness gains.

The deletion of *aceEF* proved to be sufficient to disrupt growth on glycolytic substrates. For strain SCA3, in which deletion of *aceEF* was combined with Δ*bkdAA* and Δ*ltaE*, no effect of evolution on restoring prototrophy on glucose was observed. However, one AcCoA-forming enzyme that was targeted in our in silico strain design, EutB, was overrepresented in evolved SCA3_PK-Tn_/pS438·*pta*^*Ec*^. The most abundant phospholipid phosphatidylethanolamine in bacteria is synthesized exclusively by phosphatidylserine decarboxylase [90,91]. Thus, phosphatidylethanolamine biosynthesis and its degradation to acetaldehyde could be an alternative AcCoA source.

The additional deletions identified might be a prerequisite for the selection strategy to work with carbon substrates other than (acidic) sugars. This was illustrated by the ability of all SCA strains to use citrate as the sole carbon source. In strain SCA15, all genes identified as sources of AcCoA, Ac-P, acetate, or acetaldehyde were deleted, and the *in silico* reconstruction of this strain was unable to form biomass in all FBA simulations. However, the strain still showed growth on citrate. Consequently, the additional deletion of *glcB, gltA*, and *prpC* was performed to rule out the reversibility of the three as irreversible described reactions malate synthase [92], citrate synthase [93,94], and 2-methyl citrate synthase [95,96].

The elimination of GlcB, GltA, and PrpC did not affect the growth of SCA18 on citrate. Hence, there appear to be unknown enzymatic functions in *P. putida* that establish a biochemical route from citrate to AcCoA, highlighting the role of ‘underground’ metabolic functions that come into play in the absence of the main reactions that supply biomass precursors or redox equivalents [50].

### The evolutionary integration of the phosphoketolase pathway resulted in a network-wide genome and proteome restructuring

The predictions obtained by FBA informed our strategy to implement an alternative PK-based glycolysis in a synthetic C2-auxotroph strain. Chromosomal integration of *B. adolescentis xfpk* and plasmid-based expression of *E. coli pta* provided sufficient flux through the PKT pathway to establish growth on glucose. In response to the low abundances of GntZ and Rpe observed in whole-proteome analyses, the production of both enzymes was artificially increased, further enhancing the growth on glucose. Adaptive evolution enabled strain SCA3_PK-Tn_/pS438·*pta*^*Ec*^ to make another leap toward restoring its growth ability on glucose, providing evidence for the suitability of the implemented, artificial C2-sink as a selection pressure for ALE. To identify the mechanisms responsible for the improved growth rate and biomass yield, we analyzed changes in the proteome that occurred during evolution. Remarkably, and in contrast to the local mutations normally observed in ALE programs, the effects of evolution may have stabilized a beneficial proteomic profile rather than causing a gain of function by up- or down-regulating specific proteins. This notion is supported by the change in gene copy numbers caused by the inversion of a large chromosomal segment.

### A flux split between the native ED and the synthetic PK pathways creates a metabolic imbalance in synthetic C2 auxotrophs

During batch fermentation on glucose, the evolved, PKT-dependent strain SCA3_PK-Tn_/pS438·*pta*^*Ec*^ secreted significant amounts of pyruvate and 2KG. Pyruvate accumulation as an overflow metabolite can be attributed to the removal of PDHc, and alternative entry reactions into the TCA cycle or backflow of pyruvate through gluconeogenesis would require additional ATP. Disproportionate production of 2KG, in contrast, can occur in *P. putida* in two scenarios: (i) genetic manipulations that cause bottlenecks in the supply of biomass constituents, or (ii) disrupting the intracellular redox balance (by decreasing the *supply* of NAD(P)H or increasing NADPH *demands via*, e.g., high levels of oxidative stress). In the first scenario, periplasmic dehydrogenases convert glucose into its acid forms faster than they can be utilized for biomass formation. This behavior resembles the phenomenon of acetate overflow production in glucose-grown *E. coli* [97]. In the second scenario, shortages within the pool of soluble electron carriers used to feed anabolic reactions and fuel the electron transport chain for energy production could limit bacterial growth. To maintain a high and homeostatic energy charge ([ATP] + 0.5 [ADP])/([ATP] + [ADP] + [AMP]), required to drive biosynthetic reactions [9,98,99], glucose can be utilized as an energy source without dissimilation. The electrons removed from glucose or gluconate during periplasmic oxidation are transferred to ubiquinones in the cytoplasmic membrane or FADH_2_, respectively [100,101]. Consequently, they can no longer contribute to replenishing the intracellular pool of NAD(P)H. Transport of 2KG across the cytoplasmic membrane has not been thoroughly studied in *P. putida*. It is assumed that the uptake is mediated by the major facilitator superfamily (MFS) transporter KguT [102] and was shown to occur against a concentration gradient [103], likely enabled *via* H^+^ symport. The utilization of 2KG requires the NADPH-dependent reduction of 2K6PG to 6PG [29].

In wild-type *P. putida* KT2440 growing on glucose, NADH was produced predominantly by PDHc, 2-oxoglutarate dehydrogenase (SucAB), and malate dehydrogenase (MDH). The reactions producing NADPH are, in order of their relative contribution, isocitrate dehydrogenase (Icd), glucose-6-P 1-dehydrogenase (Zwf), 6-phosphogluconate dehydrogenase (Gnd), and malic enzyme (MaeB). High flux through the reaction catalyzed by G3P dehydrogenase (GapDH) may make a variable contribution to the NADH/NADPH pool due to the simultaneous presence of isoforms with divergent substrate specificities. After inactivation of PDHc in strain SCA3_PK-Tn_/pS438·*pta*^*Ec*^, the predominant source of NADH is GapDH. NADPH is produced by Gnd and Zwf, and, to a lesser extent, MaeB and Icd. If glucose is oxidized to 2KG in the periplasm, 1 molecule of NADPH is required to reduce 2K6PG to 6PG before entering the ED pathway. Thus, the net yield of intracellular reducing equivalents obtained sums up to zero. If 6PG is fed into the PPP to supply substrates for Xfpk, the NADPH produced by Gnd can also only compensate for the initial consumption. In order to produce more soluble electron carriers, a significant metabolite flux must be delivered into the TCA cycle. This would require either a higher flux fraction through the two PK reactions or an increased contribution of anaplerotic reactions. Thereby, increased production of NAD(P)H would allow the use of 2KG, which further increases the biomass yield on glucose.

### A synthetic PK-ED pathway can be exploited for the efficient production of chemicals

The main goal of this study was to establish synthetic C2-auxotrophy as a selection scheme for synthetic metabolism through evolutionary engineering. While a significant growth optimization could be achieved, the formation of biomass was limited by insufficient production of energy-generating reducing equivalents. While 2KG overflow production needs to be addressed to increase the flux of carbon into the cytoplasm, the simultaneous activities of Edd/Eda and Xfpk provide ample substrate amounts for bioproduction with pathways that use pyruvate as the entry metabolite [e.g., isobutanol [78], D-lactic acid [104,105], or 2,3-butanediol [106]]. With these prospects, this study provides not only an example of the usefulness of carefully designed selection strains for evolutionary engineering but also reveals unconventional metabolic engineering strategies for microbial production.

## Notes

### Competing Interest Statement

The authors have declared no competing interest.

## REFERENCES

1. Shi, L. and Tu, B.P. (2015) Acetyl-CoA and the regulation of metabolism: mechanisms and consequences. Curr. Opin. Cell Biol. 33, 125–131

2. Lu, X. et al. (2019) Constructing a synthetic pathway for acetyl-coenzyme A from one-carbon through enzyme design. Nat. Commun. 10, 1378

3. Vadali, R.V. et al. (2004) Cofactor engineering of intracellular CoA/acetyl-CoA and its effect on metabolic flux redistribution in Escherichia coli. Metab. Eng. 6, 133–139

4. Yang, D. et al. (2020) Metabolic engineering of Escherichia coli for natural product biosynthesis. Trends Biotechnol. 38, 745–765

5. Mezzina, M.P. et al. (2021) Engineering native and synthetic pathways in Pseudomonas putida for the production of tailored polyhydroxyalkanoates. Biotechnol. J. 16, 2000165

6. Zhu, L. et al. (2022) Strategies for optimizing acetyl-CoA formation from glucose in bacteria. Trends Biotechnol. 40, 149–165

7. Weimer, A. et al. (2020) Industrial biotechnology of Pseudomonas putida: Advances and prospects. Appl. Microbiol. Biotechnol. 104, 7745–7766

8. Poblete-Castro, I. et al. (2012) Industrial biotechnology of Pseudomonas putida and related species. Appl. Microbiol. Biotechnol. 93, 2279–2290

9. Ebert, B.E. et al. (2011) Response of Pseudomonas putida KT2440 to increased NADH and ATP demand. Appl. Environ. Microbiol. 77, 6597–6605

10. Belda, E. et al. (2016) The revisited genome of Pseudomonas putida KT2440 enlightens its value as a robust metabolic chassis. Environ. Microbiol. 18, 3403–3424

11. Sánchez-Pascuala, A. et al. (2019) Functional implementation of a linear glycolysis for sugar catabolism in Pseudomonas putida. Metab. Eng. 54, 200–211

12. Kozaeva, E. et al. (2021) Model-guided dynamic control of essential metabolic nodes boosts acetyl-coenzyme A–dependent bioproduction in rewired Pseudomonas putida. Metab. Eng. 67, 373–386

13. Chohnan, S. et al. (1997) Changes in the size and composition of intracellular pools of nonesterified coenzyme A and coenzyme A thioesters in aerobic and facultatively anaerobic bacteria. Appl. Environ. Microbiol. 63, 553–560

14. Chang, D.E. et al. (1999) Acetate metabolism in a pta mutant of Escherichia coli W3110: Importance of maintaining acetyl coenzyme A flux for growth and survival. J. Bacteriol. 181, 6656–6663

15. Bar-Even, A. et al. (2012) Rethinking glycolysis: on the biochemical logic of metabolic pathways. Nat. Chem. Biol. 8, 509–517

16. Chavarría, M. et al. (2013) The Entner-Doudoroff pathway empowers Pseudomonas putida KT2440 with a high tolerance to oxidative stress. Environ. Microbiol. 15, 1772–1785

17. Nikel, P.I. et al. (2016) From dirt to industrial applications: Pseudomonas putida as a Synthetic Biology chassis for hosting harsh biochemical reactions. Curr. Opin. Chem. Biol. 34, 20–29

18. Patel, M.S. et al. (2014) The pyruvate dehydrogenase complexes: structure-based function and regulation. J. Biol. Chem. 289, 16615–16623

19. Scardovi, V. and Trovatelli, L.D. (1965) The fructose-6-phosphate shunt as peculiar pattern of hexose degradation in the genus Bifidobacterium. Ann. Microbiol. Enz. 15, 19–29

20. Krüsemann, J.L. et al. (2018) Artificial pathway emergence in central metabolism from three recursive phosphoketolase reactions. FEBS J. 285, 4367–4377

21. Tittmann, K. (2014) Sweet siblings with different faces: the mechanisms of FBP and F6P aldolase, transaldolase, transketolase and phosphoketolase revisited in light of recent structural data. Bioorg. Chem. 57, 263–280

22. Bogorad, I.W. et al. (2013) Synthetic non-oxidative glycolysis enables complete carbon conservation. Nature 502, 693–697

23. Henard, C.A. et al. (2015) Phosphoketolase pathway engineering for carbon-efficient biocatalysis. Curr. Opin. Biotechnol. 36, 183–188

24. Ku, J.T. et al. (2020) Metabolic engineering design strategies for increasing acetyl-CoA flux. Metabolites 10, 166

25. Hellgren, J. et al. (2020) Promiscuous phosphoketolase and metabolic rewiring enables novel non-oxidative glycolysis in yeast for high-yield production of acetyl-CoA derived products. Metab Eng 62, 150–160

26. Qin, N. et al. (2020) Rewiring central carbon metabolism ensures increased provision of acetyl-CoA and NADPH required for 3-OH-propionic acid production. ACS Synth. Biol. 9, 3236–3244

27. Wang, Q. et al. (2019) Engineering an in vivo EP-bifido pathway in Escherichia coli for high-yield acetyl-CoA generation with low CO_2_ emission. Metab. Eng. 51, 79–87

28. Nikel, P.I. and de Lorenzo, V. (2013) Engineering an anaerobic metabolic regime in Pseudomonas putida KT2440 for the anoxic biodegradation of 1,3-dichloroprop-1-ene. Metab. Eng. 15, 98–112

29. Nikel, P.I. et al. (2015) Pseudomonas putida KT2440 strain metabolizes glucose through a cycle formed by enzymes of the Entner-Doudoroff, Embden-Meyerhof-Parnas, and pentose phosphate pathways. J. Biol. Chem. 290, 25920–25932

30. Nikel, P.I. et al. (2021) Reconfiguration of metabolic fluxes in Pseudomonas putida as a response to sub-lethal oxidative stress. ISME J. 15, 1751–1766

31. Fernández-Cabezón, L. et al. (2019) Evolutionary approaches for engineering industrially-relevant phenotypes in bacterial cell factories. Biotechnol. J. 14, 1800439

32. Sandberg, T.E. et al. (2019) The emergence of adaptive laboratory evolution as an efficient tool for biological discovery and industrial biotechnology. Metab. Eng. 56, 1–16

33. Orsi, E. et al. (2021) Growth-coupled selection of synthetic modules to accelerate cell factory development. Nat. Commun. 12, 5295

34. Cros, A. et al. (2022) Synthetic metabolism for biohalogenation. Curr. Opin. Biotechnol. 74, 180–193

35. Hartmans, S. et al. (1989) Metabolism of styrene oxide and 2-phenylethanol in the styrene-degrading Xanthobacter strain 124X. Appl. Environ. Microbiol. 55, 2850–2855

36. Platt, R. et al. (2000) Genetic system for reversible integration of DNA constructs and lacZ gene fusions into the Escherichia coli chromosome. Plasmid 43, 12–23

37. Worsey, M.J. and Williams, P.A. (1975) Metabolism of toluene and xylenes by Pseudomonas putida (arvilla) mt-2: evidence for a new function of the TOL plasmid. J. Bacteriol. 124, 7–13

38. Bagdasarian, M. et al. (1981) Specific purpose plasmid cloning vectors. II. Broad host range, high copy number, RSF1010-derived vectors, and a host-vector system for gene cloning in Pseudomonas. Gene 16, 237–247

39. Arias-Barrau, E. et al. (2004) The homogentisate pathway: a central catabolic pathway involved in the degradation of L-phenylalanine, L-tyrosine, and 3-hydroxyphenylacetate in Pseudomonas putida. J. Bacteriol. 186, 5062–5077

40. Volke, D.C. et al. (2020) Synthetic control of plasmid replication enables target- and self-curing of vectors and expedites genome engineering of Pseudomonas putida. Metab. Eng. Commun. 10, e00126

41. Cavaleiro, A.M. et al. (2015) Accurate DNA assembly and genome engineering with optimized uracil excision cloning. ACS Synth. Biol. 4, 1042–1046

42. Volke, D.C. and Nikel, P.I. (2018) Getting bacteria in shape: Synthetic morphology approaches for the design of efficient microbial cell factories. Adv. Biosyst. 2, 1800111

43. Volke, D.C. et al. (2020) Physical decoupling of XylS/Pm regulatory elements and conditional proteolysis enable precise control of gene expression in Pseudomonas putida. Microb. Biotechnol. 13, 222–232

44. Genee, H.J. et al. (2015) Software-supported USER cloning strategies for site-directed mutagenesis and DNA assembly. ACS Synth. Biol. 4, 342–349

45. Wirth, N.T. et al. (2020) Accelerated genome engineering of Pseudomonas putida by I- SceI-mediated recombination and CRISPR-Cas9 counterselection. Microb. Biotechnol. 13, 233–249

46. Calero, P. and Nikel, P.I. (2019) Chasing bacterial chassis for metabolic engineering: A perspective review from classical to non-traditional microorganisms. Microb. Biotechnol. 12, 98–124

47. Calero, P. et al. (2020) A fluoride-responsive genetic circuit enables in vivo biofluorination in engineered Pseudomonas putida. Nat. Commun. 11, 5045

48. Batianis, C. et al. (2020) An expanded CRISPRi toolbox for tunable control of gene expression in Pseudomonas putida. Microb. Biotechnol. 13, 368–385

49. Ruiz, J.A. et al. (2006) dye (arc) Mutants: insights into an unexplained phenotype and its suppression by the synthesis of poly(3-hydroxybutyrate) in Escherichia coli recombinants. FEMS Microbiol. Lett. 258, 55–60

50. Volke, D.C. et al. (2022) Modular (de)construction of complex bacterial phenotypes by CRISPR/nCas9-assisted, multiplex cytidine base-editing. Nat. Commun. 13, 3026

51. Wirth, N.T. and Nikel, P.I. (2021) Combinatorial pathway balancing provides biosynthetic access to 2-fluoro-cis,cis-muconate in engineered Pseudomonas putida. Chem Catal. 1, 1234–1259

52. Choi, K.H. et al. (2006) A 10-min method for preparation of highly electrocompetent Pseudomonas aeruginosa cells: application for DNA fragment transfer between chromosomes and plasmid transformation. J. Microbiol. Methods 64, 391–397

53. Choi, K.H. et al. (2005) A Tn7-based broad-range bacterial cloning and expression system. Nat. Methods 2, 443–448

54. Choi, K.H. and Schweizer, H.P. (2006) Mini-Tn7 insertion in bacteria with single attTn7 sites: example Pseudomonas aeruginosa. Nat. Protoc. 1, 153–161

55. Nogales, J. et al. (2020) High-quality genome-scale metabolic modelling of Pseudomonas putida highlights its broad metabolic capabilities. Environ. Microbiol. 22, 255–269

56. Orth, J.D. et al. (2010) What is flux balance analysis? Nat. Biotechnol. 28, 245–248

57. Ebrahim, A. et al. (2013) COBRApy: COnstraints-Based Reconstruction and Analysis for Python. BMC Syst. Biol. 7, 74

58. Nikel, P.I. et al. (2008) Escherichia coli arcA mutants: metabolic profile characterization of microaerobic cultures using glycerol as a carbon source. J. Mol. Microbiol. Biotechnol. 15, 48–54

59. Nikel, P.I. et al. (2010) Ethanol synthesis from glycerol by Escherichia coli redox mutants expressing adhE from Leuconostoc mesenteroides. J. Appl. Microbiol. 109, 492–504

60. Nikel, P.I. et al. (2009) Metabolic flux analysis of Escherichia coli creB and arcA mutants reveals shared control of carbon catabolism under microaerobic growth conditions. J. Bacteriol. 191, 5538–5548

61. del Castillo, T. et al. (2007) Convergent peripheral pathways catalyze initial glucose catabolism in Pseudomonas putida: genomic and flux analysis. J. Bacteriol. 189, 5142–5152

62. Rennig, M. et al. (2019) Industrializing a bacterial strain for L-serine production through translation initiation optimization. ACS Synth. Biol. 8, 2347–2358

63. Huber, W. et al. (2002) Variance stabilization applied to microarray data calibration and to the quantification of differential expression. Bioinformatics 18 Suppl 1, S96–104

64. Gatto, L. et al. (2021) MSnbase, efficient and elegant R-based processing and visualization of raw mass spectrometry data. J. Proteome Res. 20, 1063–1069

65. Ritchie, M.E. et al. (2015) limma powers differential expression analyses for RNA-sequencing and microarray studies. Nucleic Acids Res. 43, e47–e47

66. Strimmer, K. (2008) fdrtool: a versatile R package for estimating local and tail area-based false discovery rates. Bioinformatics 24, 1461–1462

67. Qiu, Y.Q. (2013) KEGG Pathway Database. In Encyclopedia of Systems Biology (Dubitzky, W. et al., eds), pp. 1068–1069, Springer New York

68. Wu, T. et al. (2021) clusterProfiler 4.0: A universal enrichment tool for interpreting omics data. Innovation 2, 100141

69. van Duuren, J.B. et al. (2013) Reconciling in vivo and in silico key biological parameters of Pseudomonas putida KT2440 during growth on glucose under carbon-limited condition. BMC Biotechnol. 13, 93

70. Kohlstedt, M. and Wittmann, C. (2019) GC-MS-based ^13^C metabolic flux analysis resolves the parallel and cyclic glucose metabolism of Pseudomonas putida KT2440 and Pseudomonas aeruginosa PAO1. Metab. Eng. 54, 35–53

71. Young, J.D. (2014) INCA: a computational platform for isotopically non-stationary metabolic flux analysis. Bioinformatics 30, 1333–1335

72. Lipmann, F. and Tuttle, L.C. (1945) A specific micromethod for the determination of acyl phosphates. J. Biol. Chem. 159, 21–28

73. He, F. (2011) Bradford protein assay. Bio-Protocol 1, e45

74. Fernández-Cabezón, L. et al. (2021) Spatiotemporal manipulation of the mismatch repair system of Pseudomonas putida accelerates phenotype emergence. ACS Synth. Biol. 10, 1214–1226

75. Swain, P.S. et al. (2016) Inferring time derivatives including cell growth rates using Gaussian processes. Nat. Commun. 7, 13766

76. Karp, P.D. et al. (2019) The BioCyc collection of microbial genomes and metabolic pathways. Brief. Bioinform. 20, 1085–1093

77. Volke, D.C. et al. (2021) Cofactor specificity of glucose-6-phosphate dehydrogenase isozymes in Pseudomonas putida reveals a general principle underlying glycolytic strategies in bacteria. mSystems 6, e00014–00021

78. Nitschel, R. et al. (2020) Engineering Pseudomonas putida KT2440 for the production of isobutanol. Eng. Life Sci. 20, 148–159

79. Udaondo, Z. et al. (2018) Regulation of carbohydrate degradation pathways in Pseudomonas involves a versatile set of transcriptional regulators. Microb. Biotechnol. 11, 442–454

80. Bergman, A. et al. (2016) Functional expression and evaluation of heterologous phosphoketolases in Saccharomyces cerevisiae. AMB Express 6, 115

81. Zobel, S. et al. (2015) Tn7-Based device for calibrated heterologous gene expression in Pseudomonas putida. ACS Synth. Biol. 4, 1341–1351

82. Mutalik, V.K. et al. (2013) Precise and reliable gene expression via standard transcription and translation initiation elements. Nat. Methods 10, 354–360

83. Wu, X. et al. (2011) Comparative genomics and functional analysis of niche-specific adaptation in Pseudomonas putida. FEMS Microbiol. Rev. 35, 299–323

84. Martínez-García, E. et al. (2014) Freeing Pseudomonas putida KT2440 of its proviral load strengthens endurance to environmental stresses. Environ. Microbiol. 17, 76–90

85. Martínez-García, E. et al. (2014) Pseudomonas 2.0: genetic upgrading of P. putida KT2440 as an enhanced host for heterologous gene expression. Microb. Cell Fact. 13, 159

86. Ilves, H. et al. (2001) Involvement of sS in starvation-induced transposition of Pseudomonas putida transposon Tn4652. J. Bacteriol. 183, 5445–5448

87. Hõrak, R. and Kivisaar, M. (1998) Expression of the transposase gene tnpA of Tn4652 is positively affected by integration host factor. J. Bacteriol. 180, 2822–2829

88. Kivistik, P.A. et al. (2007) Target site selection of Pseudomonas putida transposon Tn4652. J. Bacteriol. 189, 3918–3921

89. Martínez-García, E. et al. (2014) The metabolic cost of flagellar motion in Pseudomonas putida KT2440. Environ. Microbiol. 16, 291–303

90. Schuiki, I. and Daum, G. (2009) Phosphatidylserine decarboxylases, key enzymes of lipid metabolism. IUBMB Life 61, 151–162

91. Cho, G. et al. (2021) Structural insights into phosphatidylethanolamine formation in bacterial membrane biogenesis. Sci. Rep. 11, 5785

92. McVey, A.C. et al. (2017) Structural and functional characterization of malate synthase G from opportunistic pathogen Pseudomonas aeruginosa. Biochemistry 56, 5539–5549

93. Sievers, M. et al. (1997) Purification and properties of citrate synthase from Acetobacter europaeus. FEMS Microbiol. Lett. 146, 53–58

94. Klinke, S. et al. (2000) Inactivation of isocitrate lyase leads to increased production of medium-chain-length poly(3-hydroxyalkanoates) in Pseudomonas putida. Appl. Environ. Microbiol. 66, 909–913

95. Brock, M. et al. (2000) Methylcitrate synthase from Aspergillus nidulans: implications for propionate as an antifungal agent. Mol. Microbiol. 35, 961–973

96. Maerker, C. et al. (2005) Methylcitrate synthase from Aspergillus fumigatus. FEBS J. 272, 3615–3630

97. Valgepea, K. et al. (2010) Systems biology approach reveals that overflow metabolism of acetate in Escherichia coli is triggered by carbon catabolite repression of acetyl-CoA synthetase. BMC Syst. Biol. 4, 166

98. Andersen, K.B. and von Meyenburg, K. (1977) Charges of nicotinamide adenine nucleotides and adenylate energy charge as regulatory parameters of the metabolism in Escherichia coli. J. Biol. Chem. 252, 4151–4156

99. Atkinson, D.E. (1968) The energy charge of the adenylate pool as a regulatory parameter. Interaction with feedback modifiers. Biochemistry 7, 4030–4034

100. An, R. and Moe, L.A. (2016) Regulation of pyrroloquinoline quinone-dependent glucose dehydrogenase activity in the model rhizosphere-dwelling bacterium Pseudomonas putida KT2440. Appl. Environ. Microbiol. 82, 4955–4964

101. Matsushita, K. et al. (1979) Membrane-bound D-gluconate dehydrogenase from Pseudomonas aeruginosa. Its kinetic properties and a reconstitution of gluconate oxidase. J. Biochem. 86, 249–256

102. Daddaoua, A. et al. (2010) Compartmentalized glucose metabolism in Pseudomonas putida is controlled by the PtxS repressor. J. Bacteriol. 192, 4357–4366

103. Torrontegui, D. et al. (1976) The uptake of 2-ketogluconate by Pseudomonas putida. Arch. Microbiol. 110, 43–48

104. Ishida, N. et al. (2006) D-Lactic acid production by metabolically engineered Saccharomyces cerevisiae. J. Biosci. Bioeng. 101, 172–177

105. Akita, H. et al. (2017) Production of D-lactate using a pyruvate-producing Escherichia coli strain. Biosci. Biotechnol. Biochem. 81, 1452–1455

106. Song, C.W. et al. (2019) Microbial production of 2,3-butanediol for industrial applications. J. Ind. Microbiol. Biotechnol. 46, 1583–1601

