## Supplemental Data 1 for "A synthetic C2 auxotroph of *Pseudomonas putida* for evolutionary engineering of alternative sugar catabolic routes"

### Tables

**Table S1. Oligonucleotides used in this study.** Sequences used for the construction of the same plasmid via *USER* cloning are shaded in the same tone.

| Name | Sequence (5'→3') | Application |
|---|---|---|
| pSNW-*USER*_F | AGTCGACCUGCAGGCATGCAAGCTTCT | Linearization of suicide vector pSNW for HA insertion |
| pSNW- *USER* _R | AGGATCUAGAGGATCCCCGGGTACCG | Linearization of suicide vector pSNW for HA insertion |
| Seq-pSNW_F | TGTAAAACGACGGCCAGT | Amplification and sequencing of the insert region on pSNW |
| Seq-pSNW_R | ATGACCATGATTACGCCGG | Amplification and sequencing of the insert region on pSNW |
| *aceEF*_HA1_F | AGATCCUCGAAGACTCGCTTGAAGAGG | Amplification of HA1 for the deletion of *aceEF* |
| *aceEF*_HA1_R | AGCCATGUAAGCCAGCACACTGC | Amplification of HA1 for the deletion of *aceEF* |
| *aceEF*_HA2_F | ACATGGCUTGCTCCAGGG | Amplification of HA2 for the deletion of *aceEF* |
| *aceEF*_HA2_R | AGGTCGACUCGATGAACTGCTGGTTGCG | Amplification of HA2 for the deletion of *aceEF* |
| *aceEF*_g-check_F | GTTTGGCTGGAGATTTTGGG | Genotyping for the chromosomal *aceEF* region via colony PCR |
| *aceEF*_g-check_R | CCTTGATCGGCGTGAAATAG | Genotyping for the chromosomal *aceEF* region via colony PCR |
| *aceEF*_F | GCGCGCTCATTCGCGTACCTGACAT | sgRNA for *aceEF* deletion |
| *aceEF*_R | AAACATGTCAGGTACGCGAATGAGC | sgRNA for *aceEF* deletion |
| Seq-LP_F | ACCAACTTTTCCGCTTTGCAC | Amplification and sequencing of the chromosomal landing pad |
| Seq-LP_R | CGAAAGACTGGGCCTTTCGT | Amplification and sequencing of the chromosomal landing pad |
| *gltA*_HA1_F | AGATCCUCTAGCATAAAGCATGATGGC | Amplification of HA1 for the deletion of *gltA* |
| *gltA*_HA1_R | ACATAGCUTACATGTGGCCTCCTATT | Amplification of HA1 for the deletion of *gltA* |
| *gltA*_HA2_F | AGCTATGUAAGCTGCAGTAGCCGAAC | Amplification of HA2 for the deletion of *gltA* |
| *gltA*_HA2_R | AGGTCGACUGTTGGCGGAGGTTCTCAAG | Amplification of HA2 for the deletion of *gltA* |
| *gltA*_g-check_F | TCCCTCTATAGTGGTGCGGG | Genotyping for the chromosomal *gltA* region via colony PCR |
| *gltA*_g-check_R | TACCTGCCGGAAGAGAAGGG | Genotyping for the chromosomal *gltA* region via colony PCR |
| *acoABC*_HA1_F | AGATCCUGTGCCTGAACAACAGCTGG | Amplification of HA1 for the deletion of *acoABC* |
| *acoABC*_HA1_R | ACATCTUGTTGTTCTCCGGGG | Amplification of HA1 for the deletion of *acoABC* |
| *acoABC*_HA2_F | AAGATGUAAGCCCTTTTCGTGAGCCT | Amplification of HA2 for the deletion of *acoABC* |
| *acoABC*_HA2_R | AGGTCGACUCCACTACAACGGTTTGCCC | Amplification of HA2 for the deletion of *acoABC* |
| *acoABC*_g-check_F | CGAATTTGCCGAGCATGACA | Genotyping for the chromosomal *acoABC* region via colony PCR |
| *acoABC*_g-check_R | CACCAGTCCACCTTGATCTG | Genotyping for the chromosomal *acoABC* region via colony PCR |

| | | |
|---|---|---|
| *amaC-dpkA*_HA1_F | AGATCCUATCGGCAGGAACAGTGTCC | Amplification of HA1 for the deletion of *amaC-dpkA* |
| *amaC-dpkA*_HA1_R | ACCGTTGTGUAACTCGCCAACGGGCCAAC | Amplification of HA1 for the deletion of *amaC-dpkA* |
| *amaC-dpkA*_HA2_F | ACACAACGGUGCTGGTGGAAG | Amplification of HA2 for the deletion of *amaC-dpkA* |
| *amaC-dpkA*_HA2_R | AGGTCGACUCTCTGGCGGTATTCGTGTACG | Amplification of HA2 for the deletion of *amaC-dpkA* |
| *amaC-dpkA*_g-check_F | CACGGAAAGGTTGCACTTGC | Genotyping for the chromosomal *amaC-dpkA* region via colony PCR |
| *amaC-dpkA*_g-check_R | GATCAGTTGGCTCATGCTGC | Genotyping for the chromosomal *amaC-dpkA* region via colony PCR |
| *bkdAA*_HA1_F | AGATCCUGGTCATCAGGTAGCACTCC | Amplification of HA1 for the deletion of *bkdAA* |
| *bkdAA*_HA1_R | ATCTCACAUCATGCTTTTTACGCTCGCTCGG | Amplification of HA1 for the deletion of *bkdAA* |
| *bkdAA*_HA2_F | ATGTGAGAUGAACGACCACAACAACAG | Amplification of HA2 for the deletion of *bkdAA* |
| *bkdAA*_HA2_R | AGGTCGACTCCAGGAAGATTACCGGGTCG | Amplification of HA2 for the deletion of *bkdAA* |
| *bkdAA*_g-check_F | CCTGCTCAGCAAAGGCAATC | Genotyping for the chromosomal *bkdAA* region via colony PCR |
| *bkdAA*_g-check_R | ATGGTCATGGTAGTGGTGGC | Genotyping for the chromosomal *bkdAA* region via colony PCR |
| *crc*_HA1_F | ACCTCCAGUGATGATCTGCATGACCTCACGAATGG | Amplification of HA1 for the deletion of *crc* |
| *crc*_HA1_R | ACATAAAUGGCCCCATAAATCTCGTG | Amplification of HA1 for the deletion of *crc* |
| *crc*_HA2_F | ATTTATGUAAAAGGCCATTGGGGCTGCAT | Amplification of HA2 for the deletion of *crc* |
| *crc*_HA2_R | AGCTCTTGAUAACGCCATGCTCGCTTTGGC | Amplification of HA2 for the deletion of *crc* |
| *crc*_g-check_F | ACAGCACATCGAACGGGATC | Genotyping for the chromosomal *crc* region via colony PCR |
| *crc*_g-check_R | TGCTATGGCACCTCAAGCG | Genotyping for the chromosomal *crc* region via colony PCR |
| *davBA*_HA1_F | AGATCCUCTGGGCCAGAGTGTGATAGG | Amplification of HA1 for the deletion of *davBA* |
| *davBA*_HA1_R | ACATGAAAUGACCTTGCCAGAGAG | Amplification of HA1 for the deletion of *davBA* |
| *davBA*_HA2_F | ATTTCATGUGACAGGGGCCGCTATGC | Amplification of HA2 for the deletion of *davBA* |
| *davBA*_HA2_R | AGGTCGACUTGGATCACTTCGCGTTCACG | Amplification of HA2 for the deletion of *davBA* |
| *davBA*_g-check_F | AGCGTATGTCGGGATCAAGG | Genotyping for the chromosomal *davBA* region via colony PCR |
| *davBA*_g-check_R | GGGTACCGAGGAAGAACAGG | Genotyping for the chromosomal *davBA* region via colony PCR |
| *eutBC*_HA1_F | AGATCCUAGATCTTCGCCCTGTCCC | Amplification of HA1 for the deletion of *eutBC* |
| *eutBC*_HA1_R | ATTTTTTACAUACAGAATCTCCAGAGCC | Amplification of HA1 for the deletion of *eutBC* |
| *eutBC*_HA2_F | ATGTAAAAAAUCTGTCACGGAAGGC | Amplification of HA2 for the deletion of *eutBC* |
| *eutBC*_HA2_R | AGGTCGACUACCCGATGGATTCGTAAGTC | Amplification of HA2 for the deletion of *eutBC* |

| | | |
|---|---|---|
| *eutBC*_g-check_F | CTACCCTGTCTTACGTGCTG | Genotyping for the chromosomal *eutBC* region via colony PCR |
| *eutBC*_g-check_R | TAAATCACCGATTCGTCCCG | Genotyping for the chromosomal *eutBC* region via colony PCR |
| *gabT*_HA1_F | AGATCCUCAAGGTAGGCTGACAGG | Amplification of HA1 for the deletion of *gabT* |
| *gabT*_HA1_R | ATGTGAUGTGACGCGCTTCAGAA | Amplification of HA1 for the deletion of *gabT* |
| *gabT*_HA2_F | ATCACAUAATGCCCTCATTGCGCC | Amplification of HA2 for the deletion of *gabT* |
| *gabT*_HA2_R | AGGTCGACUGTCCAGGAACACATCGAGG | Amplification of HA2 for the deletion of *gabT* |
| *gabT*_g-check_F | CTCATCGCCAACTACTCCTG | Genotyping for the chromosomal *gabT* region via colony PCR |
| *gabT*_g-check_R | CCTGATCTCCAACGAAGTGG | Genotyping for the chromosomal *gabT* region via colony PCR |
| *glcB*_HA1_F | AGATCCUACTGGGCAATACGTTCGACC | Amplification of HA1 for the deletion of *glcB* |
| *glcB*_HA1_R | ACATTGCUTGCCTCACTCTGC | Amplification of HA1 for the deletion of *glcB* |
| *glcB*_HA2_F | AGCAATGUAAGCAGCATCTGCTGCCAG | Amplification of HA2 for the deletion of *glcB* |
| *glcB*_HA2_R | AGGTCGACUGCGGTCCTAATGCTGAAGC | Amplification of HA2 for the deletion of *glcB* |
| *glcB*_g-check_F | GTTTGGATGTCTGCCAAGGC | Genotyping for the chromosomal *glcB* region via colony PCR |
| *glcB*_g-check_R | TCCTGTACCTGAAAAGCCGG | Genotyping for the chromosomal *glcB* region via colony PCR |
| *hmgABC*_HA1_F | AGATCCUAACCGATCTGGGTCACCAC | Amplification of HA1 for the deletion of *hmgABC* |
| *hmgABC*_HA1_R | AGCCATCGAUGCCTCCGGATTTTG | Amplification of HA1 for the deletion of *hmgABC* |
| *hmgABC*_HA2_F | ATCGATGGCUCAGTGAAGTGAACGGTTC | Amplification of HA2 for the deletion of *hmgABC* |
| *hmgABC*_HA2_R | AGGTCGACUGGCCATCCTCGCTTTG | Amplification of HA2 for the deletion of *hmgABC* |
| *hmgABC*_g-check_F | ATCCTGTTCGGCAAAACCAC | Genotyping for the chromosomal *hmgABC* region via colony PCR |
| *hmgABC*_g-check_R | GGTGAGCGACTACATGACTC | Genotyping for the chromosomal *hmgABC* region via colony PCR |
| *ltaE*_HA1_F | AGATCCUACCTTGTCTACGTGCTCG | Amplification of HA1 for the deletion of *ltaE* |
| *ltaE*_HA1_R | ATGGCACGGUCCTGTGAAC | Amplification of HA1 for the deletion of *ltaE* |
| *ltaE*_HA2_F | ACCGTGCCAUGTGAGCGAGAGCCGGGGCC | Amplification of HA2 for the deletion of *ltaE* |
| *ltaE*_HA2_R | AGGTCGACUCGGTCGATGAAGTGCTC | Amplification of HA2 for the deletion of *ltaE* |
| *ltaE*_g-check_F | TGGCGATGTTATGACTGGTG | Genotyping for the chromosomal *ltaE* region via colony PCR |
| *ltaE*_g-check_R | TAGAGCACATCAGCGACATG | Genotyping for the chromosomal *ltaE* region via colony PCR |
| *mmsA1*_HA1_F | AGATCCUAGACCTTCATGAACCAGCCG | Amplification of HA1 for the deletion of *mmsA1* |
| *mmsA1*_HA1_R | ACTTCCCTUACATGCAAAACTCCAGATAAACAAAGG | Amplification of HA1 for the deletion of *mmsA1* |
| *mmsA1*_HA2_F | AAGGGAAGUAGATGAAGCGGG | Amplification of HA2 for the deletion of *mmsA1* |
| *mmsA1*_HA2_R | AGGTCGACUTCCTCAGAGCCTGTCAAGC | Amplification of HA2 for the deletion of *mmsA1* |
| *mmsA1*_g-check_F | CTGTTGGCGAAACCCTGAAC | Genotyping for the chromosomal *mmsA1* region via colony PCR |
| *mmsA1*_g-check_R | GCTTCAAACACTTGGCTCGC | Genotyping for the chromosomal *mmsA1* region via colony PCR |

| | | |
|---|---|---|
| *mmsA2*_HA1_F | AGATCCUCAGGATGTCCACCGAAATTGC | Amplification of HA1 for the deletion of *mmsA2* |
| *mmsA2*_HA1_R | ACTCCTTACAUCTGCGTTCTCCTTGGAATTG | Amplification of HA1 for the deletion of *mmsA2* |
| *mmsA2*_HA2_F | ATGTAAGGAGUACGTCATGCGTATCG | Amplification of HA2 for the deletion of *mmsA2* |
| *mmsA2*_HA2_R | AGGTCGACUGCAGGTTGTTGCAGATCTTGG | Amplification of HA2 for the deletion of *mmsA2* |
| *mmsA2*_g-check_F | GACAGGCTGATGAAATGCGG | Genotyping for the chromosomal *mmsA2* region via colony PCR |
| *mmsA2*_g-check_R | TCAGGTCGAACAGGTTCAGC | Genotyping for the chromosomal *mmsA2* region via colony PCR |
| *paaEDCBA*_HA1_F | AGATCCUGTGAATGACGGCGCTTGC | Amplification of HA1 for the deletion of *paaEDCBA* |
| *paaEDCBA*_HA1_R | ACATGGCUTCACTCGAATTGTTC | Amplification of HA1 for the deletion of *paaEDCBA* |
| *paaEDCBA*_HA2_F | AGCCATGUAGACCGAGTCGGCTGCTTC | Amplification of HA2 for the deletion of *paaEDCBA* |
| *paaEDCBA*_HA2_R | AGGTCGACUCCAGCAGGGTGAAGCAGAC | Amplification of HA2 for the deletion of *paaEDCBA* |
| *paaEDCBA*_g-check_F | GTTACCGAACTGAACGCAGC | Genotyping for the chromosomal *paaEDCBA* region via colony PCR |
| *paaEDCBA*_g-check_R | GCTTGACCTTGTCGTTGAGC | Genotyping for the chromosomal *paaEDCBA* region via colony PCR |
| *pcaBDC*_HA1_F | AGATCCUATGCAGAAGTACCTGGTCAAC | Amplification of HA1 for the deletion of *pcaBDC* |
| *pcaBDC*_HA1_R | ACGGTTTACAUGCAGCGTCCTTAATCATC | Amplification of HA1 for the deletion of *pcaBDC* |
| *pcaBDC*_HA2_F | ATGTAAACCGUGATCACGGGCAG | Amplification of HA2 for the deletion of *pcaBDC* |
| *pcaBDC*_HA2_R | AGGTCGACUGAGCCAATACGCCAACTACC | Amplification of HA2 for the deletion of *pcaBDC* |
| *pcaBDC*_g-check_F | AAACCCTGGGGATGGAAAAC | Genotyping for the chromosomal *pcaBDC* region via colony PCR |
| *pcaBDC*_g-check_R | TCTGTGTGGTTCTACTTGCG | Genotyping for the chromosomal *pcaBDC* region via colony PCR |
| *prpC*_HA1_F | AGATCCUCCGTATCAAAGCTGCCGTC | Amplification of HA1 for the deletion of *prpC* |
| *prpC*_HA1_R | ATTCACAUGGTGTTTCTCCTTTCTTGAA | Amplification of HA1 for the deletion of *prpC* |
| *prpC*_HA2_F | ATGTGAAUAGCTATAGAGAGGCAACC | Amplification of HA2 for the deletion of *prpC* |
| *prpC*_HA2_R | AGGTCGACUGTGGTCGACGATCAGTTGC | Amplification of HA2 for the deletion of *prpC* |
| *prpC*_g-check_F | ACGTGTCGTTGGTGCTGTAC | Genotyping for the chromosomal *prpC* region via colony PCR |
| *prpC*_g-check_R | GTATGGCAAGCCGTCGTAGG | Genotyping for the chromosomal *prpC* region via colony PCR |
| *xfpk* _F | ATGACGAGUCCTGTTATTGGCA | Amplification of *xfpk* for integration into pGNW2·LPR |
| *xfpk* _R | ATCTCACUCGTTATCGCCAGCG | Amplification of *xfpk* for integration into pGNW2·LPR |
| pGNW2·LP_F | AGTGAGAUAGTGCTAGTGTAGATCGC | Linearization of pGNW2·LPR to replace *rfp* with *xfpk* |
| pGNW2·LP_R | ACTCGTCAUTAGAAAACCTCCTTAGCATGATTA | Linearization of pGNW2·LPR to replace *rfp* with *xfpk* |

| | | |
|---|---|---|
| *pta*$^{Ec}$_F | ATCGTTTCUAATGATTATTGAACGTTGTCGTGAA | Amplification of *pta* from the chromosome of *E. coli* DH5α |
| *pta*$^{Ec}$_R | AGTCATUGGGACCGTTCATTCAACC | Amplification of *pta* from the chromosome of *E. coli* DH5α |
| pS438·*BCD10*_F | AATGACUGGGAAAACCCTGGC | Linearization of pS438·*BCD10* for the integration of *pta*$^{Ec}$ |
| pS438·*BCD10*_R | AGAAACGAUCCTCCGCATGATTAA GATGTTTCAGTACG | Linearization of pS438·*BCD10* for the integration of *pta*$^{Ec}$ |
| *gntZ*_F | AATGCAACUGGGAATCATCGG | Amplification of *gntZ* from the chromosome of *P. putida* KT2440 |
| *gntZ*_R | AGGGGTTCAUTCGCCTTTCTTCTCGAC | Amplification of *gntZ* from the chromosome of *P. putida* KT2440 |
| *rpe*_F | ATGAACCCCUACGCTATTGCCCC | Amplification of *rpe* from the chromosome of *P. putida* KT2440 |
| *rpe*_R | ACGTTCGUAATCATGGGCGGGCCAGGGC | Amplification of *rpe* from the chromosome of *P. putida* KT2440 |
| $P_{14g}$*(BCD10)*_F | ACAAGGCUCTCGCGGCCA | Amplification of $P_{14g}$*(BCD10)* from pS628*(BCD10)→msfGFP* (Wirth and Nikel, 2021) |
| $P_{14g}$*(BCD10)*_R | AGTTGCATUAGAAACGATCCTCCGCATG | Amplification of $P_{14g}$*(BCD10)* from pS628*(BCD10)→msfGFP* |
| pTn7·xylS_F | ACGAACGUAGTGTCCTAGGCCGCGGCCGC | Linearization of pTn7·xylS (Volke et al., 2020) for the integration of $P_{14g}$*(BCD10)*, *gntZ*, and *rpe* |
| pTn7·xylS_R | AGCCTTGUAGGACACTGCACTTTATGCTGGTTATGC | Linearization of pTn7· xylS (Volke et al., 2020) for the integration of $P_{14g}$*(BCD10)*, *gntZ*, and *rpe* |
| seq_attTn7_F | CGAGCTGGTCGTGTTCGC | Genotyping of the *attTn7* region |
| seq_attTn7_R | GCACCAACTTGTTAAAGCCG | Genotyping of the *attTn7* region |

**Table S2. Reactions predicted to contribute to the formation of Ac-CoA on glucose when serially knocked out in iJN1463**. Genes in bold were deleted in strain SCA3. Genes in parentheses were not removed, as being part of a multi-enzyme complex in which at least one protein was deleted. The indicated fluxes represent the predicted rate of the first reaction leading into the respective pathway in wild-type iJN1463.

| Gene | iJN1463 reaction ID | Associated pathway | Flux towards pathway precursor (mmol $g_{CDW}^{-1}$ $h^{-1}$) |
|---|---|---|---|
| ***ace, aceF,*** (*lpdA*) | PDH<br>PDHbr | Pyruvate dehydrogenase complex | 5.76 |
| ***bkdAA***, (*bkdAB, bkdB, lpdV*) | OIVD2<br>OIVD3<br>OIVD1r | Branched chain keto acid dehydrogenase | 0.63<br>0.15<br>0.39 |
| ***ltaE*** | THRAr, THRA2r | L-threonine aldolase | 0.31 |
| *pcaB , pcaC, pcaD* | MUCCY_kt, 4CMLCL_kt, OXOAEL | 3-Dehydroshikimate degradation | 0.25 |
| *paaK* | PACCOAL | L-Phe degradation | 0.12 |
| *davB* | LYSMO | L-Lys degradation | 0.11 |
| *gabT* | APTNAT, ABTA | L-Lys degradation | 0.11 |
| *hmgA, hmgB, hmgC* | HGNTOR, FUMAC, MACACI | L-Tyr degradation | 0.084 |
| *eutB, eutC* | ETHAAL | Ethanolamine degradation | 0.05 *1 |
| *mmsA-I* | MMSAD3 | β-Alanine metabolism | 0.0004 |
| *glcB* | MALS | Malate synthase*2 | - |
| *acoA, acoB, acoC* | | Acetoin dehydrogenase complex*3 | 0 |

*1 The calculated flux towards ethanolamine was only 0.047 mmol $g_{CDW}^{-1}$ $h^{-1}$. However, phosphatidylethanolamine was shown to be the most abundant membrane component in *P. putida* KT2440, constituting about two-thirds of the total glycerophospholipid fraction (**Rühl et al., 2012**). The relatively high abundance of ethanolamine made its degradation appear a plausible potential source of Ac-CoA.

*2 According to calculated flux distributions, the glyoxylate shunt should be inactive to allow for optimal growth with the alternative PK-glycolysis (preventing the use of C2 units as energy source).

*3 The acetoin cleavage system (acetoin dehydrogenase complex) was shown to be functional in *P. putida*, producing acetyl-CoA with a mechanism analogous to that of the PDHc (**Huang et al., 1994**).

**Table S3. Quantitative physiology parameters of engineered strains in minimal medium with glucose and acetate.** Maximum exponential growth rates ($\mu_{max}$) were determined via Gaussian Process regression for growth experiments in microtiter plates. Biomass yields $Y_{X/S}$ in gram cell dry weight ($g_{CDW}$) per gram of substrate ($g_S$) were calculated for each indicated substrate (S). Given are the average values ± standard deviation from at least three biological replicates.

| Strain | Carbon sources | $\mu_{max}$ [$h^{-1}$] | Biomass yield $Y_{X/S}$ [$g_{CDW}$ $g_S^{-1}$] | |
|---|---|---|---|---|
| KT2440 | 30 mM glucose | 0.78 ± 0.07 | 0.45 ± 0.02 | (S: glucose) |
| KT2440 | 30 mM glucose<br>40 mM acetate | 0.98 ± 0.05 | - | |
| KT2440 | 5-20 mM acetate | 0.77 ± 0.14 | 0.42 ± 0.01 | (S: acetate) |
| SCA3 | 30 mM glucose | - | - | |
| SCA3 | 30 mM glucose<br>5 mM acetate | 0.66 ± 0.00 | 0.78 ± 0.02 | (S: acetate) |
| SCA3 | 30 mM glucose<br>10-30 mM acetate | 0.95 ± 0.02 | 0.82 ± 0.03 | (S: acetate) |
| SCA3 | 5-30 mM acetate | 0.61 ± 0.07 | 0.41 ± 0.03 | (S: acetate) |
| SCA15 | 30 mM glucose | No growth | No growth | (S: acetate) |
| SCA15 | 30 mM glucose<br>10-30 mM acetate | 0.53 ± 0.03 | 0.54 ± 0.02 | (S: acetate) |
| SCA15 | 5-30 mM acetate | No growth | No growth | |

**Table S4. Changes in protein expression for strain SCA3 compared to *P. putida* KT2440.** Both strains were cultured in minimal medium supplemented with 30 mM glucose and 30 mM K-Ac. Genes with symbol names in bold were deleted in SCA3. The listed table entries represent significant changes with an adjusted p-value threshold of 0.05, filtered for proteins with an identified enzymatic function according to the KEGG database (https://www.genome.jp/kegg-bin/show_pathway?ppu01100). Abundance changes with a $\log_2$(fold change) of ≥ 2 are highlighted with bold numbers. *n.d.*, not determined

| Ensembl Gene ID | Enzyme | Gene Symbol | Fold-change |
|---|---|---|---|
| *PP_5270* | **D-amino acid:quinone oxidoreductase** | *dadA-2; dadA-II* | **8.90** |
| *PP_1389* | **oxaloacetate decarboxylase** | | **6.44** |
| *PP_0542* | **ethanolamine ammonia-lyase subunit β** | *eutC* | **6.04** |
| *PP_2679* | **quinoprotein ethanol dehydrogenase** | *qedH-II* | **4.50** |
| *PP_4636* | **acetyl-CoA acetyltransferase** | *yqeF* | **3.50** |
| *PP_4116* | **isocitrate lyase** | *aceA* | **3.49** |
| *PP_3148* | **glutamine synthetase** | | **3.47** |
| *PP_5132* | **dihydrofolate reductase** | *folA* | **2.84** |
| *PP_0596* | **ω-amino acid–pyruvate aminotransferase** | | **2.73** |
| *PP_3376* | **phosphonate dehydrogenase** | *kguD; ptxD* | **2.72** |
| *PP_2341* | **6-carboxytetrahydropterin synthase QueD** | *queD* | **2.25** |
| *PP_1347* | **glutathione *S*-transferase family protein** | | **2.12** |
| *PP_0372* | **acetylornithine aminotransferase** | *aruC* | **2.05** |
| *PP_0356* | **malate synthase G** | *glcB* | **2.03** |
| *PP_0751* | **malate:quinone oxidoreductase** | *mqo-1; mqo-I* | 1.94 |
| *PP_5332* | **hypothetical protein** | | 1.93 |
| *PP_2151* | **pyridine nucleotide transhydrogenase** | *sthA* | 1.91 |
| *PP_2913* | **5-aminolevulinate dehydratase** | *hemB-1; hemB* | 1.81 |
| *PP_1261* | **2-ketoaldonate reductase/hydroxypyruvate/glyoxylate reductase** | *ghrB* | 1.71 |
| *PP_1832* | **oxidase** | | 1.67 |
| *PP_0762* | **glycerate dehydrogenase** | *hprA* | 1.60 |
| *PP_2137* | **β-ketoadipyl-CoA thiolase subunit β** | *fadA; pcaF-II* | 1.55 |
| *PP_4577* | **hypothetical protein** | | 1.53 |
| *PP_0840* | **serine acetyltransferase** | *cysE* | 1.51 |
| *PP_4012* | **isocitrate dehydrogenase** | *idh* | 1.50 |
| *PP_1332* | **UDP-N-acetylmuramoyl-L-alanyl-D-glutamate–2,6-diaminopimelate ligase** | *murE* | 1.45 |
| *PP_5183* | **glutamylpolyamine synthetase** | *spuB* | 1.45 |
| *PP_0392* | **dihydroneopterin aldolase** | *folB* | 1.44 |
| *PP_1989* | **aspartate-semialdehyde dehydrogenase** | *asd* | 1.44 |
| *PP_1346* | **glutamate *N*-acetyltransferase/amino acid acetyltransferase** | *argJ* | 1.39 |
| *PP_1031* | **IMP dehydrogenase and single strand DNA binding factor** | *guaB* | 1.36 |
| *PP_1823* | **GTP cyclohydrolase I** | *folE; folEA-I* | 1.34 |
| *PP_0722* | **ribose-phosphate pyrophosphokinase** | *prsA; prs* | 1.32 |
| *PP_2536* | **glutathione *S*-transferase family protein** | | 1.29 |
| *PP_1657* | **modified nucleoside triphosphate pyrophosphohydrolase** | *mazG* | 1.25 |
| *PP_0517* | **6,7-dimethyl-8-ribityllumazine synthase** | *ribH; PSEEN_RS02765* | 1.17 |
| *PP_1338* | **UDP-N-acetylmuramate-L-alanine ligase** | *murC* | 1.15 |
| *PP_1530* | **2,3,4,5-tetrahydropyridine-2,6-dicarboxylate N-succinyltransferase** | *dapD* | 1.10 |
| *PP_0967* | **histidinol-phosphate aminotransferase** | *hisC* | 0.81 |
| *PP_0516* | **bifunctional 3,4-dihydroxy-2-butanone 4-phosphate synthase/GTP cyclohydrolase II** | *ribBA-1; ribAB-I* | 0.80 |

| | | | |
|---|---|---|---|
| *PP_1403* | **β-D-glucoside glucohydrolase** | *bglX* | 0.78 |
| *PP_1616* | **glutathione-dependent formaldehyde dehydrogenase** | *frmA* | 0.78 |
| *PP_1611* | **2-dehydro-3-deoxyphosphooctonate aldolase** | *kdsA-1; kdsA-I* | 0.77 |
| *PP_0292* | **phosphoribosylformimino-5-aminoimidazole carboxamide ribonucleotide isomerase** | *hisA* | 0.76 |
| *PP_0560* | **type II 3-dehydroquinate dehydratase** | *aroQ-1; aroQ-I* | 0.76 |
| *PP_0842* | **cysteine desulfurase** | *iscS; iscS-I* | 0.75 |
| *PP_1470* | **homoserine dehydrogenase** | *hom* | 0.75 |
| *PP_5285* | **bifunctional 4'-phosphopantothenoylcysteine decarboxylase/phosphopantothenoylcysteine synthetase** | *coaBC; dfp* | 0.75 |
| *PP_1807* | **2-dehydro-3-deoxyphosphooctonate aldolase** | *kdsA-2; kdsA-II* | 0.73 |
| *PP_3219* | **alkansulfonate monooxygenase** | | 0.72 |
| *PP_0366* | **dethiobiotin synthetase** | *bioD* | 0.72 |
| *PP_1593* | **uridylate kinase** | *pyrH* | 0.71 |
| *PP_4122* | **NADH-quinone oxidoreductasesubunit E** | *nuoE* | 0.71 |
| *PP_1617* | ***S*-formylglutathione hydrolase/S-lactoylglutathione hydrolase** | *frmC* | 0.71 |
| *PP_1231* | **quinolinate synthase iron-sulfur cluster subunit** | *nadA* | 0.70 |
| *PP_0328* | **formaldehyde dehydrogenase** | *fdhA* | 0.69 |
| *PP_0421* | **anthranilate phosphoribosyltransferase** | *trpD* | 0.68 |
| *PP_2371* | **sulfite reductase hemoprotein subunit β** | *cysI* | 0.67 |
| *PP_4123* | **NADH-quinone oxidoreductase subunit F** | *nuoF* | 0.67 |
| *PP_5149* | **threonine deaminase** | *ilvA-2; ilvA-II* | 0.66 |
| *PP_1677* | **adenosyl-cobyric acid synthase** | *cobQ* | 0.65 |
| *PP_0581* | **3-oxoacyl-ACP reductase** | *fabG;* | 0.65 |
| *PP_3540* | **hydroxymethylglutaryl-CoA lyase** | *mvaB* | 0.64 |
| *PP_4126* | **NADH-quinone oxidoreductase subunit I** | *nuoI* | 0.63 |
| *PP_4186* | **bifunctional succinyl-CoA synthetase subunit β/glutaryl-CoA synthetase subunit β** | *sucC* | 0.63 |
| *PP_1610* | **CTP synthase** | *pyrG* | 0.63 |
| *PP_4174* | **3*R*-3-hydroxydecanoyl-ACP dehydratase** | *fabA* | 0.61 |
| *PP_4185* | **succinyl-CoA synthetase subunit α** | *sucD* | 0.61 |
| *PP_0083* | **tryptophan synthase subunit β** | *trpB* | 0.61 |
| *PP_2339* | **aconitate hydratase B** | *acnB* | 0.60 |
| *PP_4042* | **Glucose 6-phosphate 1-dehydrogenase** | *zwfA* | 0.59 |
| *PP_4679* | **acetolactate synthase small subunit** | *ilvH* | 0.57 |
| *PP_5075* | **glutamate synthase subunit β** | *gltD* | 0.57 |
| *PP_2017* | **aminopeptidase N** | *pepN* | 0.56 |
| *PP_0582* | **thiolase family protein** | | 0.55 |
| *PP_0794* | **1-phosphofructokinase monomer** | *fruK* | 0.55 |
| *PP_1481* | **medium chain aldehyde dehydrogenase** | *patD* | 0.54 |
| *PP_0989* | **glycine cleavage system protein H** | *gcvH; gcvH-I* | 0.54 |
| *PP_4065* | **methylcrotonyl-CoA carboxylase biotin-containing subunit β** | *mccB* | 0.52 |
| *PP_0988* | **glycine dehydrogenase** | *gcvP-1; gcvP-I* | **0.50** |
| *PP_0417* | **anthranilate synthase component 1** | *trpE* | **0.49** |
| *PP_4479* | **arginine N-succinyltransferase subunit β** | *aruG; astA-I* | **0.49** |
| *PP_3410* | **precorrin-4 C(11)-methyltransferase** | *cobM* | **0.48** |
| *PP_0323* | **sarcosine oxidase subunit β** | *soxB* | **0.48** |
| *PP_4922* | **phosphomethylpyrimidine synthase** | *thiC* | **0.47** |
| *PP_5186* | **acetylornithine deacetylase** | *argE* | **0.46** |
| *PP_4124* | **NADH-quinone oxidoreductase subunit G** | *nuoG* | **0.46** |
| *PP_0073* | **coproporphyrinogen-III oxidase** | *hemF* | **0.44** |
| *PP_2338* | **2-methylcitrate dehydratase** | *prpD* | **0.43** |
| *PP_3189* | **cytosine deaminase/isoguanine deaminase** | *codA* | **0.42** |
| *PP_5346* | **pyruvate carboxylase subunit B** | *oadA; pycB* | **0.42** |
| *PP_1023* | **6-phosphogluconolactonase** | *pgl* | **0.41** |

| | | | |
|---|---|---|---|
| *PP_2149* | **glyceraldehyde-3-phosphate dehydrogenase** | *gap-2; gapB* | **0.41** |
| *PP_0986* | **aminomethyltransferase** | *gcvT-1; gcvT-I* | **0.41** |
| *PP_1874* | **hydroperoxy fatty acid reductase Gpx1** | *gpx* | **0.39** |
| *PP_5128* | **dihydroxy-acid dehydratase** | *ilvD* | **0.38** |
| *PP_0365* | **malonyl-ACP *O*-methyltransferase** | *bioC* | **0.38** |
| *PP_0363* | **8-amino-7-oxononanoate synthase** | *bioF* | **0.37** |
| *PP_0293* | **imidazole glycerol phosphate synthase subunit HisF** | *hisF* | **0.37** |
| *PP_3416* | **D-gluconate kinase** | *gnuK* | **0.36** |
| *PP_0339* | **pyruvate dehydrogenase E1 component** | ***aceE*** | **0.30** |
| *PP_5033* | **urocanate hydratase** | *hutU* | **0.30** |
| *PP_0362* | **biotin synthase** | *bioB* | **0.29** |
| *PP_5347* | **pyruvate carboxylase subunit A** | *accC-2; pycA* | **0.29** |
| *PP_5184* | **glutamylpolyamine synthetase** | *spuI* | **0.25** |
| *PP_4977* | **5,10-methylenetetrahydrofolate reductase** | *metF* | **0.21** |
| *PP_3790* | **diaminopimelate epimerase** | *dapF;* | **0.20** |
| *PP_4967* | **methionine adenosyltransferase** | *metK* | **0.17** |
| *PP_0321* | **Low specificity L-threonine aldolase** | ***ltaE*** | **0.03** |
| *PP_0338* | **AceF-S-acetyldihydrolipoate** | ***aceF*** | **0.01** |
| *PP_4401* | **2-oxoisovalerate dehydrogenase subunit $\alpha$** | ***bkdAA*** | n.d. |

**Table S5. Genes located within the chromosomal region deleted in two out of four evolved SCA3$_{PK\text{-}Tn}$/pS438·*pta*$^{Ec}$.**

| Ensembl Gene ID | Enzyme | Gene Symbol |
|---|---|---|
| *PP_2985* | hypothetical protein | |
| *PP_5548* | hypothetical protein | |
| *PP_2986* | oxidoreductase | |
| *PP_2987* | hypothetical protein | |
| *PP_2988* | alcohol dehydrogenase | |
| *PP_2989* | short-chain dehydrogenase/reductase family oxidoreductase | |
| *PP_2990* | MerR family transcriptional regulator | |
| *PP_2991* | hypothetical protein | |
| *PP_2992* | hypothetical protein | |
| *PP_2994* | FMN-binding oxidoreductase | |
| *PP_2995* | DNA topology modulation kinase FlaR | |
| *PP_2996* | hypothetical protein | |
| *PP_2997* | protein CnmA | |
| *PP_2998* | 2-dehydropantoate 2-reductase | |
| *PP_2999* | glyoxalase family protein | |
| *PP_3000* | MaoC domain-containing protein | |
| *PP_3001* | CAIB/BAIF family protein | |
| *PP_3002* | shikimate dehydrogenase-like protein | |
| *PP_3003* | 3-dehydroquinate dehydratase | *aroQ-III* |
| *PP_3004* | hypothetical protein | |
| *PP_3005* | hypothetical protein | |
| *PP_3006* | EcF family RNA polymerase sigma-70 factor | |
| *PP_3007* | hypothetical protein | |
| *PP_3008* | hypothetical protein | |
| *PP_3009* | hypothetical protein | |
| *PP_3010* | hypothetical protein | |
| *PP_3011* | hypothetical protein | |
| *PP_3012* | hypothetical protein | |
| *PP_3013* | hypothetical protein | |
| *PP_3014* | hypothetical protein | |
| *PP_5549* | hypothetical protein | |
| *PP_3015* | medium-chain acyl-CoA ligase domain-containing protein | |
| *PP_3016* | lipopolysaccharide core biosynthesis protein | |
| *PP_3017* | O6-methylguanine-DNA methyltransferase | *ogt* |
| *PP_3018* | hypothetical protein | |
| *PP_3019* | carbon-nitrogen hydrolase family protein | |
| *PP_3020* | serine/threonine protein phosphatase | |
| *PP_3021* | LysE family transporter | |
| *PP_3022* | AraC family transcriptional regulator | |
| *PP_3024* | hypothetical protein | |
| *PP_5550* | hypothetical protein | |
| *PP_5551* | hypothetical protein | |
| *PP_3025* | LysE family transporter | |
| *PP_3026* | recombinase | |
| *PP_3027* | hypothetical protein | |
| *PP_3028* | hypothetical protein | |
| *PP_3029* | DNA cytosine methyltransferase family protein | |
| *PP_3030* | hypothetical protein | |
| *PP_3031* | Pyocin activator protein PrtN | *prtN* |
| *PP_3032* | DNA-binding protein Roi-like protein | |

| | | |
|---|---|---|
| *PP_5552* | **hypothetical protein** | |
| *PP_5553* | **hypothetical protein** | |
| *PP_3033* | **CI/C2 family transcriptional regulator** | |
| *PP_3034* | **hypothetical protein** | |
| *PP_3035* | **hypothetical protein** | |
| *PP_5554* | **hypothetical protein** | |
| *PP_3036* | **hypothetical protein** | |
| *PP_5555* | **hypothetical protein** | |
| *PP_3037* | **DksA/TraR family C4-type zinc finger protein** | |
| *PP_3038* | **TraC domain-containing protein** | |
| *PP_3039* | **hypothetical protein** | |
| *PP_3040* | **holin** | |
| *PP_3041* | **hypothetical protein** | |
| *PP_3042* | **terminase large subunit** | |
| *PP_3043* | **hypothetical protein** | |
| *PP_3044* | **lambda family portal protein** | |
| *PP_3045* | **ATP-dependent protease ClpP** | |
| *PP_3046* | **hypothetical protein** | |
| *PP_3047* | **hypothetical protein** | |
| *PP_3048* | **hypothetical protein** | |
| *PP_3049* | **hypothetical protein** | |
| *PP_3050* | **hypothetical protein** | |
| *PP_3051* | **pyocin R2_PP** | *gpV* |
| *PP_3052* | **hypothetical protein** | |
| *PP_3053* | **pyocin R2_PP baseplate protein** | *gpW* |
| *PP_3054* | **pyocin R2_PP baseplate/tail fiber protein** | *gpJ* |
| *PP_3055* | **pyocin R2_PP tail formation protein** | *gpI* |
| *PP_3056* | **tail fiber protein** | |
| *PP_3057* | **pectinesterase** | |
| *PP_3058* | **hypothetical protein** | |
| *PP_3059* | **tail sheath protein** | |
| *PP_3060* | **tail sheath protein** | |
| *PP_3061* | **hypothetical protein** | |
| *PP_3062* | **tail length determination protein** | |
| *PP_3063* | **pyocin R2_PP tail formation protein** | *gpU* |
| *PP_3064* | **pyocin R2_PP tail component** | *gpX* |
| *PP_3065* | **pyocin R2_PP tail formation protein** | *gpD* |
| *PP_3066* | **lytic enzyme** | |
| *PP_5556* | **bacteriophage lysis protein** | |
| *PP_5557* | **hypothetical protein** | |
| *PP_3067* | **hypothetical protein** | |
| *PP_5558* | **Xre family transcriptional regulator** | |
| *PP_3069* | **outer membrane autotransporter** | |
| *PP_3070* | **PpiC-type peptidyl-prolyl *cis-trans* isomerase** | *ppiC-I* |
| *PP_3071* | **acetoacetyl-CoA synthetase** | *aacs* |
| *PP_3072* | **ecotin** | *eco* |
| *PP_3073* | **3-hydroxybutyrate dehydrogenase** | *hbdH* |
| *PP_3074* | **citrate transporter** | *bhbP* |
| *PP_3075* | **transcriptional regulator** | |
| *PP_3076* | **ABC transporter permease** | |
| *PP_3077* | **ABC transporter permease** | |
| *PP_3078* | **ABC transporter substrate-binding protein** | |
| *PP_3079* | **peptidyl-prolyl *cis-trans* isomerase** | *ppiC-II* |
| *PP_3080* | **phospho-2-dehydro-3-deoxyheptonate aldolase** | *aroF-II* |
| *PP_5559* | **hypothetical protein** | |
| *PP_3081* | **hypothetical protein** | |

| | |
|---|---|
| *PP_5560* | **hypothetical protein** |
| *PP_3082* | **membrane protein** |
| *PP_3083* | **membrane protein** |
| *PP_3084* | **ferric siderophore receptor** |
| *PP_3085* | **transmembrane sensor** |
| *PP_3086* | **RNA polymerase sigma-70 factor** |
| *PP_3087* | **excinuclease ABC subunit A** |
| *PP_3088* | **hypothetical protein** |
| *PP_3089* | **hypothetical protein** |

**Table S6. Selected reactions involved in the formation of electron carriers and their flux ratios determined via $^{13}$C-based metabolic flux analysis.** Numbers represent relative fluxes (in %), normalized to the measured glucose consumption for each strain.

| Reaction | Equation | Flux ratio in SCA3$_{PK\text{-}Tn}$/pS438·*pta*$^{Ec}$ | Flux ratio in wildtype |
|---|---|---|---|
| **NADH-producing reactions** | | | |
| G3P dehydrogenase | G3P <-> 3PG + ATP + NADH | 40.5 | 58.3 |
| Pyruvate dehydrogenase | Pyr -> AcCoA + CO2 + NADH | 0 | 81.3 |
| 2-OG dehydrogenase | AKG -> SucCoA + CO2 + NADH | 0 | 39.4 |
| Malate dehydrogenase | Mal -> OAA + NADH | 3.2 | 70.0 |
| | | 43.7 (total) | 249 (total) |
| **NADPH-producing reactions** | | | |
| G6P dehydrogenase | G6P -> 6PG + NADPH | 21.9 | 29.7 |
| 6PG dehydrogenase | 6PG -> Ri5P + CO2 + NADPH | 36.7 | 3.9 |
| Isocitrate dehydrogenase | ICit -> AKG + CO2 + NADPH | 9.7 | 44.3 |
| Malic enzyme | Mal -> Pyr + CO2 + NADPH | 10.5 | 0.8 |
| Methylenetetrahydrofolate dehydrogenase | MEETHF -> FTHF + NADPH | 0 | 6.6 |
| | | 78.8 (total) | 85.3 (total) |
| **UQH2-producing reactions** | | | |
| Glc dehydrogenase | Gluc.per -> Gluco.per + UQH2 | 94.3 | 89.2 |
| **FADH2-producing reactions** | | | |
| Glcnt dehydrogenase | Gluco.per -> Kgluco.per + FADH2 | 77.1 | 2.6 |
| Suc-CoA dehydrogenase | Suc -> Fum + FADH2 | 6.8 | 51.8 |
| | | 83.9 (total) | 54.4 (total) |
| **Consumption of reducing equivalents** | NADH + O2 -> 3*ATP | 44.6 | 269.3 |
| | FADH2 + O2 -> 2*ATP | 83.9 | 54.4 |
| | UQH2 + O2 -> 3*ATP | 94.3 | 89.2 |
| | AKG + NADPH + NH3 -> Glu | 16.9 | 84.6 |
| | Glu + ATP + 2*NADPH -> Pro | 6.7 | 0 |

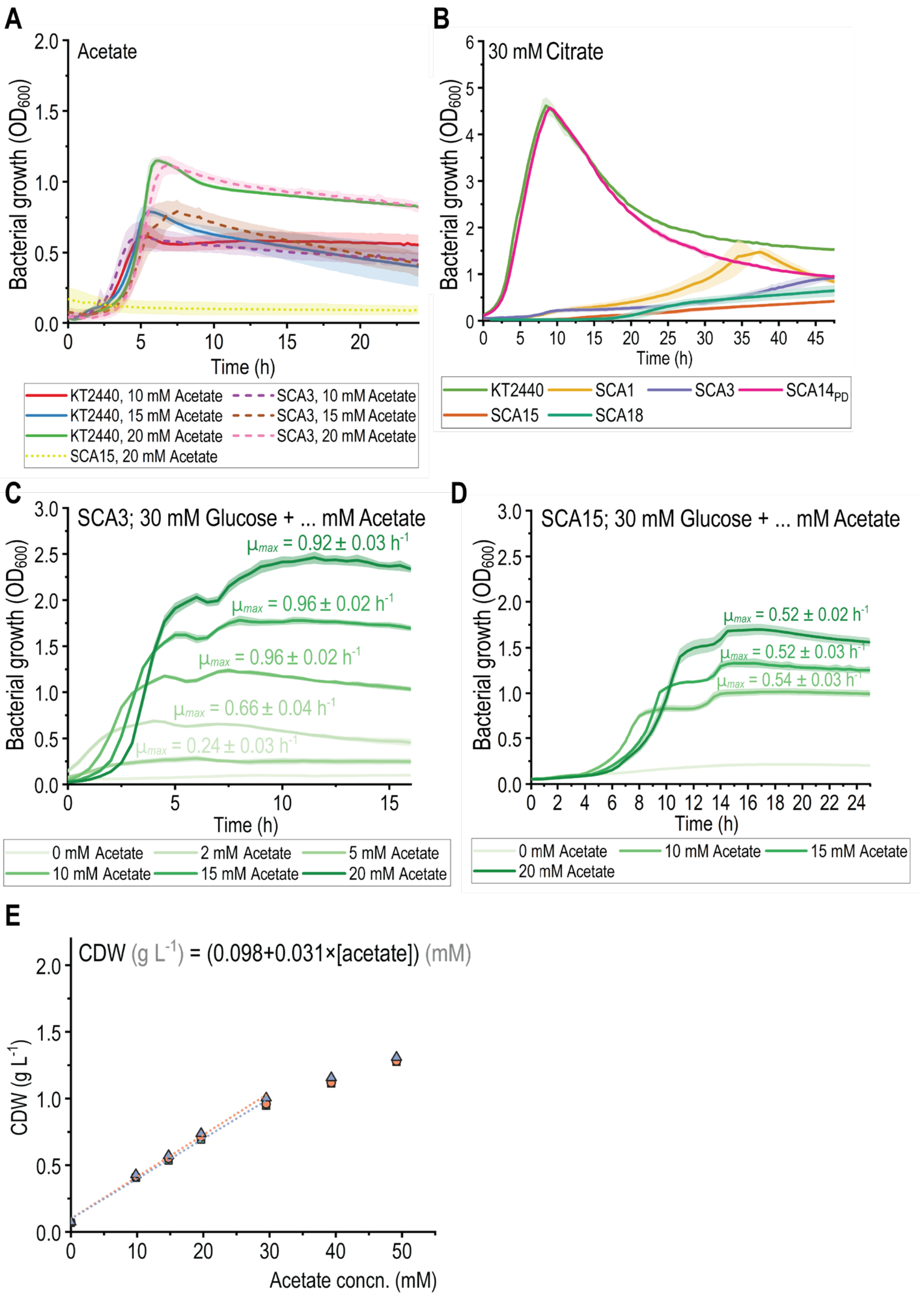

**Figure S1. Phenotypical characterization of synthetic C2-auxotroph strains.** SCA strains and wild-type *P. putida* KT2440 were cultured in microtiter plates (96-well). **(A)** Growth in DBM medium supplemented with 10 to 20 mM acetate as the sole source of carbon. **(B)** Growth in DBM medium supplemented with 30 mM citrate as the sole source of carbon. **(C)** Growth of strain SCA3 in DBM medium supplemented with 30 mM glucose and varying concentrations of acetate. **(D)** Growth of strain SCA15 in DBM medium supplemented with 30 mM glucose and varying concentrations of acetate. **(E)** Dependency of strain SCA15 on acetate for biomass formation (indicated as the concentration of cell dry weight, CDW). The strain was grown in DBM medium supplemented with 30 mM glucose and varying concentrations of acetate. Shown are the individual measurements and linear correlations of three biological replicates. Error bars, representing the standard deviation from three biological replicates, are indicated by filled areas around the curves. *CDW*, cell dry weight; *concn.*, concentration.

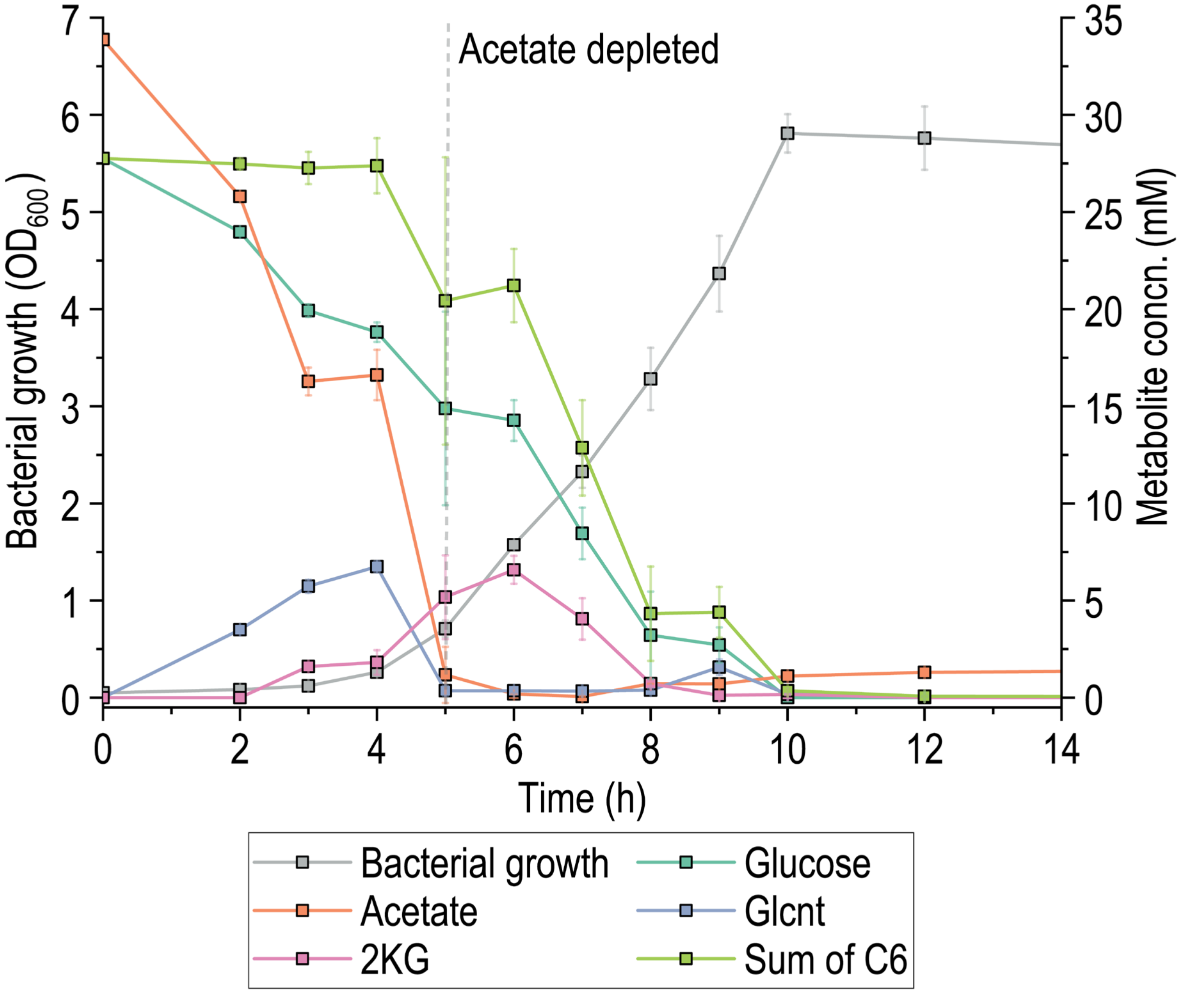

**Figure S2. Co-consumption of glucose and acetate in *P. putida* KT2440.** The strain was grown in DBM medium supplemented with 5 g $L^{-1}$ (28 mM) glucose and 2 g $L^{-1}$ (34 mM) acetate. To distinguish the utilization of glucose as energy source as opposed to the assimilation of its carbon skeleton, the three six-carbon (C6) moieties glucose (Glc), gluconate (Glcnt), and 2-ketogluconate (2KG) were combined into the *Sum of C6.* During acetate consumption, glucose is only used as an energy source (via the oxidation to Glcnt and 2KG). The assimilation of glucose-derived carbon is initiated after acetate has been depleted (indicated by a dashed line). Error bars represent the standard deviation from three biological replicates. *concn.*, concentration.

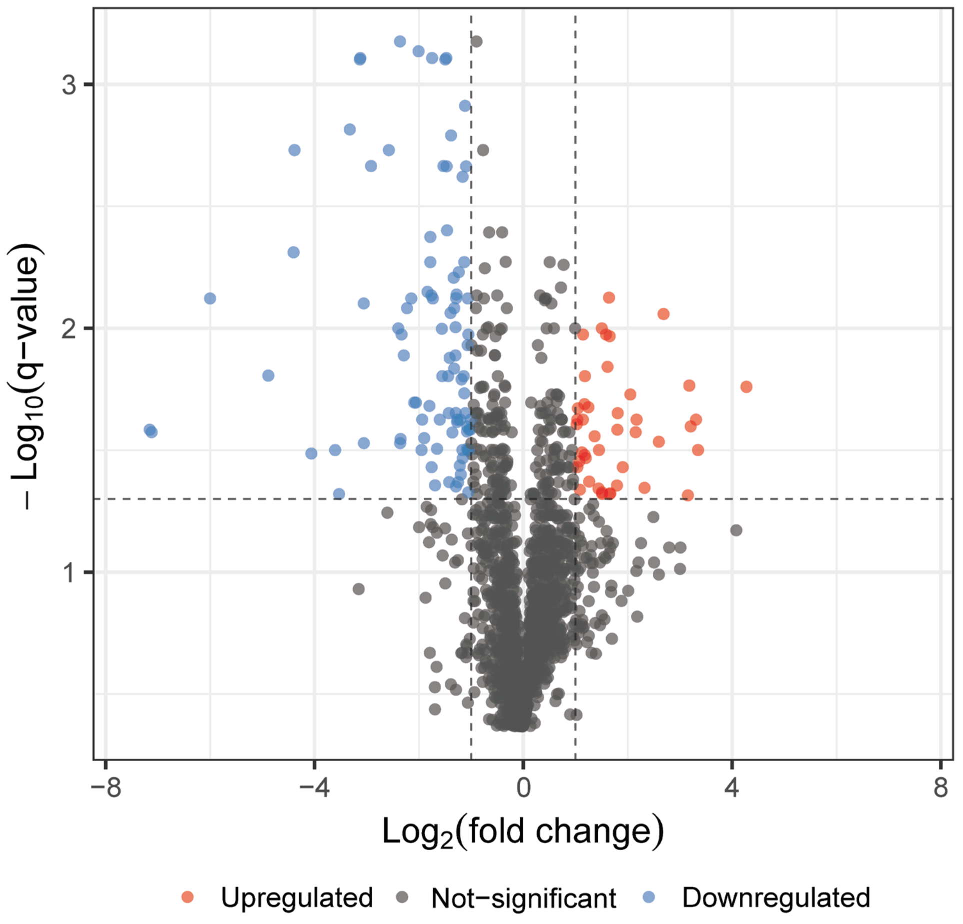

**Figure S3. Volcano plot of protein abundance data showing differentially expressed proteins in strain SCA3 compared to *P. putida* KT2440.** Both strains (with three biological replicates) were grown in DBM medium supplemented with 30 mM glucose and 30 mM K-Ac and harvested in the mid-exponential phase to measure the soluble protein content of biomass. Each point in the plot represents an individual protein. The horizontal intersection is set at a q-value (FDR) of ≤ 0.05; the vertical intersections are placed for an absolute fold-change of ≥ 2.

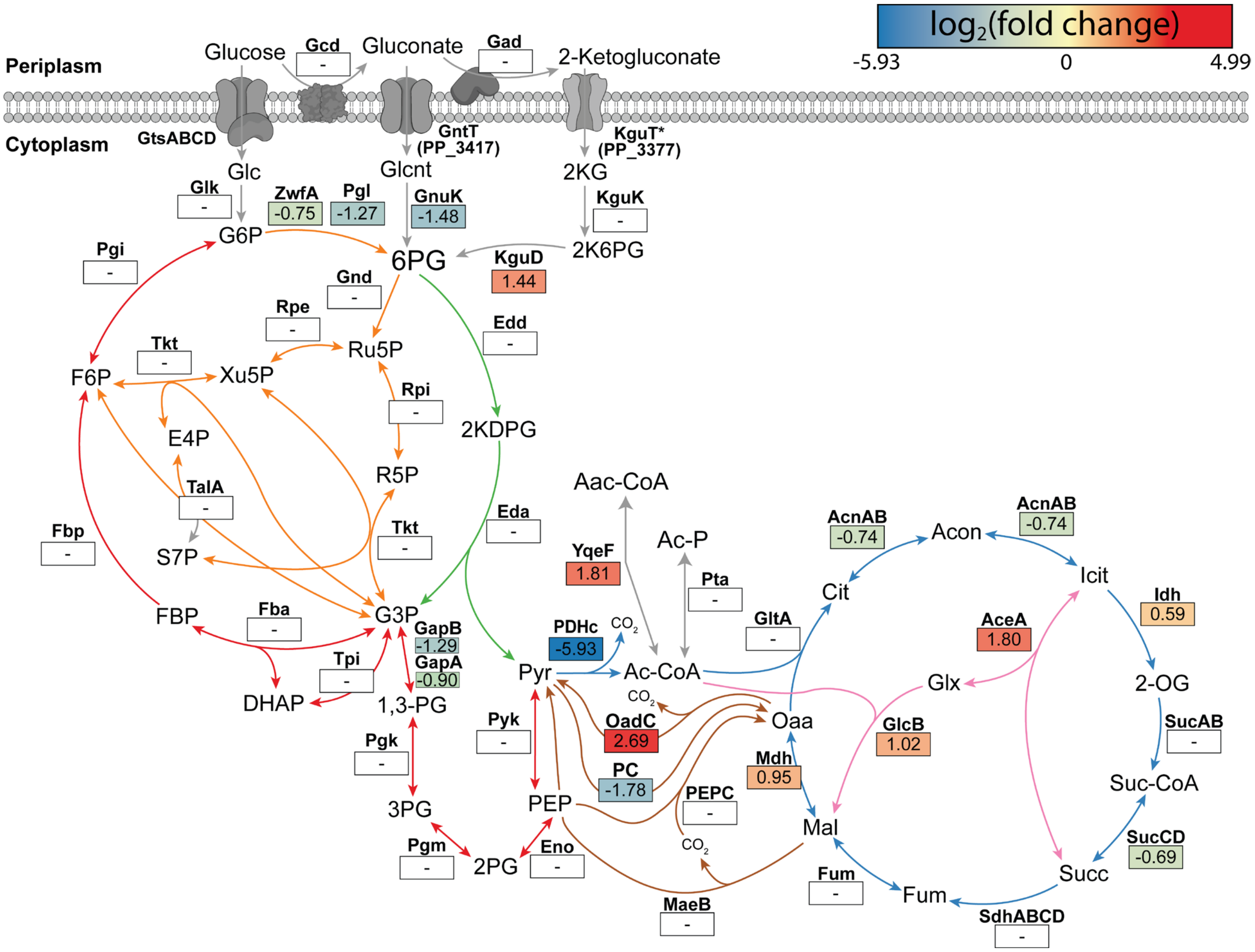

**Figure S4. Changes in protein expression within central carbon metabolism for strain SCA3 compared to *P. putida* KT2440. The two strains were grown in DBM medium supplemented with 30 mM glucose and 30 mM acetate.** Shown are the $log_2$(fold change) values for significantly differentially expressed proteins (p-value ≤ 0.05). The reactions are categorized into the ED pathway (green), the PP (orange), the EMP route (red), the TCA cycle (blue), the glyoxylate shunt (pink), and the anaplerotic reactions (brown). A list of abbreviations is given in **Figure 1** in the main text. Additional abbreviations: Aac-CoA, acetoacetyl-coenzyme A; YqeF, acetyl-CoA acetyltransferase; OadC, oxaloacetate decarboxylase.

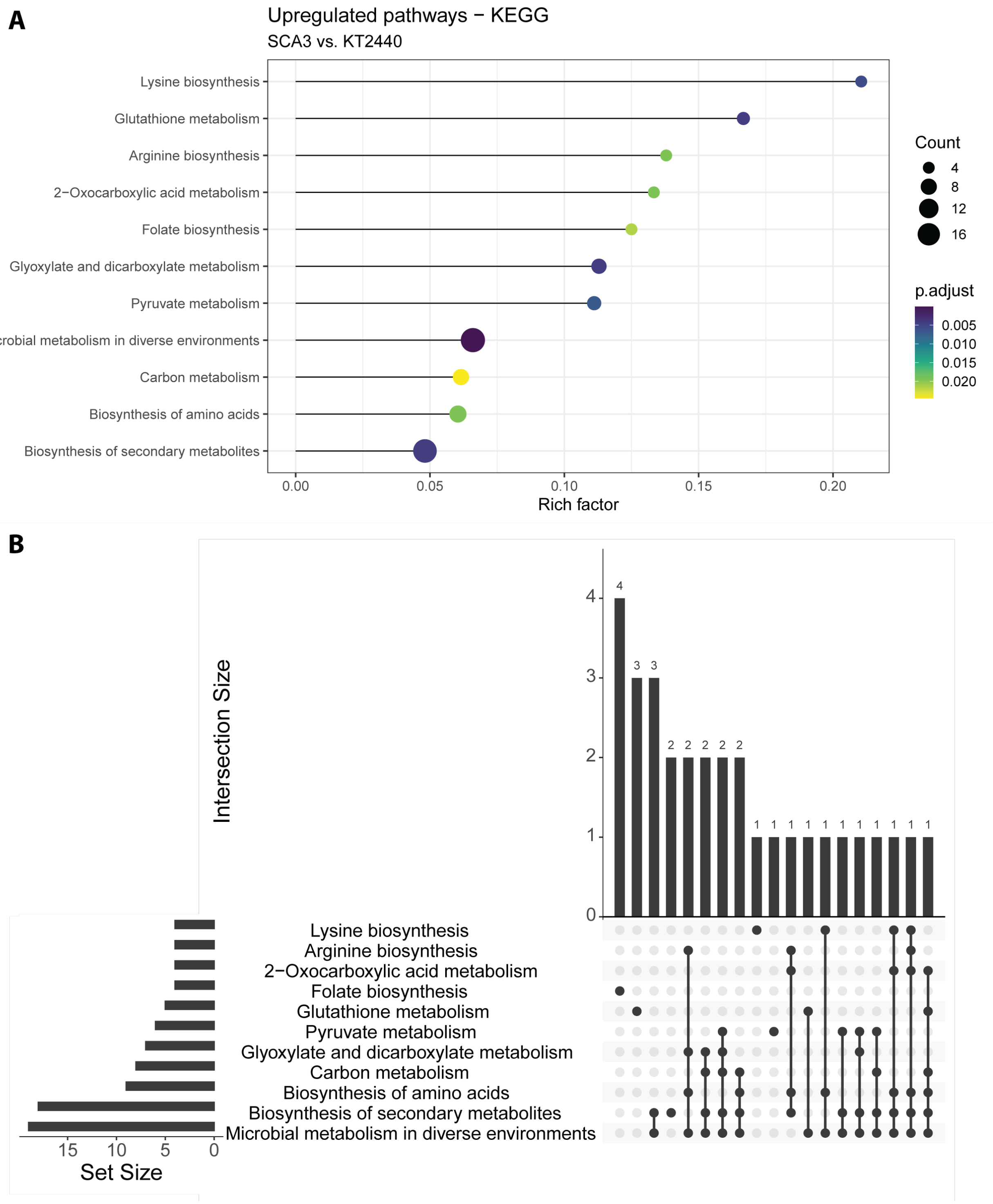

**Figure S5. Differentially enriched pathways (upregulated) in evolved vs. pre-evolved SCA3$_{PK\text{-}Tn}$/pS438·*pta*$^{Ec}$. (A)** Pathways found to be significantly upregulated with a threshold for the adjusted p value (Benjamini-Hochberg method) of 0.1. The rich factor indicates what fraction of the total sum of proteins constituting a pathway was found to be differentially expressed. **(B)** Upset plot of proteins identified in the pathway enrichment analysis.

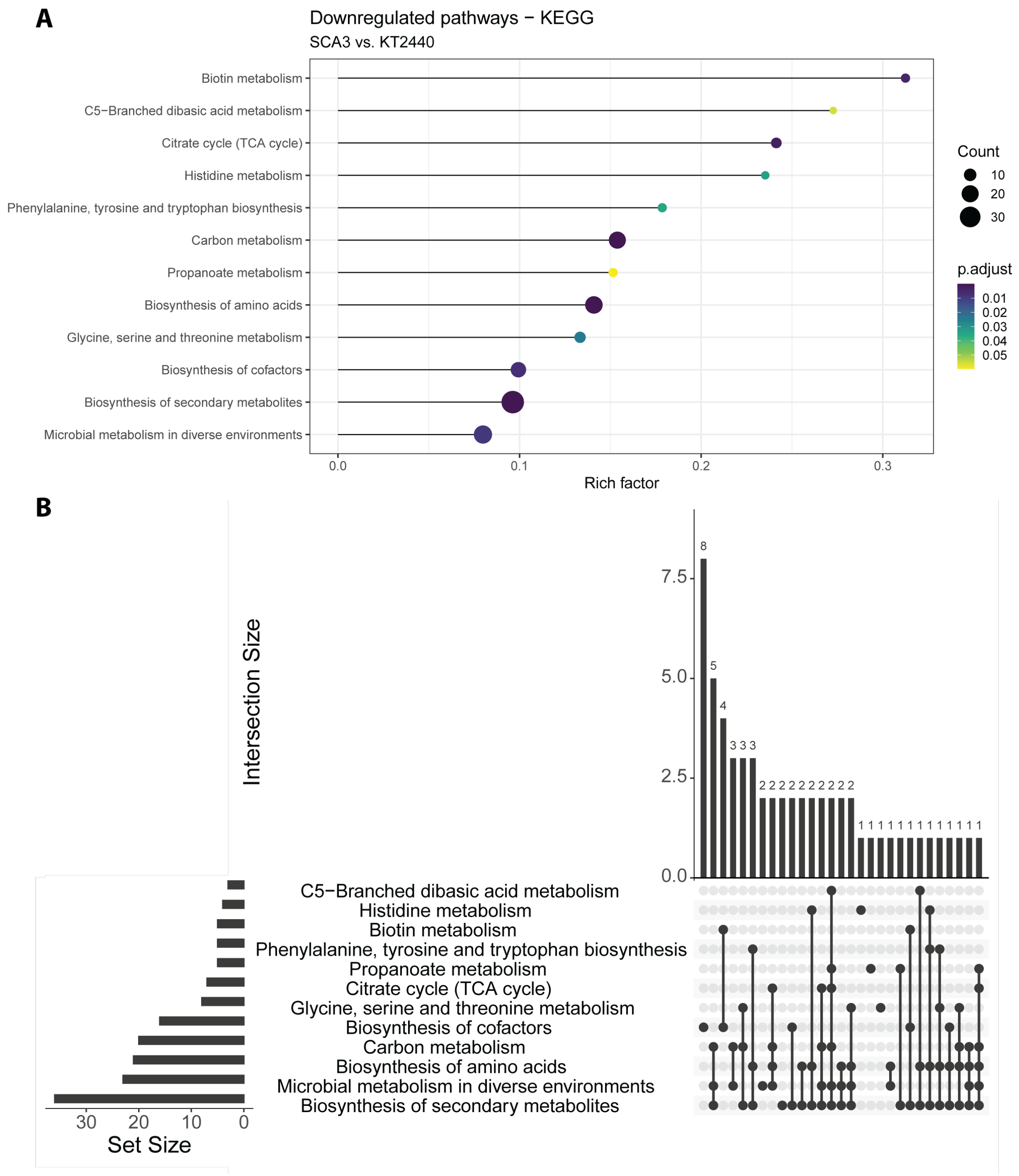

**Figure S6. Differentially enriched pathways (downregulated) in evolved vs. pre-evolved SCA3$_{PK\text{-}Tn}$/pS438·*pta*$^{Ec}$. (A)** Pathways found to be significantly downregulated with a threshold for the adjusted p value (Benjamini-Hochberg method) of 0.1. The rich factor indicates what fraction of the total sum of proteins constituting a pathway was found to be differentially expressed. **(B)** Upset plot of proteins identified in the pathway enrichment analysis.

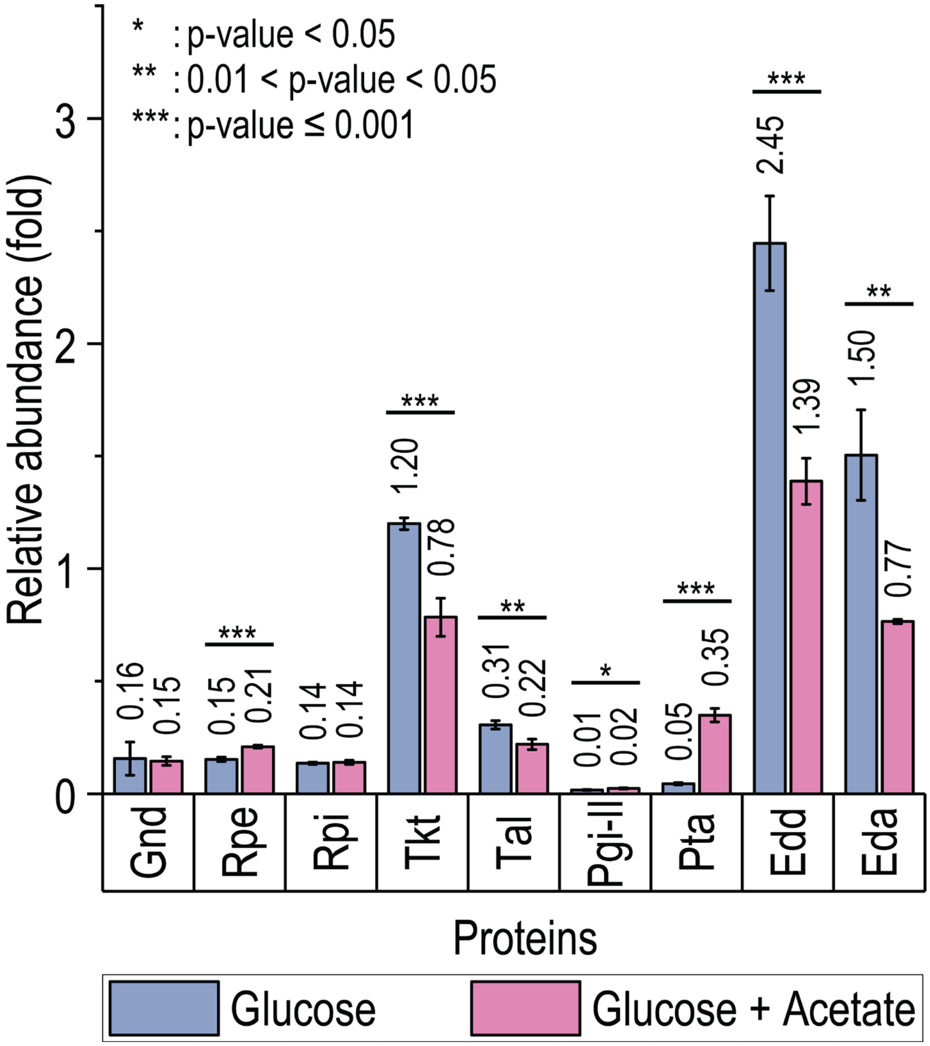

**Figure S7. Protein abundance of selected central carbon metabolism genes in *P. putida* KT2440 grown on glucose and glucose + acetate.** The strain was cultures in minimal medium supplemented with 30 mM glucose ± 30 mM K-Ac. *Vsn*-normalized values were normalized to the abundance of RpoD in each sample. Error bars indicate the standard deviation of three biological replicates.

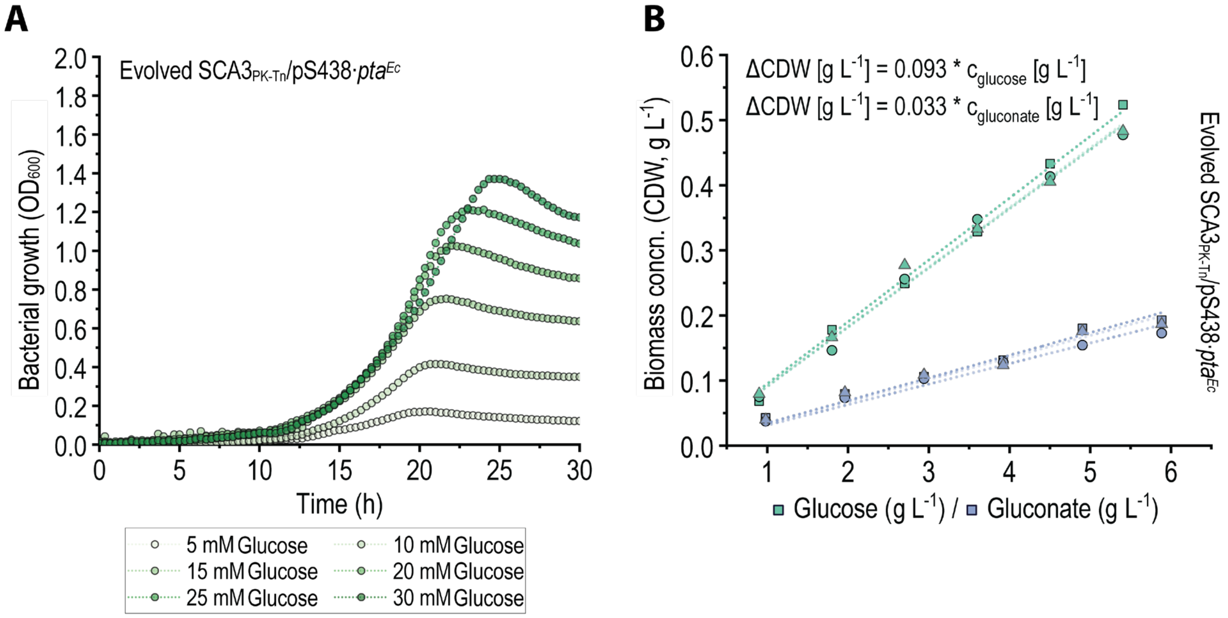

**Figure S8. Effect of evolution on the growth physiology of strain SCA3$_{PK-Tn}$/pS438·*pta*$^{Ec}$. (A)** 96-well cultivation of evolved SCA3$_{PK-Tn}$/pS438·*pta*$^{Ec}$ in DBM medium supplemented with varying glucose concentrations and 0.5 mM 3-*m*Bz. **(B)** Linear correlation of biomass produced by evolved SCA3$_{PK-Tn}$/pS438·*pta*$^{Ec}$on glucose and gluconate. The slope and intersection values in the equations displayed represent the average values from three independent correlations performed on biological replicates. Maximum observed growth rates and biomass yields are summarized in **Table S3**. *concn.*, concentration; *CDW*, cell dry weight.

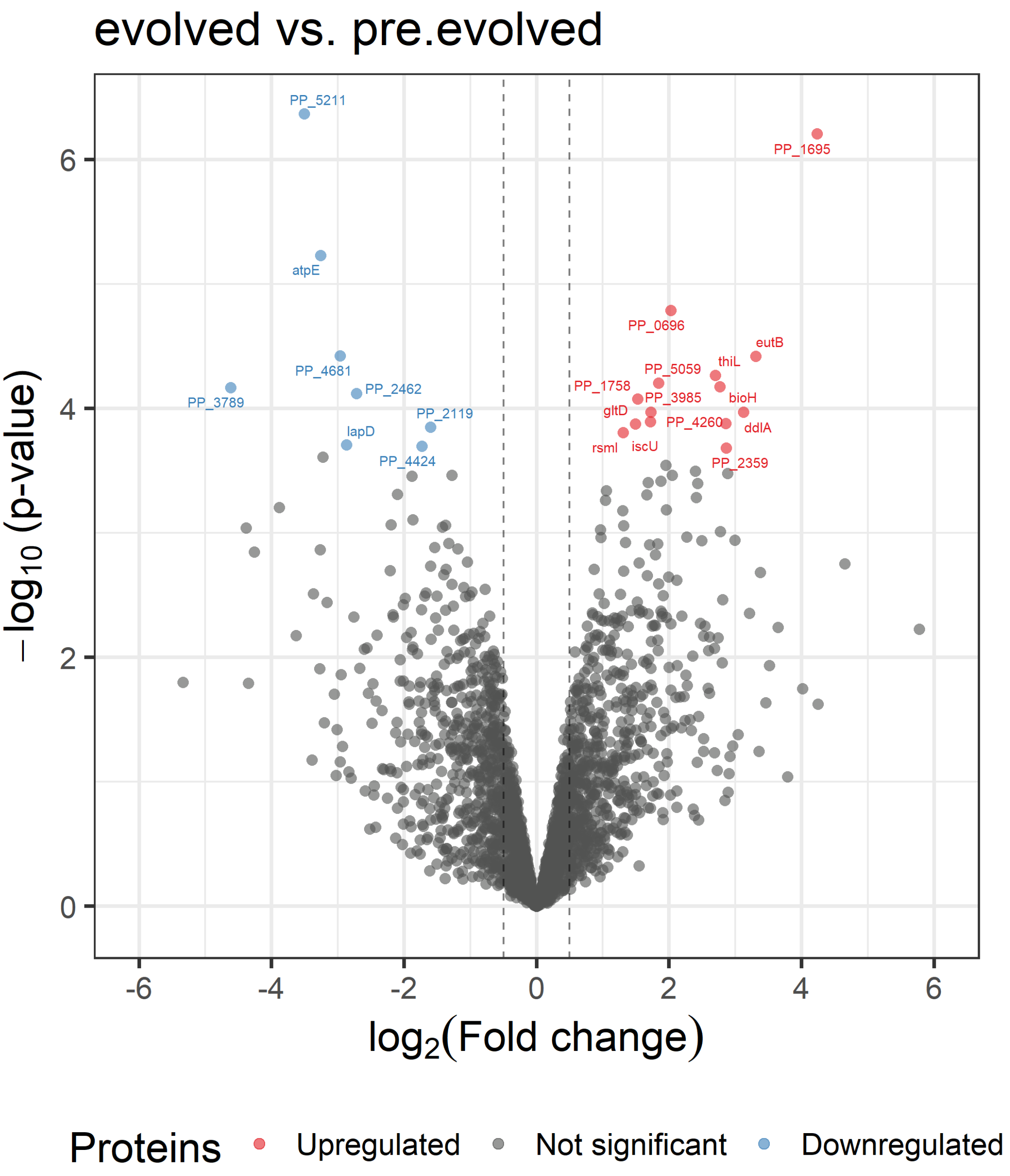

**Figure S9. Volcano plot of protein abundance data showing 28 differentially expressed proteins in evolved vs. pre-evolved SCA3$_{PK\text{-}Tn}$/pS438·*pta*$^{Ec}$.** Both strains (with four biological replicates) were grown in DBM medium supplemented with 30 mM glucose and 0.5 mM 3-*m*Bz, and harvested in the mid-exponential phase to measure the soluble protein content of biomass. Each point in the plot represents an individual protein. The horizontal intersection is set at an adjusted p-value of 0.05; the vertical intersections are placed for an absolute fold-change of ≥ 1.

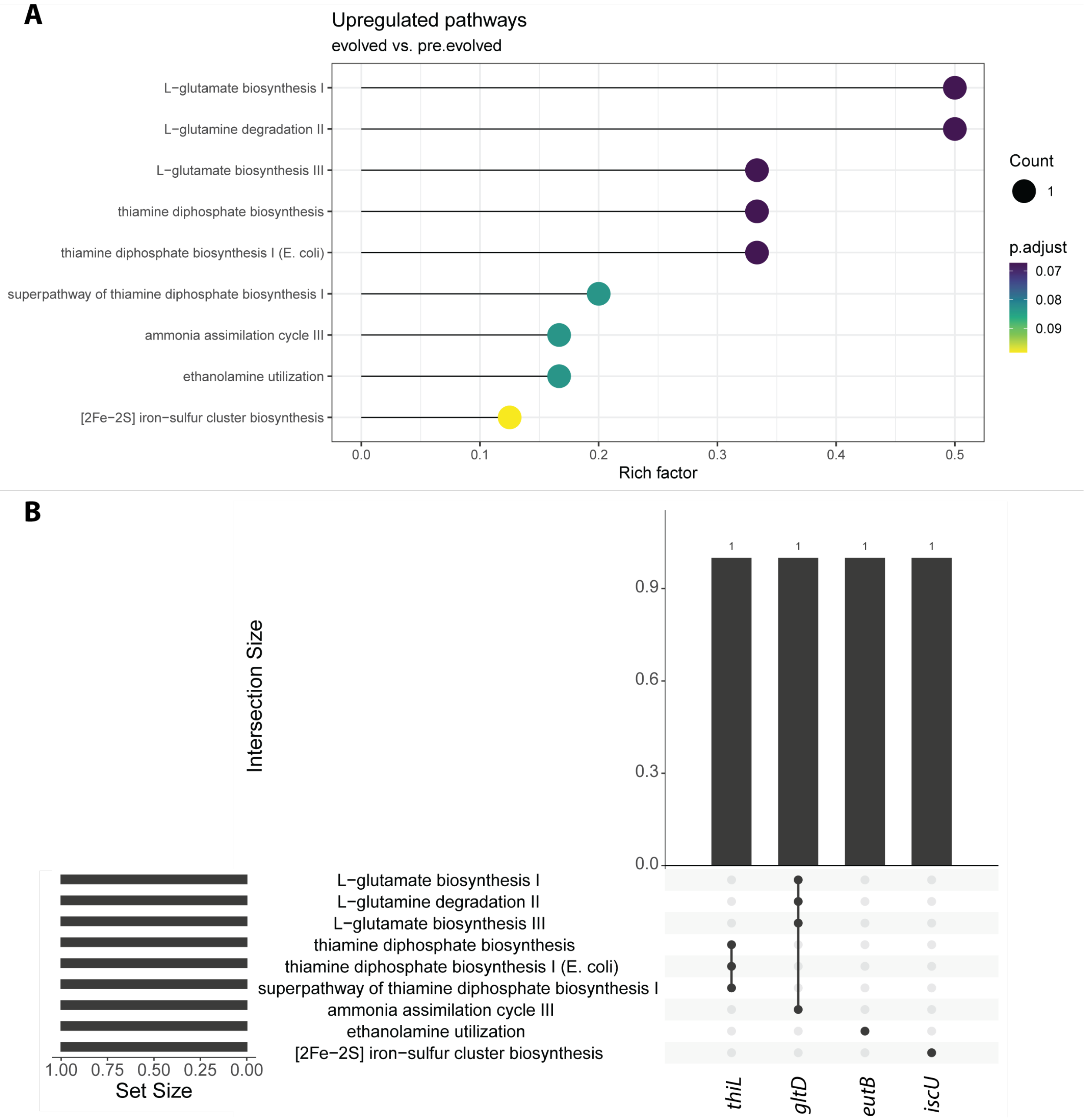

**Figure S10. Differentially enriched pathways in evolved vs. pre-evolved SCA3$_{PK\text{-}Tn}$/pS438·*pta*$^{Ec}$. (A)** Pathways from the BioCyc database (Karp et al., 2019) found to be significantly upregulated with a threshold for the adjusted p value (Benjamini-Hochberg method) of 0.1. No pathways were found to be significantly downregulated. The rich factor indicates what fraction of the total sum of proteins constituting a pathway was found to be differentially expressed. **(B)** Upset plot of proteins identified in the pathway enrichment analysis.

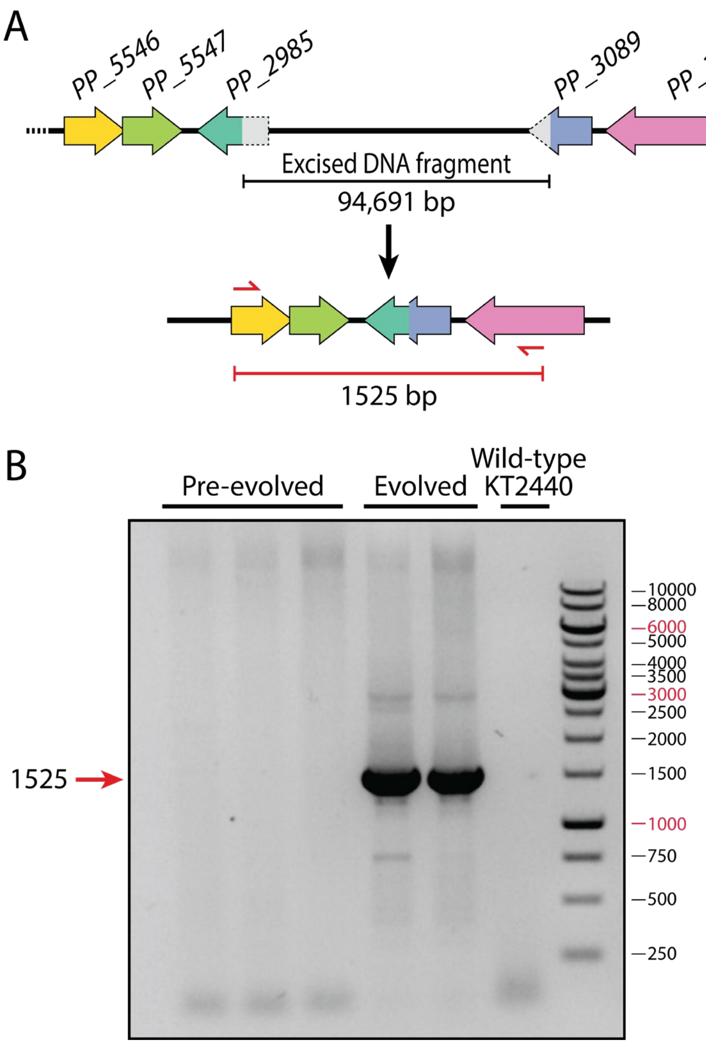

**Figure S11. Verification of DNA excision during the evolution of strain SCA3$_{PK\text{-}Tn}$/pS438·*pta*$^{Ec}$.** To verify the absence of a 94,691-bp stretch of DNA in the chromosome of 2 out of 4 evolved SCA3$_{PK\text{-}Tn}$/pS438·*pta*$^{Ec}$ clones, the respective region was amplified via PCR. **(A)** Re-arrangement of the chromosomal sequences during the excision. The binding regions of the primers used for PCR amplification are indicated in red. **(B)** Results of agarose gel electrophoresis with the amplified PCR fragments. The purified chromosomal DNA samples that had been analyzed by next-generation sequencing were used as the templates. A 1525-bp amplicon provides additional evidence for the absence of the DNA sequence identified as missing in the genome analyses.

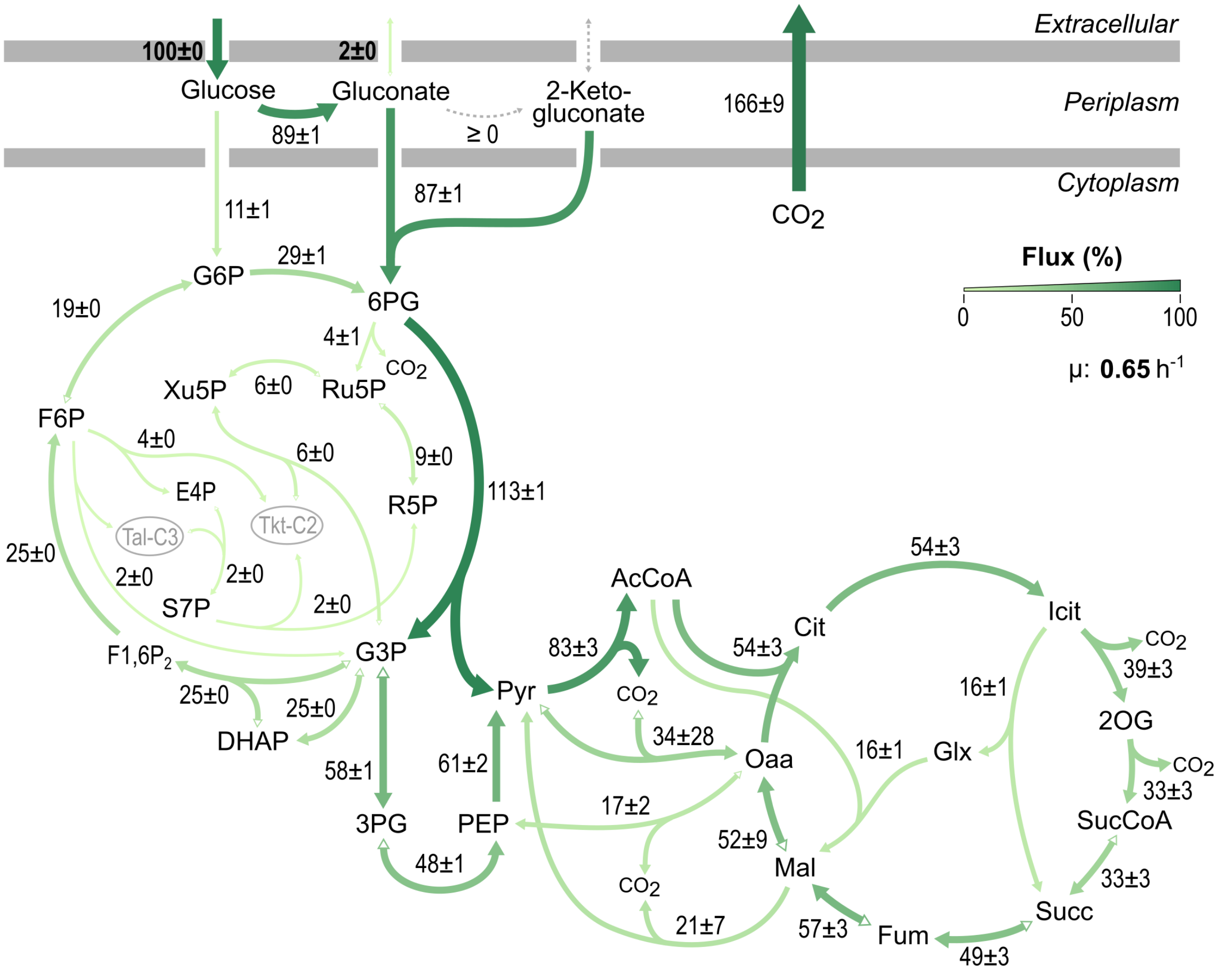

**Figure S12. In vivo carbon flux distribution of *P. putida* KT2440 grown on glucose.** Experimentally determined rates were set as fixed parameters (values in bold): glucose consumption rate, 10.45 ± 2.41 mmol $g_{CDW}^{-1}$ $h^{-1}$; gluconate secretion rate, 0.19 ± 0.07 mmol $g_{CDW}^{-1}$ $h^{-1}$; exponential growth rate, 0.65 ± 0.03 $h^{-1}$. No (net) production of 2KG and pyruvate were observed in this experiment. All fluxes were normalized to the specific glucose consumption rate. Flux values represent the mean ± standard deviations from three biological replicates.

### References

Huang, M., Oppermann, F. B., Steinbüchel, A., 1994. Molecular characterization of the Pseudomonas putida 2,3-butanediol catabolic pathway. FEMS Microbiol Lett. 124**,** 141-50.

Karp, P. D., Billington, R., Caspi, R., Fulcher, C. A., Latendresse, M., Kothari, A., Keseler, I. M., Krummenacker, M., Midford, P. E., Ong, Q., Ong, W. K., Paley, S. M., Subhraveti, P., 2019. The BioCyc collection of microbial genomes and metabolic pathways. Brief Bioinform. 20**,** 1085-1093.

Rühl, J., Hein, E.-M., Hayen, H., Schmid, A., Blank, L. M., 2012. The glycerophospholipid inventory of Pseudomonas putida is conserved between strains and enables growth condition-related alterations. Microb Biotechnol. 5**,** 45-58.

Volke, D. C., Turlin, J., Mol, V., Nikel, P. I., 2020. Physical decoupling of XylS/*Pm* regulatory elements and conditional proteolysis enable precise control of gene expression in *Pseudomonas putida*. Microbial Biotechnology. 13**,** 222-232.

Wirth, N. T., Nikel, P. I., 2021. Combinatorial pathway balancing provides biosynthetic access to 2-fluoro-*cis,cis*-muconate in engineered *Pseudomonas putida*. Chem Catalysis (accepted manuscript).
